# Comprehensive evaluation of deconvolution methods for human brain gene expression

**DOI:** 10.1101/2020.06.01.126839

**Authors:** Gavin J Sutton, Daniel Poppe, Rebecca K Simmons, Kieran Walsh, Urwah Nawaz, Ryan Lister, Johann A Gagnon-Bartsch, Irina Voineagu

**Author notes:** Corresponding author: Assoc. Prof. Irina Voineagu, School of Biotechnology and Biomolecular Sciences, University of New South Wales, Kensington, Sydney NSW 2052 Australia.

## Abstract

Gene expression measurements, similar to DNA methylation and proteomic measurements, are influenced by the cellular composition of the sample analysed. Deconvolution of bulk transcriptome data aims to estimate the cellular composition of a sample from its gene expression data, which in turn can be used to correct for composition differences across samples. Although a multitude of deconvolution methods have been developed, it is unclear whether their performance is consistent across tissues with different complexities of cellular composition. The human brain is unique in its transcriptomic diversity, expressing the highest diversity of alternative splicing isoforms and non-coding RNAs. It comprises a complex mixture of cell-types including transcriptionally similar sub-types of neurons, which undergo gene expression changes in response to neuronal activity. However, a comprehensive assessment of the accuracy of transcriptome deconvolution methods on human brain data is currently lacking.

Here we carry out the first comprehensive comparative evaluation of the accuracy of deconvolution methods for human brain transcriptome data, and assess the tissue-specificity of our key observations by comparison with transcriptome data from human pancreas and heart.

We evaluate 8 transcriptome deconvolution approaches, covering all main classes: 4 partial deconvolution methods, each applied with 9 different cell-type signatures, 2 enrichment methods, and 2 complete deconvolution methods. We test the accuracy of cell-type estimates using *in silico* mixtures of single-cell RNA-seq data, mixtures of neuronal and glial RNA, as well as nearly 2,000 human brain samples.

Our results bring several important insights into the performance of transcriptome deconvolution: **(a)** We find that cell-type signature data has a stronger impact on brain deconvolution accuracy than the choice of method. **(b)** We demonstrate that biological factors influencing brain cell-type signature data (*e.g.* brain region, *in vitro* cell culturing), have stronger effects on the deconvolution outcome than technical factors (*e.g.* RNA sequencing platform). **(c)** We find that partial deconvolution methods outperform complete deconvolution methods on human brain data. To facilitate wider implementation of correction for cellular composition, we develop a webtool that implements the best performing methods, and is available at https://voineagulab.shinyapps.io/BrainDeconvShiny/ .

## Introduction

Human tissues are mosaics of cell-types and subtypes, which are diverse in their functionalities and express distinct sets of genes. Consequently, gene expression measurements in any tissue sample are the result of two main factors: gene expression levels within constituent cell-types, and the relative abundance of these cell-types in the sample^1,2^. The relative abundance of cell-types (*i.e.* cellular composition) in turn depends on both biological^3–6^ and technical factors^7^.

To circumvent the confounding effect of cellular composition, gene expression measurements could in principle be carried out by experimentally isolating individual cell-types by laser capture micro-dissection^8,9^, cell sorting^10–12^, or single-cell RNA-seq (scRNA-seq)^13^. In practice, these approaches are limited in feasibility and cost effectiveness for human brain transcriptome studies that require large sample sizes (hundreds to thousands of samples), such as eQTL studies or gene expression studies aiming to identify low-magnitude changes in disease samples.

Several methods for *in silico* deconvolution have been developed to estimate the cellular composition of a tissue sample from its gene expression profile (reviewed in Avila Cobos *et al.*^1^). *In silico* deconvolution offers the opportunity to leverage scRNA-seq data to obtain deeper insights into bulk tissue transcriptomes generated through large-scale studies such as GTEx^14^, PsychEncode^15^, the Common Mind Consortium^16^, and BrainSpan^17^.

Deconvolution methods fall into two main categories: partial deconvolution (including enrichment approaches), and complete deconvolution (see Methods), and are conceptually similar for any tissue and any type of molecular data (transcriptome, methylome, proteome, *etc*.). However, the complexity of cellular composition, and the transcriptome similarity across cell-types varies widely across tissues. Most deconvolution methods have been developed for, or assessed on, blood/immune and tumour samples^18–20^, with limited assessment of their performance across tissues^2^. Therefore, an important outstanding question is whether transcriptome deconvolution methods perform equally well for any tissue.

We begin to address this question focussing on the human brain. The main biological factors that influence the cellular composition of brain samples (*e.g.* region, developmental stage, age^4,5^), and the technical factors involved (*e.g.* dissection protocol^7^) are distinct from those influencing cellular composition in blood. Furthermore, pure populations of cells from adult human brain are challenging to obtain, unlike blood or tumour cells. As a result, cell-type signature data are often obtained from a different brain region^21^, species^22,23^, and/or a different developmental stage^7^ than the bulk brain samples. Alternatively, cells cultured *in vitro* have been used^19^. Whether such choices influence the accuracy of brain cell-type composition estimates is unknown. In addition, gene expression changes in most psychiatric disorders, similarly to effect-sizes of common variants, are of low magnitude^24^. Therefore, to serve as useful co-variates, cell-type composition estimates need to discriminate small differences in cellular composition^3^. While a few studies have proposed methods focussed on brain tissue^5,7,23,25–27^, a comprehensive comparative assessment of the performance of deconvolution methods on brain transcriptome data is currently lacking.

Here, we performed a comprehensive evaluation of brain transcriptome deconvolution by assessing the performance of eight algorithms (four partial deconvolution, two enrichment, and two complete deconvolution methods). The partial deconvolution methods were each combined with nine types of cell-type signature data that differed in biological properties (cultured cells, immuno-purified cells, cross-species) or technical factors affecting RNA sequencing: single-nucleus RNA-seq (snRNA-seq), single-cell RNA-seq (scRNA-seq), bulk RNA-seq or CAGE-seq. These analyses were carried out on *in silico* mixtures of single-cell and single-nucleus transcriptomes, pure immuno-panned cell-types, mixtures of RNA extracted from pure populations of neurons and glial cells, as well as large-scale brain transcriptome data from the GTEx^14^ and PsychENCODE^15,28^ consortia.

Our results showed that cell-type signature data was the most important parameter for brain transcriptome deconvolution. The main biological factors influencing brain cell-type signature data, and consequently the deconvolution accuracy, were brain region and *in vitro* cell culturing. These factors had stronger effects on the deconvolution outcome than the sequencing platform (Illumina RNA-seq *vs.* Cap Analysis of Gene Expression (CAGE)). We also demonstrate that partial deconvolution methods, particularly CIBERSORT (implementing support vector regression), outperform complete deconvolution methods on human brain data. We also assessed the best approaches for correcting cell-type composition differences in differential expression analyses, and determined the magnitude of cell-type composition differences that can be effectively corrected for. Finally, we deconvolved large-scale gene expression data from GTEx and PsychENCODE consortia, and highlight the importance of assessing deconvolution accuracy on each brain dataset; for this purpose we developed a user-friendly web tool that implements the best performing methods identified in this benchmarking study (https://voineagulab.shinyapps.io/BrainDeconvShiny/).

## Results

To benchmark deconvolution methods for brain transcriptome data, we selected widely employed methods, and where possible included methods developed for brain data (Table 1). For partial deconvolution, we selected: CIBERSORT (*cib*), a highly-cited deconvolution method initially optimised for immune cell-types^18^; DeconRNASeq^29^ (*drs*) which implements the non-negative least squares approach employed by the PsychENCODE consortium^15^; MuSiC^30^(*music*), which is a single-cell-based deconvolution approach accounting for individual- and cell-specific expression variability in the signature; and dtangle^31^. For enrichment-based methods we selected xCell^19^, which has been recently applied by the GTEx consortium^32^, and BrainInABlender^7^ (*blender*), which was specifically developed for brain-derived data. Among complete deconvolution methods, we included Linseed^33^, which extends previous methods^34,35^, and the co-expression-based approach developed for brain data by Kelley et al.^5^ (*coex*).

### Assessment of deconvolution accuracy across methods

To assess deconvolution accuracy, we simulated data with known cell-type proportions using three adult human brain datasets: two snRNA-seq datasets, Velmeshev *et al.*^36^ (VL: 24,646 nuclei, 10X Chromium, Fig.1) and Hodge *et al.* from the Human Cell Atlas^37^ (CA: 11,314 nuclei, Smart-seq2, Supplementary Fig.1); as well as a scRNA-seq dataset Darmanis *et al.*^13^ (DM: 297 cells, Smart-seq, Supplementary Fig.2). For each dataset, 100 mixtures were simulated as the average expression of 500 randomly-sampled nuclei (VL, CA; Methods) or 100 randomly-sampled cells (DM; Methods). Cell-type signatures were generated as the average expression within each cell-type (Methods).

**Figure 1.**
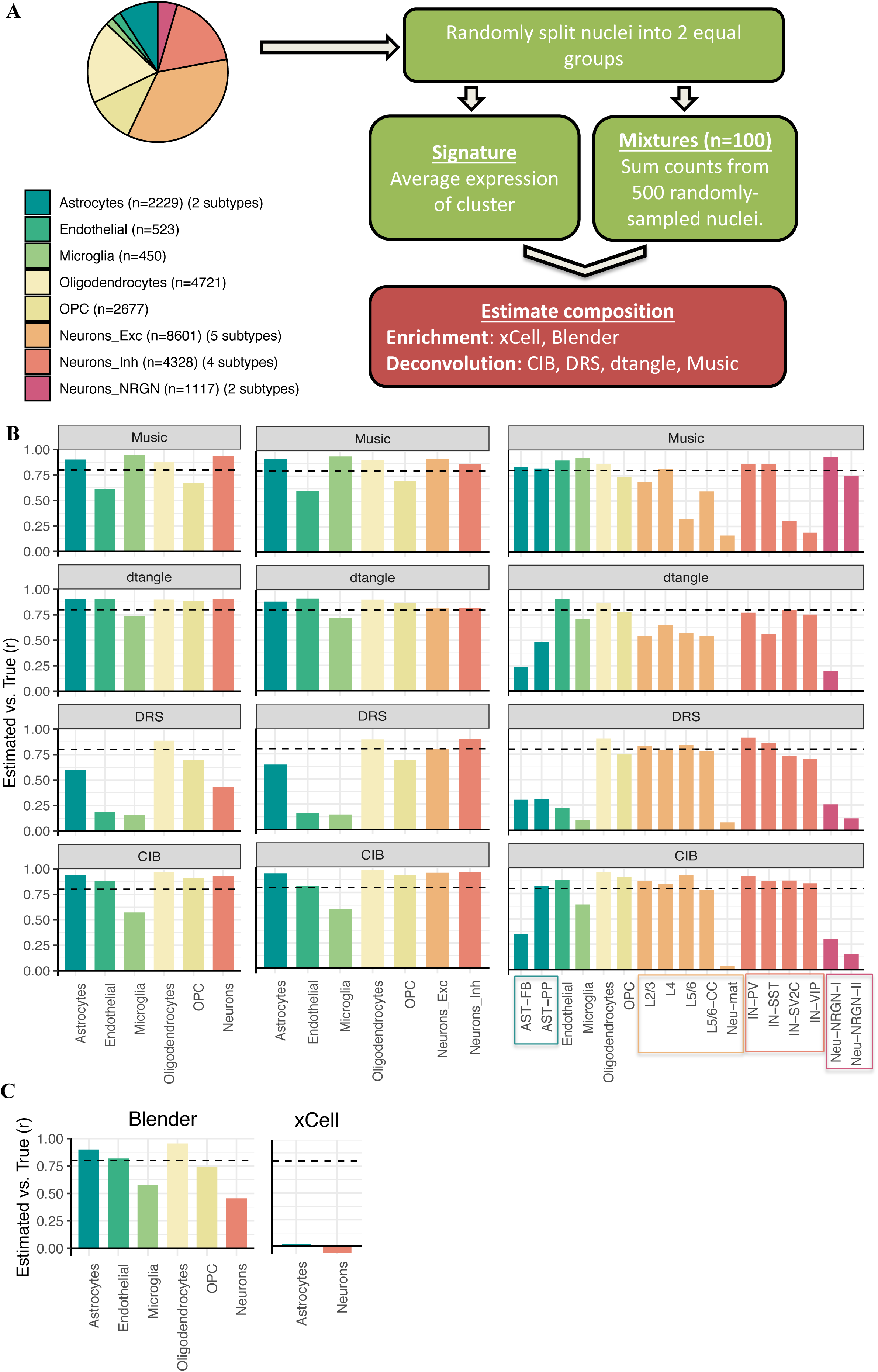
Deconvolution accuracy across methods. **A.** Simulation design. Single-nucleus RNA sequencing data was acquired from Velmeshev *et al*. and used to create *in silico* mixtures with known proportions. *Left:* Piechart displaying the composition of the dataset; n: number of cells per cell-type. For each cell type the number of sub-types is listed in between brackets. *Right:* analysis outline. *OPC*: oligodendrocyte precursor cells. *Neurons_Exc*, *Neurons*_Inh, and *Neurons*_NRGN: excitatory, inhibitory, and NRGN^+^ neurons, respectively. *DRS*: DeconRNASeq. *CIB*: *CIBERSORT*. *Blender*: BrainInABlender. **B.** Barplots of Pearson correlation coefficients (*r*) between true and estimated cell-type proportions in 100 in silico mixtures. *Left:* cells are grouped by major cell-types; *middle:* excitatory and inhibitory neuron subtypes are included in the signature; *right:* all cell-subtype labels are used in the signature. **C.** Barplots of Pearson correlation coefficients (*r*) between true proportion and cell-type enrichment scores in 100 *in silico* mixtures. *Dotted lines*: *r* = 0.8.

We first estimated cell-type proportions in these mixtures using *cib*, *drs*, *dtangle*, and *music*, and enrichment scores using *xCell* and *blender*, evaluating 6 major brain cell-types: neurons, astrocytes, oligodendrocytes, oligodendrocyte precursor cells (OPCs), microglia, and endothelia. Focusing on mixtures generated from the largest dataset (VL), we found that deconvolution accuracy was very high for *cib* (mean *r* across cell-types = 0.87), *music* (0.82), and *dtangle* (0.87), but lower for *drs* (0.50) (Fig.1B, left, and Supplementary Fig.3,4). For the two enrichment algorithms, *blender*’s accuracy was moderately high but inconsistent across cell-types, while *xCell* poorly estimated cell-type abundance (*r* = -0.06 and 0.02 for neurons and astrocytes, respectively); Fig.1C, and Supplementary Fig.4. These observations were replicated in both the CA- (Supplementary Fig.1,5,6) and DM-based simulations (Supplementary Fig.2), suggesting that (a) deconvolution of major brain cell-types is accurate across a range of partial deconvolution algorithms, with *cib* generally performing best and (b) enrichment methods are less accurate than partial deconvolution methods, with *xCell* showing particularly low accuracy.

We next assessed deconvolution accuracy on five *in vitro* RNA mixture samples of known composition (Supplementary Figure 7) and twenty-one RNA samples from pure populations of cells immuno-panned with cell-type-specific antibodies^38^; Supplementary Figure 8. In both cases, the deconvolution accuracy was very high when the signature was derived from the same source as the mixtures. For RNA mixtures the normalised mean absolute error was 0.035, 0.043, and 0.11 for *cib, drs,* and *dtangle,* respectively (Supplementary Figure 7). For RNA extracted from sorted cells, the immuno-panned cell-type was identified on average as 96.3%, 93.0%, and 92.6% abundant by *cib, drs,* and *dtangle,* respectively (Supplementary Figure 8).

#### • Deconvolution of cellular subtypes

We next explored how including cellular sub-types affected deconvolution accuracy for brain data. First, we used broad neuronal sub-types, *i.e.* excitatory and inhibitory neurons (Fig.1B middle, Supplementary Fig.3,9), and found that deconvolution accuracy was high (*r* > 0.8 for all algorithms), with *cib* performing best (*r* = 0.94 and 0.95 for excitatory and inhibitory, respectively). The accuracy for the other cell-types was largely unaffected by neurons being sub-classified (Fig.1B, middle, Supplementary Fig.3,9). This result was replicated in the CA-based simulations (Supplementary Fig.1,5,10).

When including all cell sub-types detected in the VL dataset (11 neuronal and 2 astrocyte sub-types), deconvolution with *cib* remained accurate (*r* > 0.8) for most cell populations (Fig.1B, right, Supplementary Fig.3,11,12). However, two main factors led to a reduction in accuracy for certain cell-types: low abundance of the cell-type (< 2%) and high collinearity (gene expression correlation with another cell-type *rho* > 0.95); Supplementary Fig.13,14. This observation was replicated in the CA dataset with most cell sub-types being accurately deconvolved (*r* > 0.8; Supplementary Fig.1,5,15,16) and collinearity being the main factor that led to reduced accuracy (Supplementary Fig.17,18).

#### • Deconvolution accuracy when a cell-type is missing from the reference signature data

We also explored how deconvolution was affected when a cell-type was missing from the signature. We removed one cell-type at a time from the VL-derived mixtures, and found that when an abundant cell-type was missing (Neurons, 87.4%), the deconvolution accuracy was substantially reduced (mean *r* was reduced from 0.85 to 0.41, and normalised mean absolute error increased from 0.33 to 10.3). However, when lowly-abundant cell-types were missing from the signature, the effect on deconvolution was minimal (Supplementary Figure 19). We then tested the effect of removing a sub-type of neurons, excitatory or inhibitory neurons, which are highly correlated in expression (*rho*=0.92). Deconvolution accuracy was reduced to a lesser extent then when all neurons were missing: *r* was reduced from 0.87 to 0.71 when excitatory neurons were missing, and from 0.86 to 0.76 when inhibitory neurons were missing (Supplementary Figure 19).

### The biological properties of cell-type signature data strongly influence deconvolution

We hypothesised that the brain cell-type signature data could have a major impact on the deconvolution outcome. This has been previously reported for deconvolution of blood transcriptomes^39^, and is supported by our observation that xCell performed significantly worse than the other deconvolution methods (noting that its signature data is built-in).

To investigate how the properties of the signature data influence the deconvolution outcome, we deconvolved the human brain snRNA-seq mixtures (VL) using cell-type signature data from several datasets (Methods; Fig.2) with different sequencing methods and various sources of brain tissue: human brain snRNA-seq (CA^37^, NG^40^, LK^41^); scRNA-seq from the human (DM^13^) or mouse (TS^42^) brain; bulk RNA-seq of immuno-panned cells from the human (IP^38^) or mouse brain (MM^43^); or CAGE-seq from cultured human brain cells (F5^44^).

**Figure 2.**
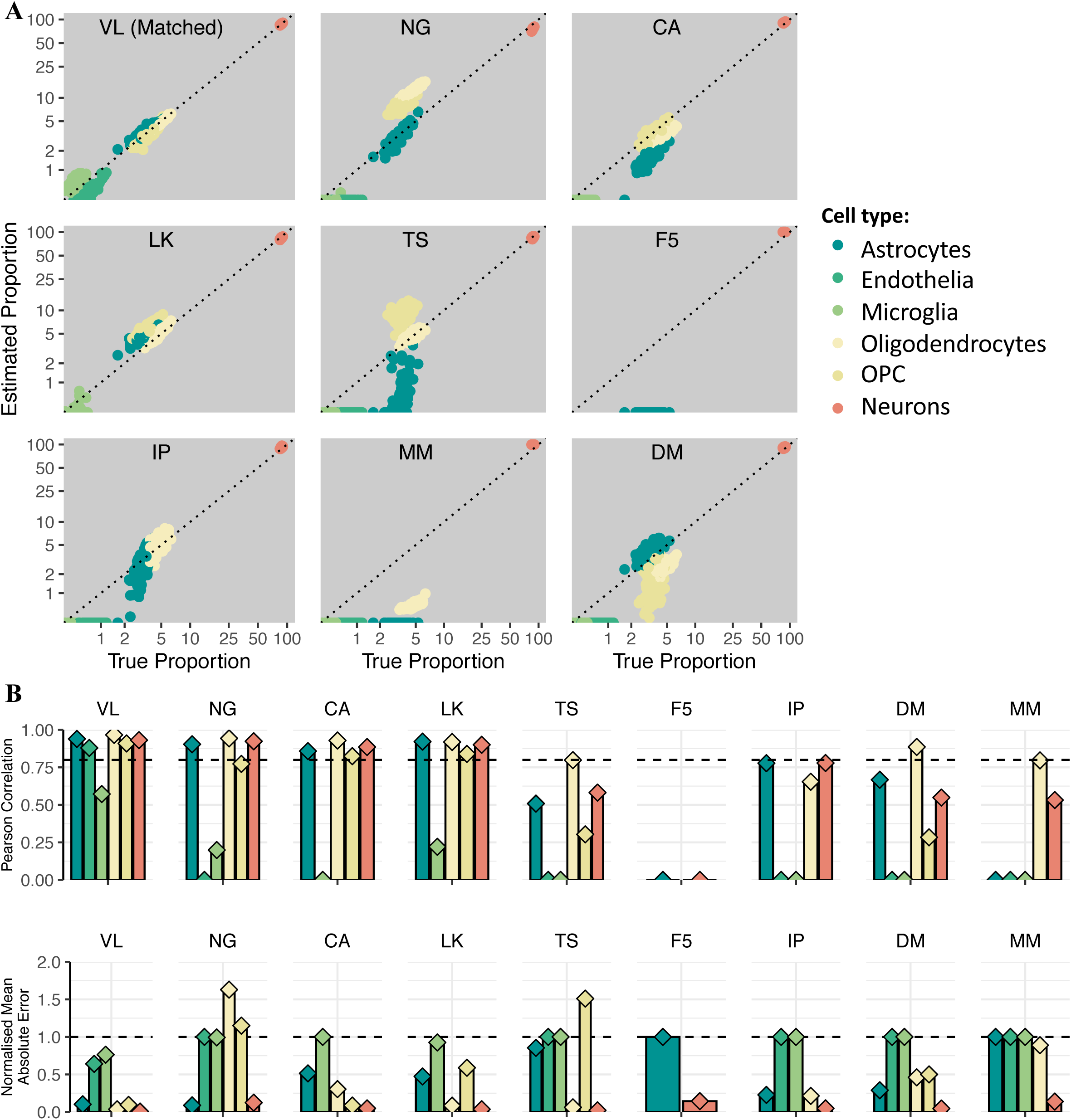
Effect of signature choice on deconvolution accuracy. **A.** Scatterplots of true and CIBERSORT-estimated proportions in VL *in silico* mixtures, for nine different signatures. *Matched:* the signature and mixture were derived from the same dataset. *VL*: Velmeshev. *NG*: Nagy. *CA*: Human Cell Atlas. *LK*: Lake. *TS*: Tasic. *Dotted line:* y=x. **B.** Barplots of normalized mean absolute error (*nmae*) for all cell-types and signatures presented in A. *Ast*: astrocytes. *End*: endothelia. *Mic*: microglia. *Oli*: oligodendrocytes. *OPC*: oligodendrocyte precursor cells. *Exc*: excitatory neurons. *Inh*: inhibitory neurons. *Dotted line:* nmae = 1. **C.** Barplots of Pearson correlation (*r*) for all cell-types and signatures presented in A. *Dotted line*: *r* = 0.8.

We found that the choice of cell-type signature data strongly affected the deconvolution accuracy. Using data from cultured brain cells (F5) dramatically reduced the accuracy (Fig.2A-B). This likely explains the poor performance of xCell, which has F5 as the main built-in signature. Using signature data from the mouse brain (TS, MM) also reduced the deconvolution accuracy (Fig.2A-B). These observations were consistent across deconvolution algorithms (Supplementary Fig. 20) and were replicated when deconvolving *in silico* mixtures based on the CA and DM data (Supplementary Fig.21,22), as well as with deconvolution of broad neuronal subtypes (Supplementary Fig.23,24). Conversely, when deconvolving RNA mixtures of known composition from cultured cells (Methods), using the cultured-cell F5 signature data performed the best despite the difference in sequencing technology (CAGE-seq *vs.* RNA-seq; Supplementary Fig.7).

Overall, these data demonstrate that the biological properties of the cell-type signature data strongly impact the deconvolution accuracy, having a more pronounced effect than the sequencing methods, and highlight *in vitro* culturing of brain cells as an important biological factor.

#### • The effect of compartment specific genes on deconvolution accuracy

Since most single-cell data from the adult human brain are generated using single-nucleus RNA-seq, while bulk RNA-seq is based on total RNA, we next investigated whether compartment-specific genes (*i.e.* those either enriched or depleted from the nucleus) influence the outcome of deconvolution. For this purpose, we generated paired bulk RNA-seq and nuclear RNA-seq from five frozen brain tissue samples (Methods), as well as snRNA-seq from the same brain samples. We identified compartment-specific genes as those differentially-expressed between the nuclear and total bulk RNA-seq (FDR < 0.05, |FC| > 1.3; Supplementary Table 1). We then carried out several deconvolution analyses with and without filtering-out the compartment-specific genes.

Firstly, we deconvolved the twenty-one bulk RNA-seq samples from sorted brain cells^38^, where true cell-type composition is known (i.e. each sample is expected to be a nearly-pure cell-type, with some experimental variability of the sorting efficiency). We deconvolved these data with either the identical cell-type signature (derived from the sorted dataset; IP), an scRNA-seq signature (DM), and four snRNA-seq signatures (VL, CA, NG, and LK) ; Supplementary Table 2. When using the IP signature, the immuno-panned cell-type was estimated as > 80% abundant in all samples. Thus we assessed the proportion of samples in which the sorted cell-type was correctly identified (i.e estimated proportion > 80%) using the scRNA-seq and snRNA-seq signatures, with or without filtering out compartment-specific genes. We found that the snRNA-seq signatures performed well even prior to filtering out compartment-specific genes, correctly identifying the sorted cell-type in an average of 86% of samples (71%-95%). As expected, the single-cell-based signature (Supplementary Fig.25) performed somewhat better, identifying the sorted cell-type in an average of 90% of samples. Removing compartment-specific genes further improved the outcome for snRNA-seq signatures: the sorted cell-type was correctly identified in an average of 88% of samples (86%-95%); Supplementary Fig.25, eliminating the difference between the scRNA-seq and snRNA-seq signatures.

To increase the complexity of the deconvolution task, we asked how accurately the five whole-tissue samples were deconvolved when using snRNA-seq data from the same individuals, as compared to a whole-cell based signature (Supplementary Fig.25). In this case, if compartment-specific genes were not removed from the cell-type signature, the correlation between cell-type proportions estimated using the snRNA-seq signature and the whole-cell signature was modest (r=0.27). However, the correlation improved substantially by filtering out compartment-specific genes (r=0.98) suggesting that this filtering approach should be considered when using snRNA-seq-based cell-type signatures.

### Reference-free complete deconvolution methods are less effective on brain gene expression data than partial deconvolution methods

Since we observed a strong effect of the choice of reference signature data on the brain deconvolution outcome, and recent studies have proposed reference-free approaches to cell-type composition^33,34,45^, we assessed the performance of two such methods on brain data. Linseed, a complete deconvolution algorithm^33^, proposes to identify cell-type specific genes by representing the expression vector of each gene as a point in N-dimensional space (where N is the number of samples). It also proposes using singular-value-decomposition (SVD) to determine the number of cell-types from the mixture data. An alternative approach, *coex*^46^, employs co-expression networks to identify modules of co-expressed genes enriched for specific cell-type markers, and then uses the module eigengene values as cell-type enrichment scores^5^.

When applying Linseed to *in silico* mixtures generated by random sampling from the three benchmarking datasets (VL, CA, DM), we found that the SVD approach did not correctly identify the number of cell-types in the mixture (Methods; Supplementary Fig.26). With the correct number of cell-types specified, Linseed performed less accurately than the partial deconvolution methods, with *r* > 0.8 achieved for only two cell-types for VL and CA, and none of the cell-types for DM (Fig.3B, Supplementary Fig.27). On the RNA mixtures however, Linseed performed very accurately (*r*=1; Supplementary Fig.28). Since Linseed relies on the detection of genes represented by points with “extreme” positions in the k-1 dimensional simplex, we hypothesized that the difference in its performance between the datasets likely results from the wider distribution of cell-type proportions in the RNA mixtures (neuronal proportions: 0-100%), than in the mixtures generated by random sampling from real brain datasets. To test this hypothesis, we generated mixtures with a broad range of cell-type abundances using VL and CA (Methods; Fig.3A,C). The performance of Linseed improved markedly on both datasets using these controlled *in silico* mixtures (Fig.3D, Supplementary Fig.29,30), with the SVD approach also better identifying the number of cell-types in the mixture (Methods).

**Figure 3.**
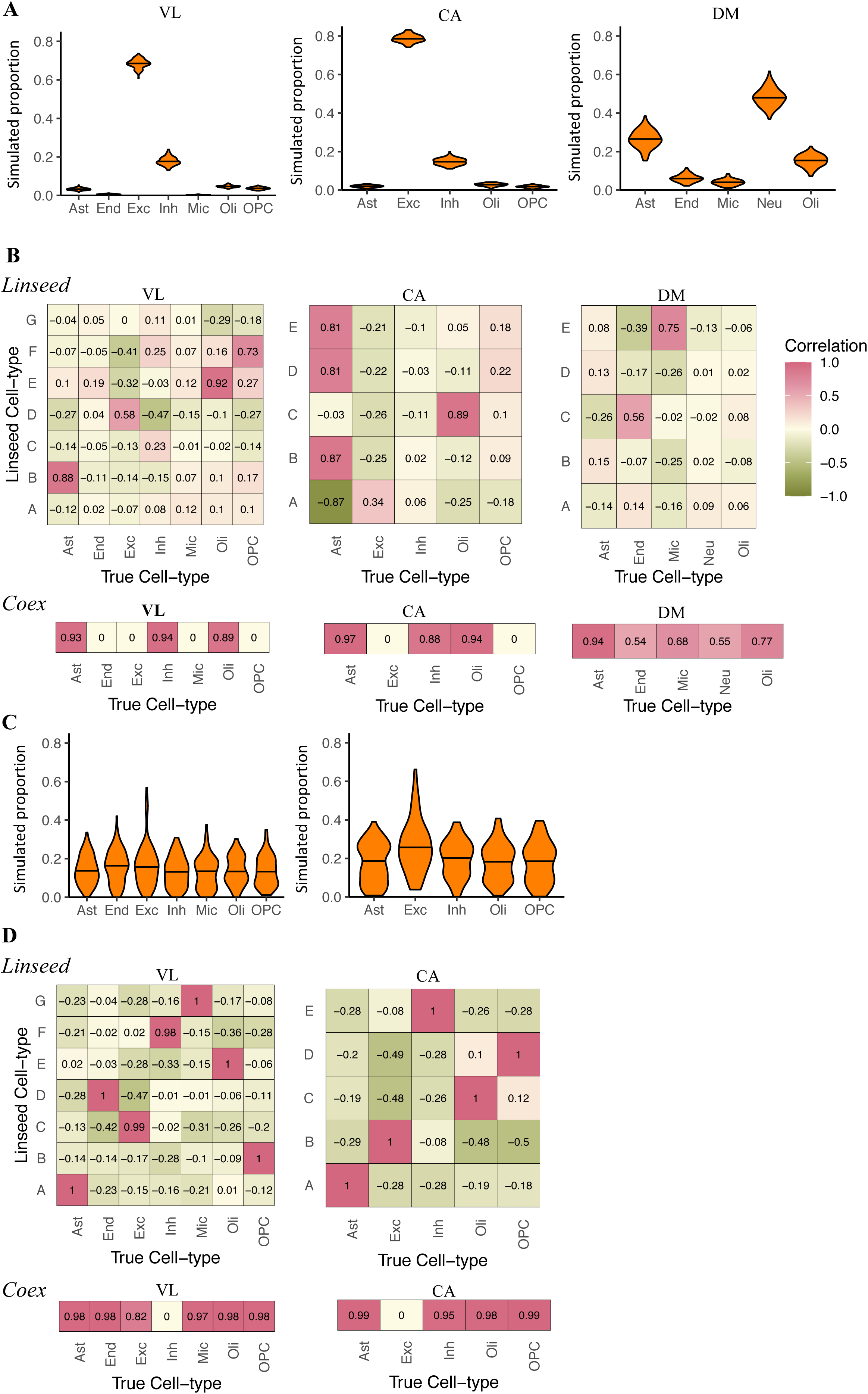
Reference-free deconvolution. **A.** Violin plots of the distribution of true cell-type proportions in VL, CA, and DM *in silico* random simulations (left, middle, and right respectively). *Black horizontal bar*: median. *Ast*: astrocytes. *End*: endothelia. *Exc*: excitatory neurons. *Inh*: inhibitory neurons. *Neu:* neurons. *Oli*: oligodendrocytes. *OPC*: oligodendrocyte precursors. **B.** Heatmaps of Pearson correlation coefficients between estimated and true cell-type proportions for random simulations based on VL, CA, and DM. *Top:* Linseed; y-axis: Cell-types defined by Linseed; x-axis: true cell-type in simulated data. *Bottom:* Coex; for each cell type the true vs. estimated correlation coefficient is displayed for the coexpression module assigned to that cell-type based on marker enrichment (Methods); zero values represent cases where no coexpression module was assigned to the corresponding cell-type. **C.** Violin plots of the distribution of cell-type abundances in simulations with wide cell-type ranges based on VL (left) and CA (right). **D.** Heatmaps of Pearson correlation coefficients between estimated and true cell-type proportions for simulations with wide cell-type ranges displayed in C, based on VL and CA. *Top:* Linseed. *Bottom:* Coex.

We found that *coex* also performed significantly less accurately than the partial deconvolution methods on the randomly-sampled mixtures (Fig.3B, Supplementary Fig.27). Since the co-expression network approach also relies on gene expression co-variation driven by differences in cell-type proportions, its performance improved on simulations with a wider range of cell-type proportions, but did not achieve accurate deconvolution for all cell-types (Fig.3D, Supplementary Fig.29,30).

These data suggest that complete deconvolution methods are less effective than partial deconvolution methods, particularly since the performance of these algorithms is related to the variance in cellular composition of the dataset, which is not known *a priori*.

### Assessment of the interplay between cell-type composition and differential gene (DE) expression analyses

We next investigated how cellular composition influences DE analyses, in particular: (i) how much should cell-type composition differ between two groups of brain samples to lead to false positive results in DE analyses, and (ii) what is the best approach for correcting cell-type composition differences in DE analyses?

We used the CA dataset^37^ (Smart-seq2, high coverage per gene) to generate simulated data for two-group DE analyses. Each dataset contained two groups of 50 samples (group A and group B). The proportion of one of the cell-types (excitatory neurons) was simulated as either higher or lower in group B than in group A by 0% to 40% (Methods; Fig.4). We then carried out DE comparing group B to group A, using either a linear model (LM) or a generalised LM as implemented in DESeq2^47^ with and without correction for cellular composition. False-positives driven by cellular composition were defined as genes differentially expressed at a false discovery rate (FDR) < 0.05 (Methods).

**Figure 4.**
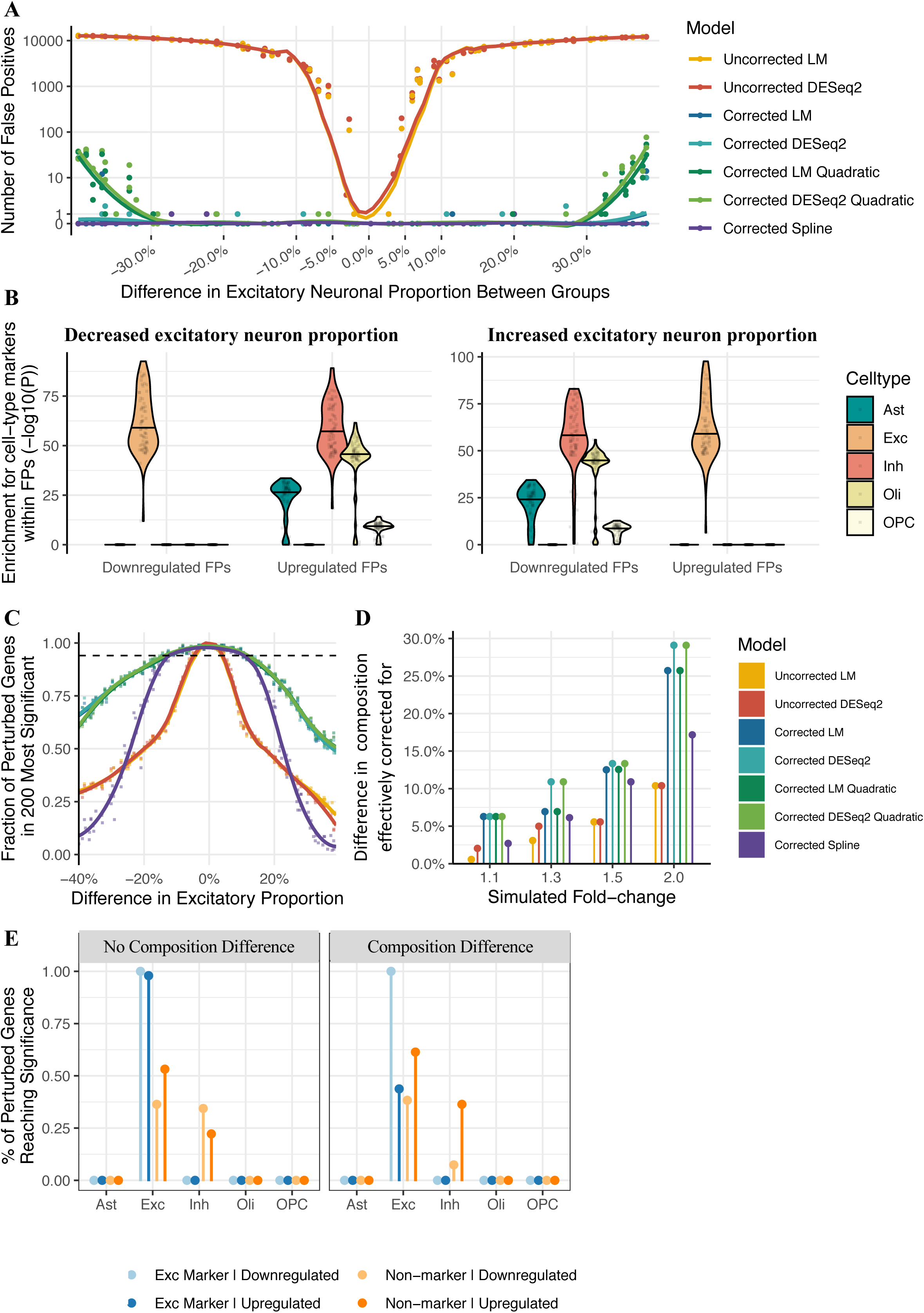
Effect of brain cell-type composition on differential expression (DE) analyses. **A.** Scatterplot of the number of false positive genes versus the simulated difference in excitatory neuron proportion between two groups of 50 samples. Each point represents a different simulated dataset. DE was assessed with either a linear model (LM) or DESeq2, with or without correction for composition, *Coloured lines:* local regression line. **B.** Cell-type marker enrichment within false positive genes. Each point represents a single simulated dataset. *Y-axis*: enrichment p-value (one-sided Fisher test); Methods. *FPs*: false positive genes. **C.** Scatterplot of the discriminatory power, *i.e.* fraction of the 200 perturbed genes in the top 200 most significantly differentially expressed genes (y-axis) versus simulated difference in excitatory neuron proportion between sample groups (x-axis) for simulated 1.5-fold expression difference. *Coloured lines*: local regression line. *Dotted line:* expected discriminatory power, *i.e.* 0.95 times the discriminatory power in the absence of cell-type composition differences between groups. **D.** Model robustness to cell-type composition differences across a range of fold-changes, quantified as the smallest composition change where discriminatory power fell below its expected value. **E.** Cell-type specific DE analysis using CIBERSORTx for simulated mixtures with 1.5-fold difference in gene expression between two groups, without (*Left*) or with (*Right*) a superimposed cell-type composition difference between the two groups. Gene expression was perturbed in excitatory neurons for 100 non-marker genes and 100 excitatory neuron marker genes. The cell type composition of simulated data is shown in Supplementary Fig. 32. Y-axis: fraction of perturbed genes significantly DE at FDR < 0.05; DE was carried out using a linear model (Methods).

We found that without correction, differences in cellular composition of less than 5% between the sample groups led to fewer than 10 false-positive DE genes. However, above 5% the number of false-positive genes increased steeply with the difference in cellular composition, reaching > 10,000 at a 20% difference in cellular composition (Fig.4A). Inclusion of excitatory neuron proportions as a covariate in the LM effectively eliminated false-positive genes (Fig.4A). We didn’t observe any additional benefit when using a spline matrix as covariate, while quadratic regression was less effective than linear regression at larger composition confounds (Fig. 4A). We also found that including cellular composition estimates in DESeq2 was similarly effective at eliminating false-positives (Fig.4A). As expected, markers of excitatory neurons were enriched among downregulated genes when the proportion of this cell-type was reduced in the test group, but enriched among upregulated genes when the proportion was increased (Fig.4B).

We next investigated the more challenging case where there are true differences in gene expression between the two groups, in addition to differences in cell-type composition. To this end, we simulated data with differences in cell-type composition as above, while also introducing gene expression changes in a set of 200 genes, of which 100 are markers of excitatory neurons and 100 are non-marker genes (Methods). Several sets of simulations were generated with a mean expression difference between groups of 1.1-, 1.3-, 1.5- or 2-fold.

To quantify how effectively cellular composition was corrected for, we calculated discriminatory power as the fraction of the 200 perturbed genes that were in the top 200 most significant DE genes. This measure rewards true-positives while penalising false-positives. We found that without correction, the discriminatory power decreased with the magnitude of cell-type composition difference between the two groups (Fig.4C; uncorrected). Correction for cell-type composition was effective at restoring discriminatory power for gene expression differences of 1.5 fold when composition differences were up to 12.5% (Fig.4D; corrected). As expected, for expression differences of lower magnitude (1.1 and 1.3) the effective correction range was narrower (6.3% and 6.9% respectively), while for stronger expression differences (2-fold) the effective correction range was wider (25.7%); Fig.4D, Supplementary Fig.31. All correction approaches performed similarly in this analysis, with the exception of spline regression which we found to be less effective (Fig.4C,D, Supplementary Fig.31).

We also investigated whether the cell-type where differential expression occurs can be uncovered through deconvolution. To this end, we used CIBERSORTx^48^, which takes a bulk mixture and estimates gene expression values in each cell-type present in the signature data; these cell-type-specific expression values can then be used to carry out cell-type specific DE analyses. We thus simulated data with 1.5-fold change in expression specific to a given cell-type, with or without superimposed differences in cell-type composition between the two groups (Supplementary Fig.32), and tested whether genes were identified as DE in the correct cell-type. As above, the 1.5-fold expression difference was simulated for 200 genes, of which 100 are markers of the perturbed cell-type and 100 are non-marker genes.

The expression difference was first simulated in excitatory neurons. In the absence of confounding cell-type composition differences between the two groups, more than 95% of the perturbed excitatory marker genes were detected as DE in the correct cell-type (excitatory neurons), while for the non-marker genes ∼45% were detected as DE in excitatory neurons and another ∼30% were incorrectly detected as DE in inhibitory neurons; Fig. 4E. The false-positive rate (*i.e.* the fraction of non-perturbed genes detected as DE) was 0% (Supplementary Fig.32).

When a composition difference was superimposed (∼10% increase in excitatory neurons; Methods), the true-positive rate was unchanged except for upregulated marker genes, where it was reduced, likely due to the fact that the composition change and the expression change were confounded (both variables were higher in group B *vs.* group A). The false-positive rate was less than 12% in all cell-types, thus drastically reduced relative to no correction for cell-type composition (32%), but higher than when correcting for composition differences in a standard linear model (0%). Similar results were observed when the gene expression change was modelled in inhibitory neurons (Supplementary Fig.32).

Overall, these results suggest that using cell-type specific gene expression for DE analyses is effective at detecting DE genes in the right cell-type when the gene expression and composition changes are not confounded, but this comes at the expense of a moderate increase in false-positives.

### Cell-type composition estimates in large-scale human brain transcriptome data

We next evaluated the performance of brain gene expression deconvolution on large-scale datasets, focussing on a dataset of control individuals (GTEx^14^, n=1,671 samples; Methods), and a dataset of autism spectrum disorder (ASD) cases and controls (PsychENCODE; Parikshak *et al.*^28^, n=251 samples; Methods). The GTEx data included samples from cerebellum (n=309), cerebral cortex (n=408), subcortical regions (n=863) and spinal cord (n=91); the Parikshak *et al.* dataset included samples from cerebellum (n=84) and cerebral cortex (n=167).

We assessed all combinations of 4 partial deconvolution methods and 9 cell-type signatures, the two enrichment methods, and *coex* as a complete deconvolution method (Supplementary Table 3). We also generated an additional signature (MultiBrain) by merging CA, IP, DM, NG, and VL signatures derived from cortex, reasoning that this approach will average-out inter-individual and technical differences, as previously proposed^39^.

The accuracy of composition estimates was first evaluated on cortical samples using goodness-of-fit, *i.e.* the Pearson correlation between measured gene expression and reconstructed gene expression values (Methods). Consistent with the results on simulated data, we found that cell-type signature data had a stronger impact on accuracy than the choice of algorithm (Supplementary Fig.33). Although there was some variation between the two datasets, the CA and MultiBrain signatures performed consistently well, while the cultured-cell*-*derived F5 and the single-nucleus LK signatures performed worst (Fig.5A,B). Cerebellar samples showed lower goodness of fit than cortical samples in both datasets (Supplementary Fig.34,35), consistent with the fact that all cell-type signatures were derived from cerebral cortex. When the biological and technical differences between the bulk data and cell-type signatures are eliminated, as is the case of our *in silico* mixtures of single-cell data, goodness-of-fit averaged ∼0.95 (Supplementary Fig.36). Further details on the deconvolution results of the GTEx and Parikshak *et al.* data (correlation and absolute values of cell-type abundance estimates) are included in Supplementary Note.

**Figure 5.**
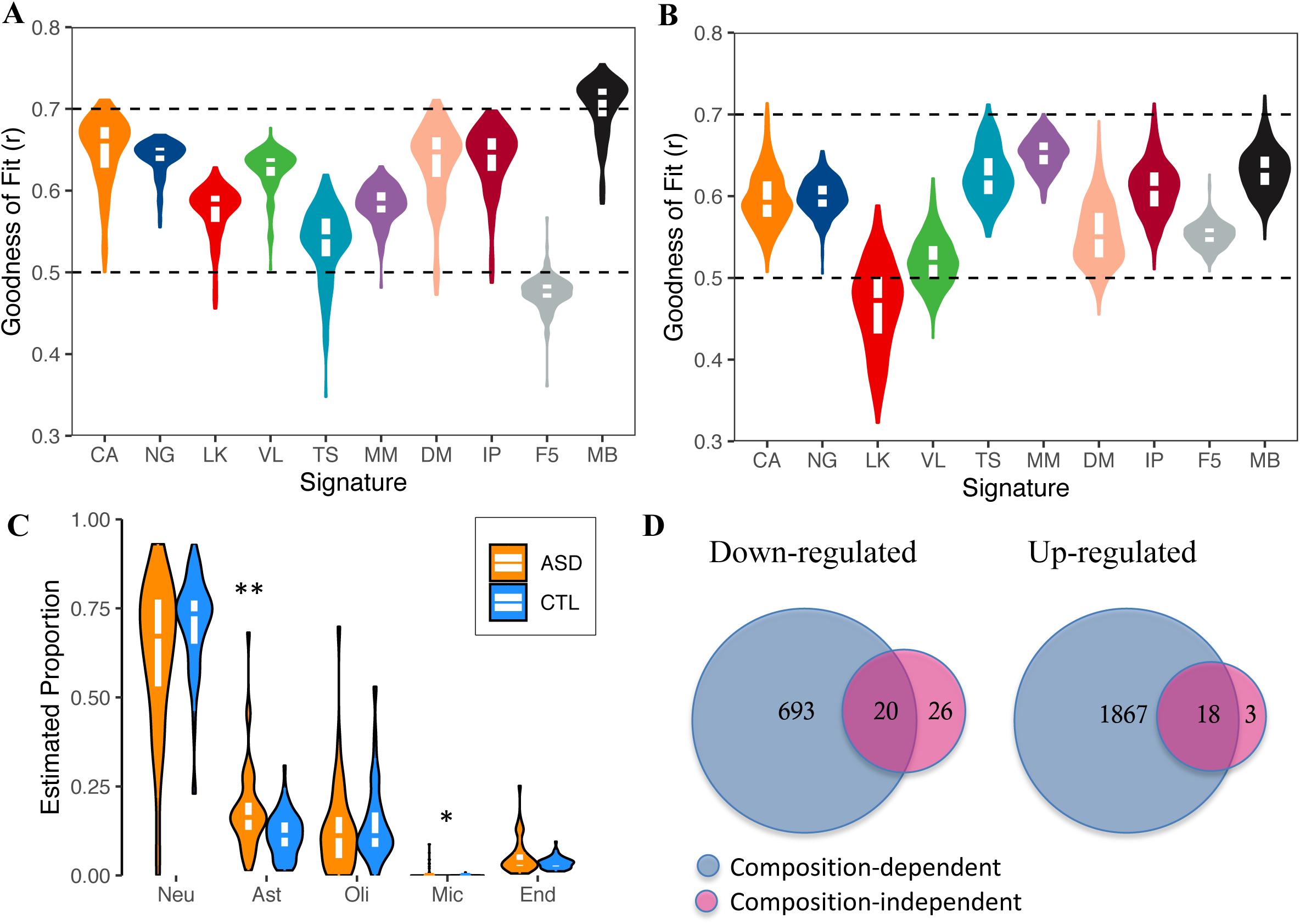
Cell-type composition estimates in large-scale human brain transcriptome data. **A-B.** Goodness of Fit across signatures in cortex samples from the GTEx consortium (A) and Parikshak *et al.* (B). Deconvolution was performed using CIBERSORT. The top, middle, and bottom of the white internal boxes mark the 75^th^, 50^th^, and 25^th^ percentiles, respectively. **C.** Composition estimates in ASD (n=43) and Control (n=63) cortical samples. Deconvolution was performed using CIBERSORT and the MultiBrain signature. *ASD*: autism spectrum disorder. *CTL*: control. *: p<0.01. ***: p<0.001. **D.** Venn diagrams of the overlap between composition-dependent and composition-dependent DE genes between ASD and CTL samples.

While the important role of signature data has been previously reported^39^, here we uncovered the effect of novel biological factors that affect deconvolution accuracy, in particular *in-vitro* culturing and brain region. Since *in-vitro* culturing may be relevant to other tissues as well, we investigated whether using cultured-cell-derived or tissue-derived signature data influences the accuracy of deconvolution for two additional tissues in GTEx: pancreas and heart. We found that deconvolution accuracy for left heart ventricle and arterial appendage samples was significantly reduced when using signature data from cultured cells, while the deconvolution accuracy for pancreas data was mildly reduced (Supplementary Fig.37). These data suggest that the influence of biological factors on cell-type signature data vary across tissues, indicating that tissue-specific benchmarking of deconvolution approaches is warranted.

Finally, we applied the results of the cell-type composition analyses to get further insights into genes differentially expressed in brain tissue samples from ASD cases^28^. Cell-type proportion estimates (CIB/*MultiBrain*), showed significantly higher astrocyte proportions in ASD cortex samples compared to controls (difference in means: 7.2%, p=0.0002, Wilcoxon rank sum test; Fig.5C). This result recapitulates recent single-nucleus data from ASD brain validated by immunohistochemistry^36^, which showed higher proportion of astrocytes in ASD cortex samples. There were also significantly higher proportions of microglia (0.7%, p=0.003; Wilcoxon rank sum test), although the overall abundance of microglia was low.

We next carried out differential expression (DE) analyses either without correction for cellular composition (composition-dependent; CD) or including cell-type proportion estimates from CIB/*MultiBrain* in the model (composition-independent; CI); Methods. Astrocyte, oligodendrocyte, and microglial proportions were included as covariates. CD analyses identified 713 down- and 1885 up-regulated genes (Fig.5D). In contrast, when correcting for composition estimates in our CI analyses, we identified only 46 down- and 21 up-regulated genes (Fig.5D). Of these, 20 down- and 18 up-regulated genes overlapped between CI and CD analyses (Fig.5D). Thus, 26 down-regulated and 3 up-regulated genes were uncovered by the CI analysis, and have not previously been reported as DE in ASD^28^. Conversely, 693 down-regulated and 1867 up-regulated genes were identified in the CD analysis only, and thus likely reflect differences in cellular composition between the ASD and control samples, rather than gene expression dysregulation (Supplementary Table 4). The CD upregulated genes were enriched for immune and inflammatory genes (Supplementary Table 4) as well as astrocyte and microglial markers (*p*=2.5×10^-11^ and 4.7×10^-36^, respectively), consistent with their higher proportions in ASD samples. Notably, one of the top up-regulated novel genes, *CXXC4,* which encodes a protein involved in Wnt signalling, has also been identified as upregulated in ASD CTX layer 4 neurons by single-nucleus RNA-seq^36^. In addition, *CXXC4* was identified as the top associated gene in a GWAS meta-analysis of schizophrenia and ASD^49^. These data indicate that correction for cellular composition can identify novel, disease-relevant gene expression changes.

## Discussion

Here, we began to address the question of tissue-specificity in transcriptome deconvolution, by carrying out a comprehensive benchmarking of deconvolution methods on brain transcriptome data. We assessed eight deconvolution methods, as well as multiple parameters of deconvolution: the biological and technical properties of the cell-type signature data; the effect of deconvolving brain cell sub-types; the effect of missing cell-types in the signature data; and the effect of nuclear-enriched or depleted transcripts on deconvolution using snRNA-seq signatures. We also investigated how effectively cell-type composition differences can be corrected in DE analyses.

It has previously been shown that cell-type signature data has a strong effect on deconvolution accuracy^39,50^. In blood, the microarray platform was the main factor driving differences between cell-type signature datasets^39^. For deconvolution of solid tumours, accurate estimation of immune cell-type composition required tumour-derived cell-type signatures, rather than blood-derived signatures^50^. For brain transcriptomes, we found that cell-type signature data had a stronger impact than the choice of method in all cases studied: simulated single-cell mixture data, RNA mixtures of known composition, immuno-panned cells, and large-scale post-mortem transcriptome data. We also found that for brain transcriptomes, biological factors outweighed technical factors, and among biological factors *in vitro* culturing (Supplementary Fig.33-35) and brain region (cortex *vs.* cerebellum) (Supplementary Fig.33-35) had the strongest impact. *In vitro* culturing also affected the performance of deconvolution for other tissues (heart and pancreas), but to different extents (Supplementary Fig.37), highlighting the importance of tissue-specific benchmarking.

We found that snRNA-seq derived cell-type signatures performed well, particularly the Human Cell Atlas data (CA), which has high-coverage, while low sequencing depth (LK) led to reduced accuracy. Removing compartment-specific genes from the snRNA-seq signatures improved the deconvolution accuracy (Supplementary Fig.25).

Another factor known to influence deconvolution accuracy is the absence of cell-types present in mixtures from the signature data^51,52^. Consistent with previous results^51,52^, we found that if an abundant brain cell-type was missing from the signature data, the deconvolution accuracy was reduced, particularly for cell-types highly correlated with the missing cell-type (Supplementary Fig.19). The absence of a lowly-abundant cell-type, such as microglia and endothelia, had a minimal impact on deconvolution accuracy, suggesting that signature datasets missing these cell types can be used in deconvolution of brain data.

Since neuronal sub-types are highly similar in gene expression profiles, we investigated how different deconvolution methods handled co-linearity in brain transcriptome data. We found that CIBERSORT best handled co-linearity, and deconvolution of brain cell sub-types was accurate provided that they were not lowly abundant (<2%) or highly collinear with other cell-types (*rho* > 0.95); (Supplementary Fig.14,18).

It was previously shown that semi-supervised and unsupervised complete deconvolution methods underperform relative to supervised (*i.e.* partial) deconvolution methods^51,52^. Our results support these observations, and we further determine that the range of cell-type composition across samples in the bulk dataset is a major factor influencing the performance of complete deconvolution methods (Figure 3, Supplementary Fig.27-30).

When assessing the interplay between cellular composition and DE analyses, we found that false-positives are induced in DE analyses by as low as 5-10% difference in cell-type composition (Figure 5A). Inclusion of cell-type composition estimates as covariates effectively eliminated composition-induced false-positive genes, and restored discriminatory power for gene expression differences of 2-fold when cell-type composition differences were up to ∼25% (Figure 5C,D, Supplementary Fig.31).

The deconvolution of large GTEx data and PsychENCODE data showed that the best-performing signature may differ across datasets, and thus it is worth assessing goodness-of-fit for multiple signatures when deconvolving brain gene expression data. Notably, in both datasets, and across all deconvolution methods, there was a wide range of estimated cell-type proportions in any given brain region (Supplementary Note). This is consistent with data from the PsychENCODE consortium^15^, which used an NLS-based approach (similar to the one implemented in *drs*) and reported a similarly wide range of proportion of neurons across 1867 dorsolateral prefrontal cortex samples: 2-54%. (http://resource.psychencode.org, PEC_DER-24_Cell-Fractions-Normalised). Such a wide range is also observed in brain methylome deconvolution^53^ (0-50%) and likely reflects technical variability in dissection rather than biological inter-individual variability.

Overall, for deconvolution of brain transcriptome data we recommend that **(a)** CIBERSORT and either dTangle or MuSiC are good choices of methods **(b)** cell-type signature data should be well matched to the bulk samples, in terms of *in vitro* culture state and brain region, **(c)** cellular sub-types should only be included in deconvolution if they are > 2% abundant and < 95% correlated with other cell-types/sub-types, **(d)** when using snRNA-seq based signatures, removal of nuclear-specific genes (Supplementary Table 1) from the signature should be considered, **(e)** only attempt to use reference-free deconvolution methods if the bulk dataset is known to have a wide range of cell-type compositions.

To facilitate the choice of cell-type signature data, we provide the ten cell-type signatures compiled here as an R package (https://github.com/Voineagulab/brainyR), and developed a web tool which implements the best performing algorithms and all the cell-type signatures, as well as calculation of goodness-of-fit, in a user-friendly format, available at: https://voineagulab.shinyapps.io/BrainDeconvShiny/.

## Methods

### Datasets accessed and pre-processing

#### • Brain RNA-seq datasets

**Bulk brain gene expression data from Parikshak *et al.***^28^ were obtained from Github (https://github.com/dhglab/Genome-wide-changes-in-lncRNA-alternative-splicing-and-cortical-patterning-in-autism/releases). Exon-level count data was obtained for 251 post-mortem samples (rRNA-depleted), including frontal cortex, temporal cortex, and cerebellar vermis samples from 48 ASD and 49 control individuals, aged 2-67 (Supplementary Table 5; see Parikshak *et al*. (2016) for complete metadata).

Gene-level normalised data was generated by aggregating exon counts followed by reads per kilobase per million reads (RPKM) normalisation using the total exonic length of each gene (Ensembl V19 (hg19) assembly). A minimum expression threshold was then set at > 1 RPKM in at least 40 samples (*i.e.*, half of the number of samples in the least-represented region).

Outlier samples removed in the Parikshak *et al.* study were also removed from our analyses, leaving 121 ASD (43 frontal cortex, 39 temporal cortex, 39 cerebellum) and 126 control (45 frontal cortex, 36 temporal cortex, 45 cerebellum) samples; Supplementary Table 5.

**Bulk brain gene expression data from GTEx**^14^ were obtained as gene-level read counts from the 2016-01-05 release (V7) at https://gtexportal.org/home/datasets. Counts were RPKM normalised as above. A minimum expression threshold was set at > 1 RPKM in at least 88 samples (*i.e.* the number of samples in the least-represented brain region).

#### • Brain cell-type-specific gene expression datasets and generation of cell-type signatures

Information about samples used and final expression values are available in in Supplementary Tables 5 and 6, respectively. Metrics of signature similarity are presented in Supplementary Figures 38-39.

***F5*** *(FANTOM5)*: Cap Analysis of Gene Expression (CAGE) data for robust CAGE peaks was obtained from the FANTOM5 consortium: http://fantom.gsc.riken.jp/5/data/44. Tag-per-million normalised CAGE peak expression levels were aggregated by sum at gene level. Data from cultured neuron (n=3) and astrocyte (n=3) samples were averaged to generate the F5 neuron and astrocyte signatures. A minimum expression threshold was set at > 1 tag-per-million in at least one cell-type.

***IP*** *(immuno-purified)*: RNA-seq data from cells immunopurified from human adult brain tissue extracted during surgery were obtained from Zhang *et al.* 2016^38^. FPKM-level data were accessed from Table S4 of Zhang *et al*. for neurons (n=1), astrocytes (n=12), oligodendrocytes (n=5), microglia (n=3), endothelia (n=2). Cell-types derived from foetal brain were excluded (*i.e.*, foetal astrocytes). Samples of the same cell-type were averaged to generate the IP signature. A minimum expression threshold was set at > 1 FPKM in at least one of the five cell-types in the final signature matrix.

***MM*** *(Mus musculus)*: RNA-seq data from immunopurified mouse brain tissue was obtained from Zhang *et al.* 2014^43^. FPKM-level data were accessed from https://web.stanford.edu/group/barres_lab/brain_rnaseq.html, in which biological replicates of cell-type transcriptomes (neurons, astrocytes, oligodendrocytes, microglia, and endothelia were already aggregated across samples. Mouse genes were mapped to human orthologues using Gene ID homology information from http://www.informatics.jax.org/downloads/reports/HOM_MouseHumanSequence.rpt.

Expression data from oligodendrocyte precursors and newly-formed oligodendrocytes were excluded. A minimum expression threshold was set at > 1 FPKM in at least one of the five cell-types in the final signature matrix.

***DM*** *(Darmanis):* Human brain single-cell gene expression data from the middle temporal gyrus generated by Darmanis *et al.* (2015)^13^ were downloaded as count-level data from https://github.com/VCCRI/CIDR-examples/tree/master/Brain54. To generate the DM signature, RPKM or counts-per-million (CPM) expression was averaged across samples of each cell-type (*i.e.* astrocyte (n = 62), neuron (161), microglia (16), mature oligodendrocytes (38), oligodendrocyte precursor cells (OPCs) (18), or endothelia (20). Cell-types derived from foetal brain (quiescent neurons and replicating neurons) were excluded. A minimum expression threshold was set at > 1 RPKM or CPM in at least one cell-type in the final signature matrix.

***LK*** *(Lake)*: Gene expression data for 10,319 human adult frontal cortex nuclei were accessed from Lake *et al*. 2018^41^. Seurat^55^ was used to pre-process raw count expression data, removing nuclei with 1) fewer than 1000 counts or 2) 200 expressed genes, or 3) >5% of counts attributed to mitochondrial genes, or 4) a number of reads >99.5^th^ percentile of its dataset. Only 3930 nuclei passed these QC criteria. To generate the LK signature, RPKM or CPM values were averaged across nuclei of each cell-type: astrocytes (97), excitatory neurons (2611), inhibitory neurons (1051), oligodendrocytes (96), OPCs (46), and microglia (22). An expression profile for neurons was also generated, as the average of all excitatory and inhibitory nuclei. A minimum expression threshold of > 1 RPKM or CPM in at least one cell-type was required. Note that endothelia were excluded for having fewer than 10 nuclei (7).

***VL*** *(Velmeshev)*: 10X Chromium for single-nucleus data from the post-mortem adult human brain were accessed Velmeshev *et al.*^36^. Only nuclei from control prefrontal cortex samples were included. Seurat processing, cell-type aggregation, and thresholding were performed as described above in LK After filtering, 24,556 nuclei remained, classified as astrocytes (2229), excitatory neurons (9718), inhibitory neurons (4238), oligodendrocytes (4721), OPCs (2677), microglia (450), and endothelia (523).

***CA*** *(Cell Atlas):* Count-level exon expression data for NeuN+ sorted adult nuclei from the middle temporal gyrus were acquired from the Human Cell Atlas^37^. Seurat processing, cell-type aggregation, and thresholding were performed as described above in LK. After filtering, 15,524 nuclei remained, classified as astrocytes (291), excitatory neurons (10492), inhibitory neurons (4118), oligodendrocytes (313), OPCs (238), microglia (63). Endothelia were excluded for having fewer than 10 representatives (9).

***NG*** *(Nagy)*: 10X Chromium single-nucleus expression data from the adult human post-mortem human prefrontal cortex were accessed from Nagy *et al.*^40^ . Only nuclei from control samples were included. Seurat processing, cell-type aggregation, and thresholding were performed as described above in LK. After filtering, 23,168 nuclei remained, classified as astrocytes (1195), excitatory neurons (14624), inhibitory neurons (5940), oligodendrocytes (757), OPCs (505), microglia (85), and endothelia (62).

***TS*** *(Tasic)*: Exon-level SmartSeq2 single-cell expression data from the adult mouse cortex were accessed from Tasic *et al.*^42^. Only cells from the Anterior Lateral Motor Cortex were included. Further, cells labelled by the authors as low quality or with no class were excluded. Seurat processing, cell-type aggregation, and thresholding were performed as described above in LK. After filtering, 8075 nuclei remained, classified as astrocytes (195), excitatory neurons (3851), inhibitory neurons (3767), oligodendrocytes (69), OPCs (24), microglia (80), and endothelia (89).

**MB** (*Multibrain*): this composite signature was generated by quantile-normalising and averaging the RPKM-level expression of the CA, IP, DM, NG, and VL signatures for five cell-types (neurons, astrocytes, oligodendrocytes, microglia, and endothelia). All signatures are cortical in origin but represent a range of purification protocols (scRNA-seq by SmartSeq (DM), snRNA-seq by 10X (VL, NG), snRNA-seq by SmartSeq (CA), and immuno-panning (IP))

#### • Heart and pancreas RNA-seq datasets

**Bulk gene expression data from GTEx**^14^ for pancreas (n=268) and heart (n=310 and 417 atrial appendage and left ventricle, respectively) were processed as per the GTEx brain samples, except the pancreas samples were normalised to the level of transcripts-per-million (TPM)

**Cell-type-specific RNA-seq data from pancreas alpha and beta cells** were obtained from three studies as described below. For each dataset, genes were excluded if they were not protein-coding, or if they were expressed at < 1 TPM in both cell-types.

***EN*** *(Enge)*: count-level expression data for single-cells from freshly-isolated, FACS-sorted human pancreas were acquired from Enge *et al.*^56^. Data were normalised to the level of transcripts-per-million (TPM), using the total exonic length of each gene per the Ensembl V19 (hg19) assembly. The expression signature of alpha and beta cells was generated as the average of 998 alpha and 348 beta cells.

***BL*** *(Blodgett)*: TPM-level expression data for bulk RNA-seq from freshly-isolated, FACS-sorted alpha and beta cells from human pancreas were acquired from Blodgett *et al*.^11^. The expression signature of alpha and beta cells was generated as the average of 7 adult alpha-cell and 7 adult beta-cell bulk RNA-seq samples.

***FS*** *and **FG** (Furuyama)*: count-level expression data for human pancreas alpha and beta cells were acquired from Furuyama *et al.*^12^. After TPM normalisation, the **FS** (Furuyama Sorted) signature was constructed from freshly-isolated, FACS sorted alpha and beta cells (average of 5 replicates each), while the **FG** (Furuyama GFP) signature consists of isolated alpha and beta cells subjected to 1-week of culturing. These cells had been transduced with a GFP expression vector for imaging purposes^12^ (average of 4 and 6 replicates, respectively).

**Cell-type-specific RNA-seq data from heart** were accessed from three publicly available datasets, containing cardiomyocytes (CM), cardiac endothelia (EC), cardiac fibroblasts (FC), and smooth muscle cells (SMC). For each dataset genes were excluded if they were not protein-coding, or if they were expressed at < 1RPKM across all four cell-types.

***F5*** *(FANTOM5):* Cap Analysis of Gene Expression (CAGE) data for robust CAGE peaks was obtained from the FANTOM5 consortium: http://fantom.gsc.riken.jp/5/data/44. Tag-per-million normalised CAGE peak expression levels were aggregated by sum at gene level. n=3, 4, 6, and 3 for CM, EC, FC, and SMC, respectively.

***EN*** *(ENCODE):* FPKM-level RNA-seq data for cultured primary cells were accessed from the ENCODE consortium^57^; n=2 for all cell-types.

***SC*** *(Single-cell):* single-cell RNA-seq data from freshly-isolated tissue samples were accessed from Wang *et al.* (2020)^58^ (GSE109816). Only left atrial samples were used. Cell-type specific expression was generated as the average RPKM of all cells in each classification. n=1934, 1111, 257, and 427 for CM, EC, FC, and SMC, respectively.

### RNA-seq data generated in the present study and data pre-processing

#### • RNA Mixture Experiment

Total RNA was extracted from human primary astrocytes and from neurons derived from human foetal neural progenitors.

Human primary astrocytes (Lonza, #CC-2565) stably expressing GFP from pCMV6-AC-GFP had been generated by selection with G418 (Thermo Fisher Scientific, #10231027) at 800μg/ml. Cells were cultured in RPMI GlutaMAX™ (Thermo Fisher Scientific, #35050061) supplemented with 10% foetal bovine serum, 1% streptomycin (10,000 μg/ml), 1% penicillin (10,000 units/ml) and 1% Fungizone (2.5 μg/ml) and seeded into 6-well tissue culture plates at a density of 0.5 × 10^6^ cells 24 hours prior to RNA extraction. Total RNA was extracted using TRIzol® reagent and a Qiagen miRNeasy kit and treated with 1 µl DNase I (Thermo Fisher Scientific, #AM2238) per 10 μg of RNA.

Neuronal differentiation of human neural progenitors stably transfected with pLRC-GFP was carried out for 2 weeks as previously described^59^. RNA extraction was carried out using a Qiagen miRNeasy kit, with on-column DNase digestion. RNA from differentiated neurons was kindly provided by Dr. Brent Fogel (UCLA)^59^.

RNA mixtures were generated by mixing neuronal and astrocyte RNA in mass ratios of 40:60, 45:55, 50:50 neuron:astrocyte (n=1 for each ratio). In addition, a pure neuronal RNA sample and pure astrocyte RNA samples (n=3) were also included (Supplementary Table 7).

Library preparation using the Illumina TruSeq Stranded kit (http://www.illumina.com/products/truseq_stranded_total_rna_library_prep_kit.html) and sequencing on a NextSeq 500 Illumina sequencer were carried out at the UNSW Ramaciotti Centre for Genomics, generating 75 bp paired-end reads (Supplementary Table 7). Sequencing reads were mapped to the human genome (hg19) using STAR v2.5.2b^60^ with the following parameters: --outSJfilterOverhangMin 5 5 5 5 --alignSJoverhangMin 5 -- alignSJDBoverhangMin 5 --outFilterMultimapNmax 1 --outFilterScoreMin 1 -- outFilterMatchNmin 1 --outFilterMismatchNmax 2 --chimSegmentMin 5 --chimScoreMin 15 --chimScoreSeparation 10 --chimJunctionOverhangMin 5.

Gene counts for GENCODE V19 annotated genes were obtained from the STAR output and RPKM-normalised.

***IH (in house) cell-type signature*** data includes the RPKM-normalised data from the the pure neurons and astrocyte samples (averaged across the 3 replicates for astrocytes). Data was thresholded for a minimum of 1 RPKM in at least one cell-type.

***RNA mixture data*** consists of RPKM-normalised data from the three RNA mixture samples. Genes expressed at < 2 RPKM in at least one sample were filtered out.

#### • Bulk RNA-seq data generated from brain tissue

Brain tissue samples were obtained from the NICHD Brain and Tissue Bank, and included frontal cortex samples (BA9/10) from 2 control, 2 ASD, and 1 Fragile-X premutation carrier individuals. For each brain sample, frozen tissue was pulverised using a CellCrusher (https://cellcrusher.com/) and the tissue was then divided for nuclear RNA extraction and RNA extraction from bulk tissue.

### Nuclei Isolation

Around 30 mg of tissue was lysed in 2.5 ml lysis buffer (10 mM Tris-HCl, 3 mM MgCl2, 10 mM NaCl, 0.005% NP40) for 17 minutes on ice. After lysis, 2.5 ml of ice-cold PBS was added to the sample and tissue was homogenized using a Pasteur pipette until no large chunks were visible. Tissue was then filtered through a 30 µm strainer and centrifuged at 500 g for 5 minutes at 4°C. Supernatant was removed and the pellet was resuspended in 400 µl PBS with 1% BSA and DAPI. DAPI-positive singlet nuclei were sorted using a BD Influx with a 70 µm nozzle at 20 PSI to collect approx. 100,000 nuclei per sample.

### Bulk RNA extraction and library generation

To extract RNA from bulk tissue, the Qiagen mini RNA prep kit was used following the manufacturer’s instructions, including a DNAse treatment step. From sorted nuclei, RNA was extracted by a hot Trizol extraction method. Nuclei were washed in PBS and resuspended in Trizol at 65°C and incubated on a shaker at 1,300 rpm for 15 min. RNA was enriched using a guanidinium HCl buffer and silica-coated magnetic beads with a DNAse I treatment step. RNA amounts and quality were assessed on a TapeStation using RNA Screen Tape (Agilent), and 20-100 ng of total RNA was used per replicate to generate RNA-seq libraries. ERCC ExFold RNA Spike-In mixes (Thermo Scientific) were added as internal control. Libraries were prepared using the TruSeq Stranded mRNA library prep kit (Illumina), using TruSeq RNA unique dual index adapters. Libraries were quantified by qPCR on a CFX96/C1000 cycler (Biorad) and sequenced on a Novaseq 6000 (Illumina) for 2x 53 bp as paired-end, generating around 25 M reads per sample.

Sequencing reads were mapped to the human genome (hg38) using STAR v2.5.2b^60^ with the following parameters: --outSJfilterOverhangMin 15 15 15 15 --alignSJoverhangMin 15 --alignSJDBoverhangMin 15 --outFilterMultimapNmax 1 --outFilterScoreMin 1 --outFilterMatchNmin 1 --outFilterMismatchNmax 2 --chimSegmentMin 15 --chimScoreMin 15 --chimScoreSeparation 10 --chimJunctionOverhangMin 15 --bamRemoveDuplicatesType UniqueIdenticalNotMulti. Note that nuclear samples were mapped to a pre-mRNA hg38 transcriptome.

Gene counts for GENCODE V19 annotated genes were obtained from the STAR output and RPKM-normalised.

Nuclear enrichment was confirmed using the expression of the nuclear-specific transcript MALAT1 (22.1-fold enrichment in nuclear samples, p = 6.7×10^-5^, t-test; Supplementary Table 8)

#### • Single-nucleus RNA-seq data generated from bulk brain tissue

snRNA-seq data were generated from the same five brain samples described in the previous section, but from a different chunk of the dissection.

##### Nuclei Isolation

Around 30 mg of tissue was lysed in 400 µl of lysis buffer (10 mM Tris-HCl, 3 mM MgCl2, 10 mM NaCl, 0.005% NP40) in 1.5 ml tubes and broken down with a pellet pestle. Tissue was dissociated by passing through a polished silanized Pasteur pipette 3-4 times, then incubated on ice for 10 minutes. Dissociation was repeated at 5 minutes and 10 minutes. After incubation, the dissociated tissue was added to 2.5 ml of wash buffer in a 15 ml falcon tube. The sample was then passed through a 30 µm strainer into a 50 ml falcon tube and centrifuged for 5 minutes at 500 x g at 4°C in a swinging bucket centrifuge. Following centrifugation, the supernatant was removed and sample resuspended in 100 µl of wash buffer (PBS with 1% BSA) for every 30 mg of tissue used. Using only 100 µl of the resuspended sample, 180 µl of a 1.8 M sucrose solution (made with Sigma Nuclei Pure Prep kit) was added and homogenized using a P1000 pipette. In a 2 ml Eppendorf tube, 1 ml of a 1.3 M sucrose solution with 1% BSA was placed. 280 µl of the nuclei suspension mixed with sucrose was slowly layered on top of the 1.3 M sucrose solution. The sucrose gradient was centrifuged for 10 minutes at 3,000 x g at 4°C in a swinging bucket centrifuge. After centrifugation, the debris from the top of the sucrose gradient were removed by soaking a Kimwipe from the top of the tube, slowly lowering it together with the sinking meniscus until a volume of less than 100 µl remained, which was removed with a pipette. Nuclei were resuspended in 20-50 µl of wash buffer and 10 µl of the suspension was stained with Trypan Blue to count for concentration.

##### Library generation and sequencing

The 10X Genomics 3’ v2 and v3 single cell expression kit was used to generate single nuclei RNA-seq libraries. Using 16,000 nuclei in total to aim for 10,000 nuclei recovery, the standard protocol was used according to manufacturer instruction with the following alterations: 17 PCR cycles in total for cDNA amplification and 13 PCR cycles in total for library amplification. Libraries were then sequenced on a Novaseq 6000 (Illumina) generating around 100 M reads per sample.

##### Sequence Alignment and UMI counting

A pre-mRNA transcriptome was built using the Cell Ranger mkref command and default parameters starting with the refdata-cellranger-GRCh38-1.2.0 transcriptome as per the instructions provided by 10X Genomics. Reads were demultiplexed by sample index using the Cell Ranger mkfastq command. Fastq files were aligned to the custom transcriptome, cell barcodes were demultiplexed, and UMIs corresponding to genes were counted using the cell ranger count command using default parameters. Cell Ranger version 2.1.0 was used for all steps.

##### Data preprocessing

Cell Ranger output was pre-processed using Seurat v3^55^. Filtering-out criteria for nuclei: < 500 counts, or < 200 expressed genes, or >20% of counts attributed to mitochondrial genes, or total number of reads in the top 99.5^th^ percentile of its dataset. UMI counts were log2-transformed and normalised for library size and mitochondrial percentage, and finally scaled. Nuclei from all individuals were then integrated using canonical correlation analysis in Seurat, setting the numbers of dimensions to be 30.

After retransformaing and renormalising data, clustering was performed using tSNE^61^ on the top 35 principal components of the 2000 most variable genes, with the resolution parameter set to 1.5 (Supplementary Fig.40). Clusters were annotated using SingleR^62^ to transfer cell-type annotation labels from the NG signature.

##### Signature generation

A separate cell-type specific signature was generated from each of the five individuals. This was calculated as the average RPKM of each individual’s cells within each cluster. Only cell-types represented in all individuals were used (Neurons, Astrocytes, and Oligodendrocytes).

### Simulated datasets

#### • Simulations for assessing deconvolution accuracy

**Randomly sampled single-nucleus mixtures** were generated using data from VL^36^ and the CA^37^ datasets.

Simulated data was generated separately from each dataset. Seurat v3 was used to pre-process raw count expression data, removing nuclei with 1) fewer than 1000 counts or 200 expressed genes, 2) >5% of counts attributed to mitochondrial genes, or 3) a number of reads >99.5^th^ percentile of its dataset. In addition, cells assigned to a cell-type or cell-subtype with fewer than 200 cells were excluded. Next, the dataset was randomly split into two: half was used to generate cell-type signatures and half for simulated mixture. One hundred mixtures were simulated by summing the counts of 500 randomly-sampled single nuclei. Random sampling was performed without replacement.

**Randomly sampled single-cell mixtures** were generated using single-cells from the Darmanis *et al.* dataset^13^, using a method largely as above but with three key differences: first, only cells classified as one of neurons, astrocytes, oligodendrocytes, OPCs, microglia, or endothelia were included, without regard for the number of representatives (*i.e.,* non-hybrid cells from adult samples); second, the number of cells aggregated per mixture was only 100, owing to the lower number of total cells (285); and finally, the dataset was not randomly split into two for mixture and signature generation.

We confirmed that the single-nucleus and single-cell simulated mixtures, had similar expression distributions to data from bulk brain tissue, and were not zero-inflated (Supplementary Fig.41).

**Single-nucleus mixtures with a wide range of cell-type compositions** were generated using single nuclei from the VL and CA datasets. 100 mixtures were simulated. To obtain a defined range of cell-type proportions in the mixture, for each cell-type *j* we randomly sampled without replacement between 1 and *n_j_* nuclei where *n_j_* = (*n/k*)/(*s_j_*/min(*s*)) where *n*=500, the chosen number of cells per mixture; *k* is the number of cell-types in each dataset; *s* is the vector of total library sizes for all *k* cell-types; *s_j_* is the total library size for cell-type *j*.

If more than 500 total nuclei were randomly-sampled by this approach, then a random subset of 500 was kept; conversely, if fewer than 500 nuclei were initially sampled, then additional nuclei were randomly-sampled from any cell-type until 500 was reached. Mixtures were simulated by summing the counts of these single nuclei followed by counts-per-million normalisation.

#### • Simulations for assessing the interaction between composition and differential expression

##### Simulated data with cell-type composition differences between sample groups

Single-nucleus mixtures for DE analyses were generated using snRNA-seq data from the CA dataset. Nuclei were classified as one of Excitatory, Inhibitory, Oligodendrocyte, OPC, or Astrocyte. Each simulation was created as a dataset of 100 samples, split into groups A and B of 50 samples each. Each sample in group A (the reference group) was generated as the summed expression of randomly selected *n* excitatory neurons and 500-*n* non-excitatory cells, where *n* was a randomly selected integer from [200-300] so that the simulated proportion of excitatory neurons varies between 40-60%). Samples in group B (test group) were generated as per group A, except n was sampled from [200+*k*, 300+*k*] for increased proportions or [200-*k*, 300-*k*] for decreased proportions where *k* varied from 0 to 195 with a step of 5. All sampling was performed without replacement. Differential expression analyses for group B *vs.* group A were performed on each dataset as described in the “Differential Expression” section below.

##### Simulated data with cell-type composition and gene expression differences between sample groups

The expression of expression of 200 genes was altered by 1.1-, 1.3-, 1.5-, or 2-fold in the above simulated mixtures, in group A samples only. The 200 genes selected for perturbation included the top 100 excitatory neuron marker genes and 100 randomly-selected non-marker genes. Half of each set was randomly assigned to be upregulated or downregulated.

To simulate cell-type-specific expression differences, the expression alteration was introduced only to nuclei from the cell-type-of-interest (*i.e.* excitatory or inhibitory neurons) prior to aggregation.

### Estimation of cellular composition

#### • Overview of deconvolution methods

In general, deconvolution methods model gene expression data from a tissue sample (vector *X*) as the sum of gene expression levels in the cell-types of which it’s comprised (“signature” expression matrix, *S*), weighted by the proportion of each cell-type in the sample (vector *P*), formalized as *X ∼ S*P.* Deconvolution methods fall into two broad categories – partial and complete – as described below.

***Partial or supervised deconvolution***^6,18,29,31,63–68^ estimates the proportion of cell-types in a sample based on experimentally measured gene expression values from pure cell-types, *i.e.* determines P knowing X and S.

It is worth noting that the signature expression data (S) often comes from a different source than the bulk tissue data (X), and thus an intrinsic assumption of most partial deconvolution methods is that gene expression in a given cell-type is the same regardless of the source of cells (thus genetic background and environmental conditions including culture conditions are ignored)^1,68^. The most frequently employed methods for partial deconvolution are Non-negative Least Squares (*i.e.* optimising *X ∼ S*P* using a least-squares approach where P should be non-negative) (*e.g.* DeconRNASeq^29^), and Support Vector Regression (*e.g.* CIBERSORT^18^).

A simplified version of partial deconvolution consists of calculating an enrichment score, rather than a proportion, for each cell-type (*e.g.* xCell^19^, or BrainInABlender^7^). While this approach is intuitive, it has several limitations: its accuracy is harder to assess (as one cannot calculate error measures or goodness-of-fit), and its biological interpretation is often unclear since the scale of enrichment scores is variable.

***Complete or reference-free/unsupervised deconvolution*** consists of estimating both the proportion of cell-types and cell-type specific expression, *i.e.* determining both P and S knowing X^33–35,45,69^. This is an under-determined problem, which requires biologically motivated constraints.

#### • Deconvolution methods used

Cell-type composition was estimated using four partial deconvolution methods **(DeconRNASeq**^29^, **dtangle**^31^, **MuSiC**^30^, and **CIBERSORT**^18^), two enrichment methods with in-built signatures (**BrainInABlender**^7^ and **xCell**^19^), and two complete deconvolution methods: **Linseed**^33^, and a co-expression based approach proposed by Kelley *et al.*^5^ (referred to as **Coex**).

All algorithms were run in R v3.6. All data used for deconvolution were RPKM-normalised expression values without log2 transformation^70^ unless noted below.

**CIBERSORT v1.04** was run using the *CIBERSORT* R package obtained from https://cibersort.stanford.edu with default parameters.

**DeconRNASeq v1.26** was run using the *DeconRNASeq* Bioconductor R package with default parameters.

**Music** v0.1.1 was run using the music_prop() function from R package available at https://github.com/xuranw/MuSiC. Raw count data was used as input for both signatures and mixtures. Only single-cell- or single-nucleus-derived signatures were used; their individual cells/nuclei were not aggregated, metadata about the individual-of-origin was included as well as predefined cell-type labels.

**dtangle v0.3.1** was run using the *dtangle* CRAN R package. Cell-type markers were selected as the top 1% of markers using its *find_markers* function with method=”diff”. Data was log2 transformed with an offset 0.5, as recommended^31^.

**BrainInABlender v0.9** was run using the R package obtained from https://github.com/hagenaue/BrainInABlender using default parameters. Cell-type signature data built into BrainInABlender is derived from numerous resources of brain cell-type specific expression, including human data from Darmanis *et al.*^13^, and various mouse datasets (full list in Hagenauer *et al.,* 2018). Both publication-specific indices and an averaged index are generated; we used the averaged index as the enrichment score in all analyses.

**xCell v1.1.0** was run using the R package from https://github.com/dviraran/xCell using default parameters with the built-in signature data. Cell-type signature data for neurons and astrocytes are built in xCell, and are derived from *in vitro* cultured data from FANTOM5^44^, and ENCODE^57^. xCell generates a “Raw” and a “Transformed” enrichment score; we used the latter as a measure of enrichment.

**Coex** was carried out by constructing co-expression networks using the *blockwiseModules* function from the WGCNA R package^46,71^, with the following parameters: deepSplit = 4, minModuleSize = 150, mergeCutHeight = 0.2, detectCutHeight = 0.9999, corType = “bicor”, networkType = “signed”, pamStage = FALSE, pamRespectsDendro = TRUE, maxBlockSize = 30000. The beta power was selected for each network so that the scale-free topology fit *r*^2^ was > 0.8 and median connectivity < 100 (Supplementary Information Code). Genes were assigned to the module with the highest kME (correlation with the module eigengene), provided *kME* > 0.5, and *p* < 0.05 (BH-corrected Student’s t-test). Co-expression networks were built on log2 transformed RPKM values, offset by 0.5.

A cell-type module (CTM) was defined as the module most significantly enriched for the top 100 markers of a given cell-type, requiring an enrichment p-value <10^-5^ and odds ratio >5.

Enrichment was assessed using a one-sided Fisher’s Exact Test. Cell-type markers were defined using the *find_markers* function in the dtangle R package applied to the matching cell-type signature data for simulations, and MB for GTEx and Parikshak. Cell-type enrichment scores were defined as the CTM’s eigengene values (*i.e.*, first principal component values of genes included in the CTM), as per Kelley *et al.*^5^.

**Linseed** v0.99.2 was run using the R package from https://github.com/ctlab/LinSeed. We used a collinearity threshold of p=0.01 to filter genes. Output was transformed to sum-to-one.

We also tested the SVD approach to determine the number of cell-types in the mixture data, which involves looking for the plateau (Supplementary Figure 26). For the VL-based simulations, with 7 cell-types, the estimated k was greater than 10 for random and 7 for wide-range mixtures. For the CA-based simulations, with 5 cell-types, the estimated k was ∼5-7 for random and 5 for wide-range mixtures. For the RNA mixtures which consisted of 2 cell-types, the estimated k was 3. For the DM mixtures, which consisted of 5 cell-types, the estimated k was more than 10. Therefore, we used the known k value for all mixtures.

#### • Deconvolution of specific datasets

Parikshak, GTEx and RNA mixture data were deconvolved using RPKM-normalised signatures and mixtures, while for single-cell and single-nucleus simulated datasets, signatures and mixtures were CPM-normalised, if raw count data was available (otherwise the normalised data available from the original publication was used).

### Assessment of deconvolution accuracy

For simulated datasets, deconvolution accuracy was assessed by two measures: (i) Pearson correlation between true and estimated proportions and (ii) normalised mean absolute error (nmae) calculated as mean error divided by the mean of true proportions, where error is the per-sample absolute difference between estimate and true proportion. Note that nmae can only be calculated when estimates are bounded between 0-1 *i.e.* are proportions rather than relative enrichment scores like Blender’s or xCell’s output.

For datasets without a ground truth, such as bulk brain samples, goodness-of-fit was evaluated as the Pearson correlation for each sample’s reconstructed and observed expression, log2-transformed with an offset of +0.5.

First, observed expression and cell-type signature data were quantile normalised. Then, for each sample, reconstructed expression values were calculated using the following formula:

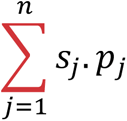

where, *j* denotes a cell-type, *s_j_* is the vector of gene expression in cell-type *j* (from the signature matrix), and *p_j_* is the estimated proportion of cell-type *j* in the sample, and *n* is the number of cell-types.

### Differential expression (DE) analyses

#### Differential expression analyses in simulated data

DE between group A and group B in simulated single-nucleus mixtures was assessed using either a linear model on log2-transformed CPM values offset by +0.5 as implemented in the *lm* function in R, or a generalised linear model implemented in DESeq2^47^ on count data.

Excitatory neuron proportions were included as covariates in the model either as linear term, a quadratic term, or after conversion to a spline matrix using the bs() function from the R *splines* package, setting degree = 3 and knots at its 25^th^, 50^th^, and 75^th^ percentiles.

Multiple testing correction was conducted using the Benjamini-Hochberg approach^72^.

Cell-type marker enrichment analyses were performed by one-sided Fisher’s exact test for 100 markers per cell-type using the CA cell-type signature. Markers were defined using the *find_markers* function from dtangle^31^, setting marker_method = “diff”. Only simulations where the number of false positive genes was >100 were tested for cell-type enrichment, to ensure adequate power for the test.

Discriminatory power was calculated as the fraction of the 200 true perturbed genes that were in the top 200 most significant genes by p-value. No significance threshold was applied to DE p-values.

#### Cell-type-specific DE analyses

Cell-type-specific expression was first extracted using the high-resolution algorithm of CIBERSORTx^48^ webtool at https://cibersortx.stanford.edu/ with default settings using the CA signature. Due to the webtool’s computational constraints, a subset of 1000 genes was analysed, including the 100 perturbed cell-type marker genes, the 100 perturbed non-marker genes, the top 100 markers for each of the four other cell-types, and 400 randomly-selected non-marker genes. The resultant cell-type-specific expression data was used for DE analyses, using a linear model on log2 transformed data offset by +0.5. Multiple testing correction was conducted using the Benjamini-Hochberg approach, adjusting for the size of the full transcriptome rather than that of the smaller subset^72^.

#### DE analyses of ASD and control samples

DE was carried out using DESeq2 v1.22.2^47^ on count-level expression data. The same samples used by Parikshak *et al.*^28^ for DE were included in our analyses: 106 samples (43 ASD, 63 controls; Supplementary Table 5). Differential expression was carried out using a Wald test with Benjamini-Hochberg correction for multiple testing as implemented in DESeq2^47^. Composition-dependent DE adjusted for the following covariates: Age, Sex, Sequencing Batch, Brain Bank, Region, RIN, and the first two principal components of sequencing metadata, per Parikshak *et al.*^28^. Composition-independent DE used the same covariates as above, but adding the estimated proportions of astrocytes and any other cell-types not significantly correlated with astrocyte proportions (p<0.05, Pearson correlation test) *i.e.,* oligodendrocyte and microglia (Supplementary Figure 42), to minimise co-linearity.

#### Determination of compartment-specific genes

Compartment-specific genes were identified using the RNA-seq data generated from bulk brain tissue (5 total RNA and 5 nuclear RNA samples) pre-processed as described above, and log2-transformed with an offset of 0.5. Compartment specific genes were identified as genes DE between groups using a linear model as implemented in the *lm* function in R (absolute fold-change > 1.3 and a Benjamini-Hochberg-adjusted^72^ p-value < 0.05).

### Other analyses

Gene ontology (GO) and pathway enrichment analyses were conducted using gProfiler2 v0.2^73^ in R, setting exclude_iea=TRUE and all other parameters as default. P-values were BH-corrected^72^. Only results from GO, KEGG, Reactome, Human Phenotype, and Wikipathways were reported, with filtering performed after multiple-testing correction.

For all set enrichment analyses, the background was set to the relevant list of all expressed genes.

### Note on cell-type proportions vs. RNA proportions

Since cell-types differ in their total RNA content, transcriptome deconvolution estimates proportions of RNA from each cell-type, rather than proportions of cells *per se*^33^. It is important to note that bulk RNA-seq sequences a mixture of RNA molecules (primarily protein coding, after poly-A selection or ribo-depletion), and thus the goal of transcriptome deconvolution is in fact to estimate the proportion of the sequenced RNA molecules coming from a given cell-type (pRNA), rather than the proportion of cells. *A-priori,* pRNA (not pCt) should be relevant for reconstruction of gene expression data, and thus useful as a co-variate in differential expression analyses. To test this hypothesis, we deconvolved pseudo-bulk data from the Velmeshev *et al.* snRNA-seq dataset (10 individuals), where we know both pCt and pRNA (calculated as the proportion of RNA-seq reads from each cell-type), and found that deconvolution estimates perfectly correlate with pRNA but less so with pCt (Supplementary Figure 43), consistent with previous data^33^. Note that pRNA and pCt are themselves correlated in this dataset (*r*=0.86). We then assessed goodness-of-fit for these pseudo-bulk data using either pRNA or pCt to reconstruct gene expression. We found that goodness-of-fit was always higher when using pRNA (Supplementary Figure 43). These data demonstrate that pRNA, the output of transcriptome deconvolution, is the appropriate measure to use for re-constructing gene expression data and thus as a co-variate in DE analyses. For simplicity, we refer to pRNA as “cell-type proportions” throughout the manuscript.

## Data availability

The sequencing data generated in the present study is available on GEO/SRA accession number GSE175772.

## Code availability

Data analysis code is available at: https://github.com/Voineagulab/BrainCellularComposition.

All brain cell-type signatures are available as an R package: https://github.com/Voineagulab/brainyR.

Deconvolution with the top performing algorithms is implemented at: https://voineagulab.shinyapps.io/BrainDeconvShiny/.

## Acknowledgements

This work was supported by an ARC Future Fellowship and a UNSW Scientia Fellowship to I.V. and an RTP PhD Scholarship to G.J.S.

## Supplementary Figure Legends

**Supplementary Figure 1. Generation and deconvolution of a replication simulated dataset using single-nuclei from the CA dataset**. **A.** Process for generating 100 in silico mixtures. **B.** Barplots show Pearson correlation coefficients (r) between true and deconvolution-estimated proportions in 100 *in silico* mixtures. The left column shows results when only major cell-type labels are used in the signature; the middle column shows results when a mix of major cell-type and cell-subtype labels are used in the signature; the right column shows results when only cell-subtype labels are used in the signature. *Dotted line*: *r*=0.8.

**Supplementary Figure 2. Benchmarking deconvolution algorithms on simulated mixtures using data from Darmanis *et al.* A.** Simulation design. **B.** Scatterplots of estimated proportion (or enrichment score) and true proportion for each cell-type. *Ast*: astrocytes. *End*: endothelia. *Mic*: microglia. *Neu*: Neurons. *Oli*: oligodendrocytes. *Red dotted line*: y=x. Grey line: regression line. **C.** Barplots of normalised mean absolute error *(nmae*; left) and Pearson correlation coefficients between true and estimated proportions (*r*; right) based on 100 *in silico* mixtures.

**Supplementary Figure 3.** Barplots of normalised mean absolute error (*nmae*) between true and deconvolution-estimated proportions in 100 VL-derived *in silico* mixtures. *Nmae* was calculated as the average error divided by the average true value. **A.** Using only major cell-type labels in the signature. **B.** Using a mix of major cell-type and cell-subtype labels in the signature. **C.** Using all cell-subtype labels are the signature. *Dotted black line*: *nmae* = 1.

**Supplementary Figure 4.** Scatterplots of true and deconvolution-estimated proportions in 100 VL-derived *in silico* mixtures. Each row represents a different algorithm. The signature used only major cell-types. *Solid black line*: regression line. *Nmae*: normalised mean absolute error. *r*: Pearson correlation coefficient. Note that nmae was not calculated for xCell and Blender as their output is an enrichment score rather than a proportion

**Supplementary Figure 5.** Barplots of normalised mean absolute error (*nmae*) between true and deconvolution-estimated proportions in 100 CA-derived *in silico* mixtures. *Nmae* was calculated as the average error divided by the average true value. **A.** Using only major cell-type labels in the signature. **B.** Using a mix of major cell-type and cell-subtype labels in the signature. **C.** Using all cell-subtype labels are the signature. *Dotted black line*: *nmae* = 1.

**Supplementary Figure 6.** Scatterplots of true and deconvolution-estimated proportions in 100 CA-derived *in silico* mixtures. Each row represents a different algorithm. The signature used only major cell-types. *Solid black line*: regression line. *Nmae*: normalised mean absolute error. *r*: Pearson correlation coefficient. Note that nmae was not calculated for xCell and Blender as their output is an enrichment score rather than a proportion

**Supplementary Figure 7. Deconvolving mixtures of RNA from cultured neurons and astrocytes. A.** Outline of RNA mixtures and its corresponding in-house (IH) signature. **B.** Scatterplots of estimated and true proportions of neurons using CIB, DRS and DTA, combined with the matching IH signature. Note that the MUS algorithm was not used, as the algorithm is only compatible with single-cell-level data. **C.** Scatterplots of neuron enrichment scores obtained with Blender (left) and xCell (right). **D.** Scatterplots of astrocyte enrichment scores obtained with Blender (left) and xCell (right). **E**. Scatterplots of true versus estimated neuronal proportion when using CIB and mismatched signatures. All signatures contained just neuronal and astrocyte expression values.

**Supplementary Figure 8. Estimated proportions in immuno-panned purified brain.** *Thick horizontal line*: mean. *Neu*: neurons. *Ast*: astrocytes. *Oli*: oligodendrocytes. *Mic*: microglia. *CIB*: CIBERSORT. *DRS*: DeconRNASeq. *DTA*: dtangle. Note that MuSiC was not applied as it requires single-cell or –nucleus data for its signature.

**Supplementary Figure 9**. Scatterplots of true and deconvolution-estimated proportions in 100 VL-derived *in silico* mixtures. The signature used a range of cell-subtypes and major cell-types. **A.** CIBERSORT deconvolution. **B.** DeconRNASeq. **C.** dtangle. **D.** MuSiC. *Solid black line*: regression line. *Nmae*: normalised mean absolute error. *r*: Pearson correlation coefficient. *Neurons_Inh and Neurons_Exc*: Inhibitory and excitatory neurons, respectively.

**Supplementary Figure 10. Scatterplots of true and deconvolution-estimated proportions in 100 CA-derived *in silico* mixtures**. Each row represents a different algorithm. The signature used a range of cell-subtypes and major cell-types. **A.** CIBERSORT deconvolution. **B.** DeconRNASeq. **C.** dtangle. **D.** Music. *Solid black line*: regression line. *Nmae*: normalised mean absolute error. *r*: Pearson correlation coefficient. *Neurons_Inh and Neurons_Exc*: Inhibitory and excitatory neurons, respectively.

**Supplementary Figure 11. Scatterplots of true and deconvolution-estimated proportions in 100 VL-based *in silico* mixtures**. The signature used all cell-subtypes from the original publication by Velmeshev *et al.* (2019). **A.** CIBERSORT deconvolution. **B.** DeconRNASeq. *Solid black line*: regression line. *Nmae*: normalised mean absolute error. *r*: Pearson correlation coefficient.

**Supplementary Figure 12. Scatterplots of true and deconvolution-estimated proportions in 100 VL-based *in silico* mixtures**. The signature used all cell-subtypes from the original publication by Velmeshev *et al.* (2019). **A.** dtangle deconvolution. **B.** MuSiC. *Solid black line*: regression line. *Nmae*: normalised mean absolute error. *r*: Pearson correlation coefficient.

**Supplementary Figure 13. Heatmap of Spearman correlations between cell-subtypes in the VL dataset.** Labels are taken from the original publication. Numbers in brackets on the right axis labels indicate the number of nuclei in that class.

**Supplementary Figure 14. Effect of cell-type abundance and collinearity on deconvolution accuracy in VL-based simulations.** Each point represents a cell-subtype in the Velmeshev dataset. Points are labelled by text indicating the cell-subtype classification. Colours represent a binary code for good and poor deconvolution performance. *Rho*: Spearman correlation coefficient. *X axis*: mean abundance across the 100 simulated mixtures. *Y axis*: the highest correlation a cell-subtype has to any of the other cell-subtypes in the dataset, indicating collinearity. Note that “o” and “a” are partially overlapping at x=0.5, y = 0.9, as are “k” and “I” at x=15 and y = 0.96.

**Supplementary Figure 15**. Scatterplots of true and deconvolution-estimated proportions in 100 CA-based *in silico* mixtures. The signature used all cell-subtypes from the original publication by Hodge *et al.* (2019). **A.** CIBERSORT deconvolution. **B.** DeconRNASeq. *Solid black line*: regression line. *Nmae*: normalised mean absolute error. *r*: Pearson correlation coefficient.

**Supplementary Figure 16. Scatterplots of true and deconvolution-estimated proportions in 100 CA *in silico* mixtures.** The signature used all cell-subtypes from the original publication by Hodge *et al.* (2019). **A.** dtangle deconvolution. **B.** MuSiC deconvolution. *Solid black line*: regression line. *Nmae*: normalised mean absolute error. *r*: Pearson correlation coefficient.

**Supplementary Figure 17. Heatmap of Spearman correlations between cell-subtypes in the CA dataset**. Labels are taken from the original publication. Numbers in brackets on the right axis labels indicate the number of nuclei in that class.

**Supplementary Figure 18. Effect of cell-type abundance and collinearity on deconvolution accuracy in CA-based simulations.** Each point represents a cell-subtype in the CA dataset. Points are labelled by text indicating the cell-subtype classification. Colours represent a binary code for good and poor deconvolution performance. *Rho*: Spearman correlation coefficient. *X axis*: mean abundance across the 100 simulated mixtures. *Y axis*: the highest correlation a cell-subtype has to any of the other cell-subtypes in the dataset, indicating collinearity. Note that the following labels are partially overlapping: “l”, “g”, and “o” at x=3, y=0.87; and “b”, ”f”, and “h” at x=7, y=0.95

**Supplementary Figure 19. Effect of removing cell-types or cell-subtypes from the signature matrix.** For each cell-type, its mean abundance in the mixtures is shown in backets, and scatterplots display the deconvolution accuracy when all cell types are present in the signature (x-axis) *vs.* when the cell type is absent from the signature (y-axis). Accuracy is measured as either *r* correlation coefficient (left panel) or normalised mean absolute error (right panel). Calculations of mean *r* and mean NMAE for the x-axis label do not include the absent cell-type, and thus differ across plots. *Dotted red line:* y = x.

**Supplementary Figure 20. Effect of varying the signature in VL-based simulated mixtures**. Heatmaps of Pearson correlation (*r*; **A.**) and normalised mean absolute error (nmae; **B.**) for estimated versus true proportion when varying the reference signature. The mixtures are 100 *in silico* VL simulations. Signatures only included a pan-neuronal expression profile, rather than excitatory or inhibitory sub-types. Blank squares indicate that the cell-type was not present in the signature, and thus no statistic was calculated. Grey squares indicate NA, indicating that the cell-type was present in the signature but the statistic could not be calculated; for *r*, this means there was no variance in the composition estimates, typically meaning all 100 samples’ estimates were 0 or 1. For more details about signature characteristics, see methods.

**Supplementary Figure 21. Effect of varying the signature in CA-based simulated mixtures**. Heatmaps of Pearson correlation (*r*; **A.**) and normalised mean absolute error (nmae; **B.**) for estimated versus true proportion when varying the reference signature. The mixtures are 100 *in silico* CA simulations. Signatures only included a pan-neuronal expression profile, rather than excitatory or inhibitory sub-types. Blank squares indicate that the cell-type was not present in the signature, and thus no statistic was calculated. Grey squares indicate NA, indicating that the cell-type was present in the signature but the statistic could not be calculated; for *r*, this means there was no variance in the composition estimates, typically meaning all 100 samples’ estimates were 0 or 1. For more details about signature characteristics, see methods.

**Supplementary Figure 22. Effect of varying the signature while including neuronal subtypes in DM-based simulated mixtures.** Heatmaps of Pearson correlation (*r*; top panel) and normalised mean absolute error (nmae; bottom panel) for estimated versus true proportion when varying the reference signature. Signatures only included a pan-neuronal expression profile, rather than excitatory or inhibitory sub-types. *Neu*: Neurons. *Ast*: Astrocytes. *Oli*: Oligodendrocytes. *Mic*: Microglia. *End*: Endothelia. For more details about signature characteristics, see methods.

**Supplementary Figure 23. Effect of varying the signature while including neuronal subtypes in VL-based simulated mixtures.** Heatmaps of Pearson correlation (*r*; **A.**) and normalised mean absolute error (nmae; **B.**) for estimated versus true proportion when varying the reference signature. The mixtures are 100 *in silico* VL simulations. All signatures here include information about the broad neuronal subtype (excitatory or inhibitory). Blank squares indicate that the cell-type was not present in the signature, and thus no statistic was calculated. Grey squares indicate NA, indicating that the cell-type was present in the signature but the statistic could not be calculated; for *r*, this means there was no variance in the composition estimates, typically meaning all 100 samples’ estimates were 0 or 1. For more details about signature characteristics, see methods.

**Supplementary Figure 24. Effect of varying the signature on CA-based simulated mixtures.** Heatmaps of Pearson correlation (*r*; **A.**) and normalised mean absolute error (nmae; **B.**) for estimated versus true proportion when varying the reference signature. The mixtures are 100 *in silico* CA simulations. All signatures here include information about the broad neuronal subtype (excitatory or inhibitory). Blank squares indicate that the cell-type was not present in the signature, and thus no statistic was calculated. Grey squares indicate NA, indicating that the cell-type was present in the signature but the statistic could not be calculated; for *r*, this means there was no variance in the composition estimates, typically meaning all 100 samples’ estimates were 0 or 1. For more details about signature characteristics, see methods.

**Supplementary Figure 25. The role of compartment-specific genes when using snRNA-seq signatures. A and B:** Estimated proportions for pure samples of immuno-panned cell-types, using whole-cell and snRNA-seq signatures. (A) all genes were included in the signature. (B) compartment-specific genes were filtered-out. *Thick horizontal line*: mean. *Dotted vertical line*: separates whole-cell from snRNA-seq signatures. *Neu*: neurons. *Ast*: astrocytes. *Oli*: oligodendrocytes. *Mic*: microglia. **C.** Scatterplot of estimated proportions for bulk brain samples using the RNA-seq signature, IP (x-axis) or the snRNA-seq signature derived from the same individual (y-axis). *Left*: all genes were included in the signature; *right*: compartment-specific genes were filtered-out. *Individual*: NICHD brain bank id number. Cell-type proportions were estimated using CIBERSORT.

**Supplementary Figure 26. Proportion of variance in gene expression explained by singular-value decomposition.** Linseed proposes that the saturation point of the curve (*i.e.* the number of linearly-independent components that contribute to the mixture) is its number of constituent cell-types. **A-B.** VL-derived mixtures based on random sampling (A) and those simulated with a wide range and variance in cell-type composition (B). **C-D.** CA mixtures based on random sampling (C) and and those simulated with a wide range and variance in cell-type composition (D). **E.** DM mixtures based on random sampling. **F.** RNA mixtures.

**Supplementary Figure 27. Scatterplots of reference-free deconvolution estimates versus true proportion**. **A-C.** Linseed. Plots are only shown for inferred cell-types correlated to a true cell-type at r > 0.5. A. VL-derived random simulations. B. CA-derived random simulations. C. DM-derived random simulations. *Dotted line*: y = x. **D-F.** Coex. Plots are only shown for inferred cell-types that were assigned to a true cell-type through marker enrichment analysis (Fisher Test, p<1×10^-5^, odds ratio > 5; Methods) . D. VL-derived random simulations. E. CA-derived random simulations. F. DM-derived random simulations.

**Supplementary Figure 28.** Scatterplot of proportions estimated by Linseed in the RNA mixtures of neurons and astrocytes setting number of cell types *k*=2. *Black line*: regression line. *Red dotted line*: y=x.

**Supplementary Figure 29.** Scatterplots of reference-free deconvolution estimates versus true proportion in simulations with increased cell-type variance. Simulations were based on VL single-nuclei. **A.** Linseed. **B.** Coex.

**Supplementary Figure 30.** Scatterplots of reference-free deconvolution estimates versus true proportion in simulations with increased cell-type variance. Simulations were based on CA single-nuclei. **A.** Linseed. **B.** Coex.

**Supplementary Figure 31. Interplay between confounds in excitatory cell-type proportion and differential expression for true up- or down-regulated genes**. **A.** Scatterplots show the relationship between the confound in excitatory proportion within a simulation, and the discriminatory ability (fraction of known perturbed genes in the 200 genes with the smallest p-value) (left), the true positive rate for marker genes of excitatory neurons (middle), and the true positive rate for non-marker genes (right). *Top-left*: true fold-change of 1.1. *Top-right*: true fold-change of 1.3. *Bottom-left*: true fold-change of 2. *Coloured lines*: local regression line. *Dotted line in the left* panel: 0.95 times the discriminatory ability for LM when x = 0. **B.** Model robustness to composition confounds. Barplots show the smallest composition decrease where discriminatory ability fell below 0.95 of the baseline (*i.e.* that from an uncorrected linear model on a simulation with no composition confound). Labels are per A.

**Supplementary Figure 32. Cell-type-specific differential expression analysis using CIBERSORTx. A.** Composition distribution of simulated datasets. *Left*: simulated data without a composition difference between the two groups. *Right*: simulated data with a composition difference between the two groups. Each group contained 50 samples. **B.** Gene expression was perturbed 1.5-fold in Group B in inhibitory neurons for 100 non-marker genes plus 100 inhibitory neuronal marker genes. CIBERSORTx was used to extract cell-type-specific expression (Methods), with a linear model then run to assess differential expression in each cell-type. The plot displays the fraction of the true perturbed genes with an FDR < 0.05. Note that the fraction was calculated using only the subset of perturbed genes which were detected in the given cell-type. **C and D.** False positive rate across the different simulations when expression was perturbed in either excitatory neurons (C) or inhibitory neurons (D).

**Supplementary Figure 33. Median goodness-of-fit in large bulk brain RNA-seq datasets across signatures and algorithms. A.** GTEx. **B.** Parikshak *et al..* Each panel aggregates results from samples from a given region. Rows represent signatures, and columns represent algorithms. Within each cell, the number of top is the median goodness-of-fit, while the number in parentheses below is its rank across all algorithm/signature combinations. Colours represent rank, ranging from purple (worst performance and high rank) to yellow (best performance and low rank)

**Supplementary Figure 34. Violin plots of goodness of fit across signatures and regions in GTEx data.** The top, middle, and bottom of the white internal boxes mark the 75th, 50th, and 25th percentiles, respectively. Note that the data presented in the top-left panel (CIB/Cortex) was also shown in Figure 5A.

**Supplementary Figure 35. Violin plots of goodness of fit across signatures and regions in the Parikshak dataset.** The top, middle, and bottom of the white internal boxes mark the 75th, 50th, and 25th percentiles, respectively. Note that the data presented in the top-left panel (CIB/Cortex) was also shown in Figure 5B.

**Supplementary Figure 36. Goodness of fit across signatures in simulated data.** Each point represents one of the hundred mixtures per simulated dataset. **A.** VL-based simulation. **B.** CA-based simulation. **C.** DM-based simulation. Note that the order of signatures along the x-axis is based on median goodness-of-fit in that panel, and therefore differs between panels.

**Supplementary Figure 37. Violin plots of goodness-of-fit in two non-brain tissues in the GTEx dataset**. **A.** Pancreas samples. **B.** Heart left ventricle samples. **C.** Heart atrial appendage samples. *Fresh:* signature derived from freshly-processed human tissue. *Cultured:* signature derived from cultured cells. The bottom, middle, and top of the white boxes mark the first, second, and third quantiles, respectively.

**Supplementary Figure 38. Heatmaps of Spearman correlations across signatures**. **A.** Across nine brain signatures. *Top left*: Neurons. *Top middle*: Astrocytes. *Top right*: legend. *Bottom left*: Oligodendrocytes. *Bottom middle*: Microglia. *Bottom right*: Endothelia. **B.** Across four pancreas signatures. *Left*: alpha cells. R*ight*: beta cells. Legend is per the top right panel of A. **C.** Across three heart signatures. *From left to right*: cardiomyocytes, smooth muscle cells, fibroblasts, and endothelial cells. Legend is per the top right panel of A.

**Supplementary Figure 39. Heatmaps of intersection between the top 100 cell-type marker genes across signatures**. **A.** Across nine brain signatures. **B.** Across four pancreas signatures. Legend is per the bottom right panel of A. **C.** Across three heart signatures. *CM*: cardiomyocytes. *SMC*: smooth muscle cells. *FB*: fibroblasts. *EC*: endothelial cells. Legend is per the bottom right panel of A.

**Supplementary Figure 40. tSNE dimensionality reduction plot of snRNA-seq data generated as part of the present study**. Nuclei were annotated using the SingleR package to transfer labels from the NG signature. *Ast*: astrocytes. *End*: endothelia. *Exc*: excitatory neurons. *Inh*: inhibitory neurons. *Oli*: oligodendrocytes. *OPC*: oligodendrocyte precursor cells.

**Supplementary Figure 41. Distribution of gene expression values in real and simulated brain mixtures.** *Brain*: bulk brain RNA-seq from Parikshak et al. (2016). *Simulations*: *in silico* mixtures simulated from the corresponding dataset. *DM*: scRNA-seq from Darmanis *et al.*. *CA*: snRNA-seq data from the Human Cell Atlas. *VL*: snRNA-seq data from Velmeshev *et al.* (2019). Simulations contained 500 cells/mixtures for CA and VL, and 100 cells/mixture for DM. *Nuclei, Cells*: single-nuclei and single-cells from the corresponding dataset. Ten samples were randomly for each plot to minimise overplotting. Note: DM simulation is in units of RPKM rather than CPM.

**Supplementary Figure 42. Heatmap of Pearson correlations for cell-ty[e proportion in samples used for ASD analyses**. Proportions were estimated using CIBERSORT and the Multibrain signature.

**Supplementary Figure 43. The relationship between deconvolution estimates, RNA proportions (pRNA) and cell-type proportion (pCt). A.** Boxplots of RNA content per cell, reflected in the number of unique molecular identifiers (UMIs) per-cell across cell types in the VL dataset. *Ast:* astrocytes. *End:* Endothelia. *Exc:* Excitatory Neurons. Inh: Inhibitory Neurons. *Mic*: Microglia. *Oli*: Oligodendrocytes. *OPC*: Oligodendrocyte Precursor Cells. **B.** Scatterplot of pCt vs. pRNA in pseudo-bulk samples from 10 individuals. **C.** Scatterplot of Estimated proportion (y-axis) versus true pCt (left) or pRNA (right)**. D.** Scatterplot of goodness-of-fit when reconstructing gene expression using pCt or pRNA. *Dotted black lines:* y=x

**Supplementary Figure 44. Heatmap of correlations in neuronal estimates across signatures and algorithms in bulk brain datasets. A.** GTEx. **B.** Parikshak *et al.*. Black squares represent NA, where the cell-type estimate had a variance of 0 (typically all estimates being all 0 or all 1). *Dotted black line:* y=0.5.

**Supplementary Figure 45. Heatmap of correlations in astrocyte estimates across signatures and algorithms in bulk brain datasets. A.** GTEx. **B.** Parikshak *et al.*. Black squares represent NA, where the cell-type estimate had a variance of 0 (typically all estimates being all 0 or all 1). *Dotted black line:* y=0.5.

**Supplementary Figure 46. Heatmap of correlations in oligodendrocyte estimates across signatures and algorithms in bulk brain datasets. A.** GTEx. **B.** Parikshak *et al.*. Black squares represent NA, where the cell-type estimate had a variance of 0 (typically all estimates being all 0 or all 1). *Dotted black line:* y=0.5.

**Supplementary Figure 47. Heatmap of correlations in microglial estimates across signatures and algorithms in bulk brain datasets. A.** GTEx. **B.** Parikshak *et al.*. Black squares represent NA, where the cell-type estimate had a variance of 0 (typically all estimates being all 0 or all 1). *Dotted black line:* y=0.5.

**Supplementary Figure 48. Heatmap of correlations in endothelial estimates across signatures and algorithms in bulk brain datasets. A.** GTEx. **B.** Parikshak *et al.*. Black squares represent NA, where the cell-type estimate had a variance of 0 (typically all estimates being all 0 or all 1). *Dotted black line:* y=0.5.

**Supplementary Figure 49. Distribution of neuronal deconvolutions estimates in bulk brain datasets. A.** GTEx. **B.** Parikshak *et al.*. The top, middle, and bottom of the white internal boxes mark the 75th, 50th, and 25th percentiles, respectively. *Dotted black line:* y=0.5.

**Supplementary Figure 50. Distribution of astrocyte deconvolutions estimates in bulk brain datasets. A.** GTEx. **B.** Parikshak *et al.*. The top, middle, and bottom of the white internal boxes mark the 75th, 50th, and 25th percentiles, respectively. *Dotted black line:* y=0.5.

**Supplementary Figure 51. Distribution of oligodendrocyte deconvolutions estimates in bulk brain datasets. A.** GTEx. **B.** Parikshak *et al.*. The top, middle, and bottom of the white internal boxes mark the 75th, 50th, and 25th percentiles, respectively. *Dotted black line:* y=0.5.

**Supplementary Figure 52. Distribution of microglial deconvolutions estimates in bulk brain datasets. A.** GTEx. **B.** Parikshak *et al.*. The top, middle, and bottom of the white internal boxes mark the 75th, 50th, and 25th percentiles, respectively. *Dotted black line:* y=0.5.

**Supplementary Figure 53. Distribution of endothelial deconvolutions estimates in bulk brain datasets. A.** GTEx. **B.** Parikshak *et al.*. The top, middle, and bottom of the white internal boxes mark the 75th, 50th, and 25th percentiles, respectively. *Dotted black line:* y=0.5.

## Tables

**Table 1.** Description of algorithms benchmarked in this study. *: For brevity, DeconRNASeq, CIBERSORT, and BrainInABlender will be referred to in-text as DRS, CIB, and Blender, respectively. **: the identities of unlabeled cell-types were inferred through cell-type marker enrichment (Methods)

## Supplementary Tables

**Supplementary Table 1.** Differential expression analysis comparing nuclear and whole-cell brain tissue preparations.

**Supplementary Table 2.** Composition estimates in pure immunopanned brain cell-types using either all genes or stable genes. All signatures contained Neurons, Astrocytes, Oligodendrocytes, and Microglia. The estimates shown are those for the corresponding pure cell-type only. The column name shows the algorithm and signature combination used. *Above_0.8*: percentage of samples in which the composition estimate is > 0.8.

**Supplementary Table 3.** Composition estimates in GTEx and Parikshak *et al.* bulk brain transcriptomes, across signatures and methods

**Supplementary Table 4.** Differentially expression analysis results for ASD samples vs. controls for composition-dependent (CD) and composition-independent (CI) analyses. DEGs: genes significant at FDR< 0.05. GO: gene ontology terms significant at FDR< 0.05.

**Supplementary Table 5.** List of datasets accessed and the samples included in the present study from each dataset.

**Supplementary Table 6.** Cell-type specific gene expression signature data. Expression values are normalised and filtered as described in Methods. Each tab shows expression for a different signature.

**Supplementary Table 7.** Summary of RNA-seq data generated from mixtures of RNA from cultured cells.

**Supplementary Table 8.** Summary of RNA-seq and snRNA-seq data generated for nuclear versus whole-cell comparisons.

