## Supplementary Figures for "Comprehensive evaluation of deconvolution methods for human brain gene expression"

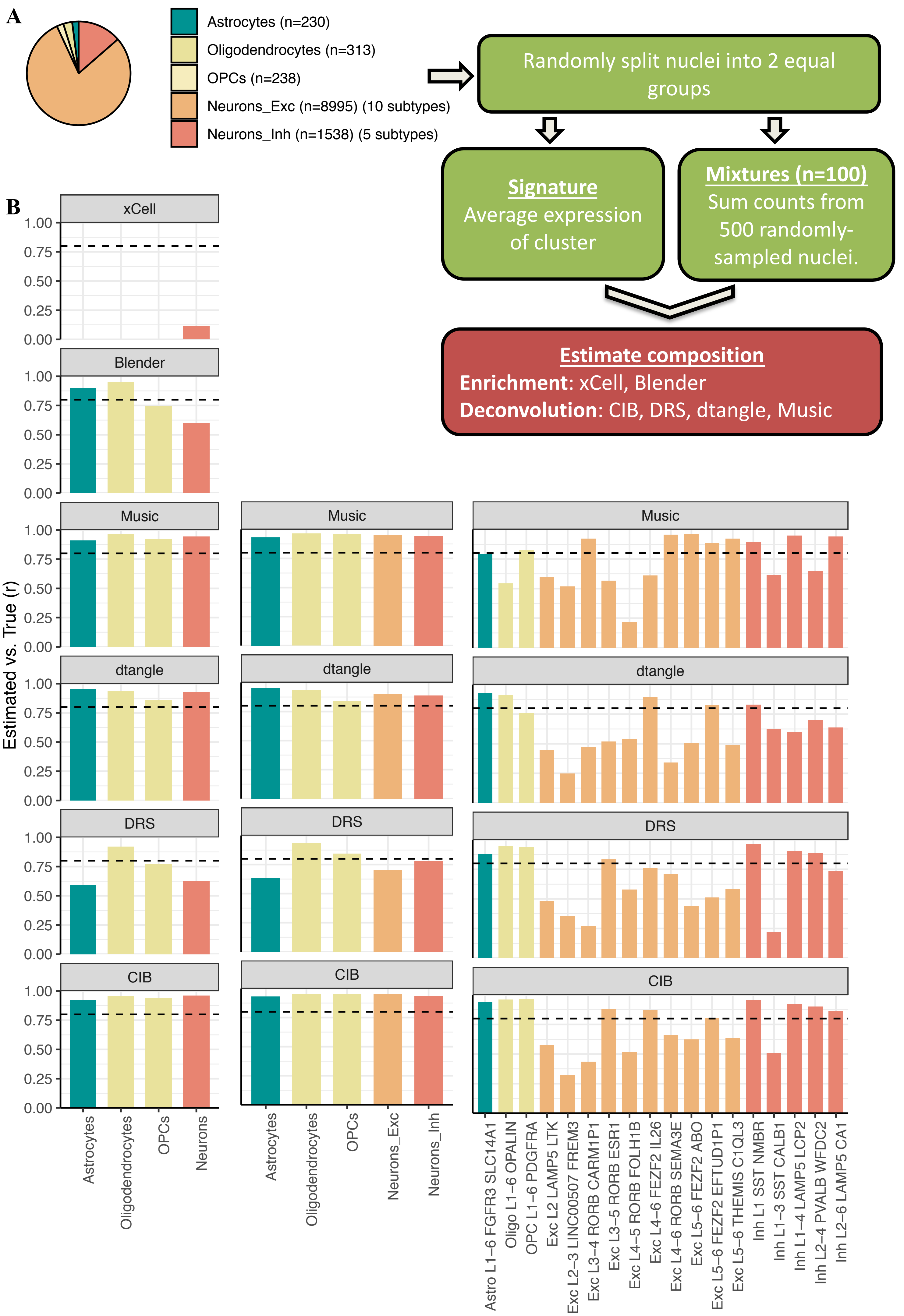


**Supplementary Figure 1. Generation and deconvolution of a replication simulated dataset using single-nuclei from the CA dataset**. **A.** Process for generating 100 in silico mixtures. **B.** Barplots show Pearson correlation coefficients (r) between true and deconvolution-estimated proportions in 100 *in silico* mixtures. The left column shows results when only major cell-type labels are used in the signature; the middle column shows results when a mix of major cell-type and cell-subtype labels are used in the signature; the right column shows results when only cell-subtype labels are used in the signature. *Dotted line*: *r*=0.8.

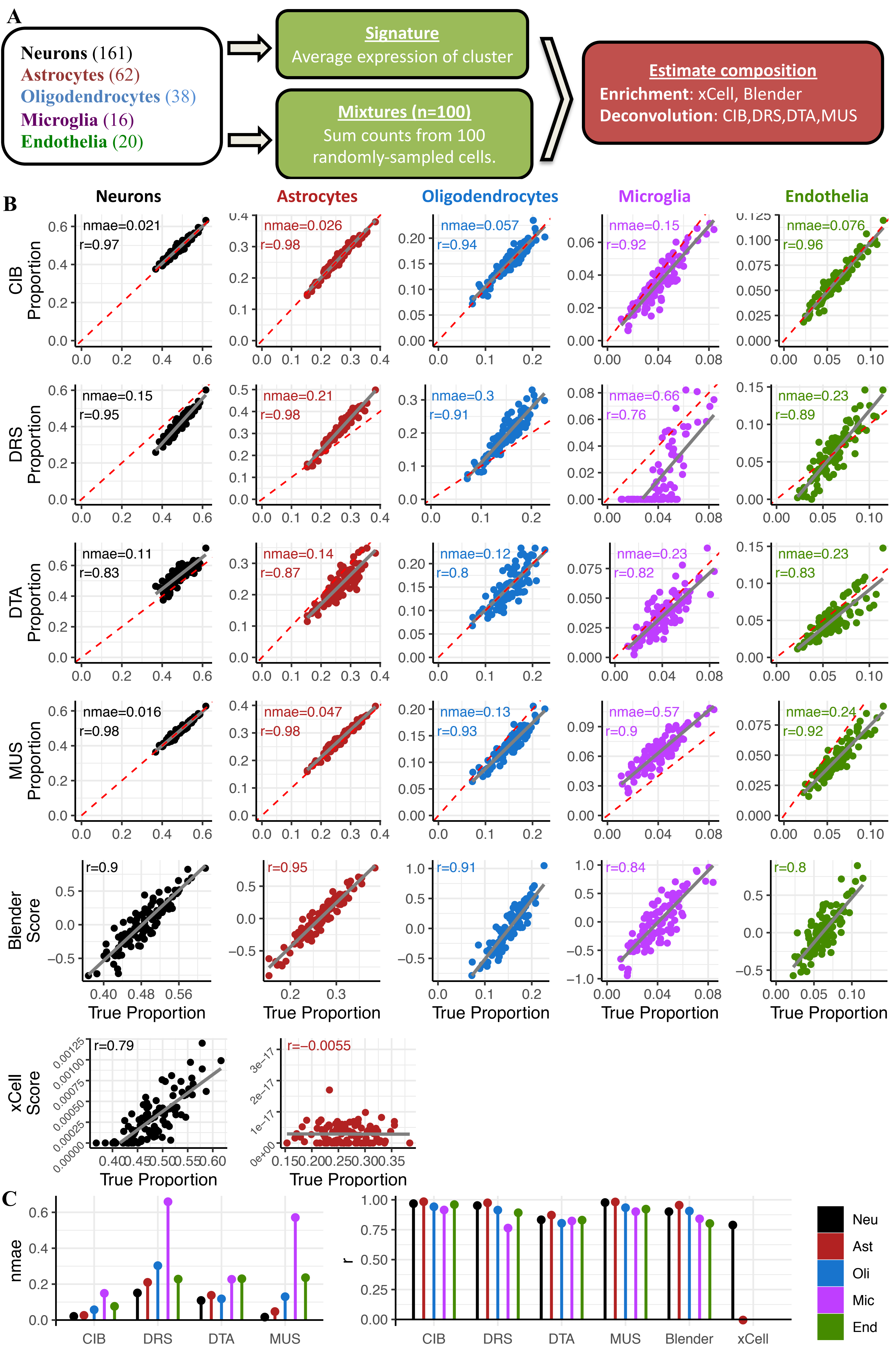


**Supplementary Figure 2. Benchmarking deconvolution algorithms on simulated mixtures using data from Darmanis *et al.*** **A.** Simulation design. **B.** Scatterplots of estimated proportion (or enrichment score) and true proportion for each cell-type. *Red dotted line*: y=x. Grey line: regression line. **C.** Barplots of normalised mean absolute error *(nmae*; left) and Pearson correlation coefficients between true and estimated proportions (*r*; right) based on 100 *in silico* mixtures.

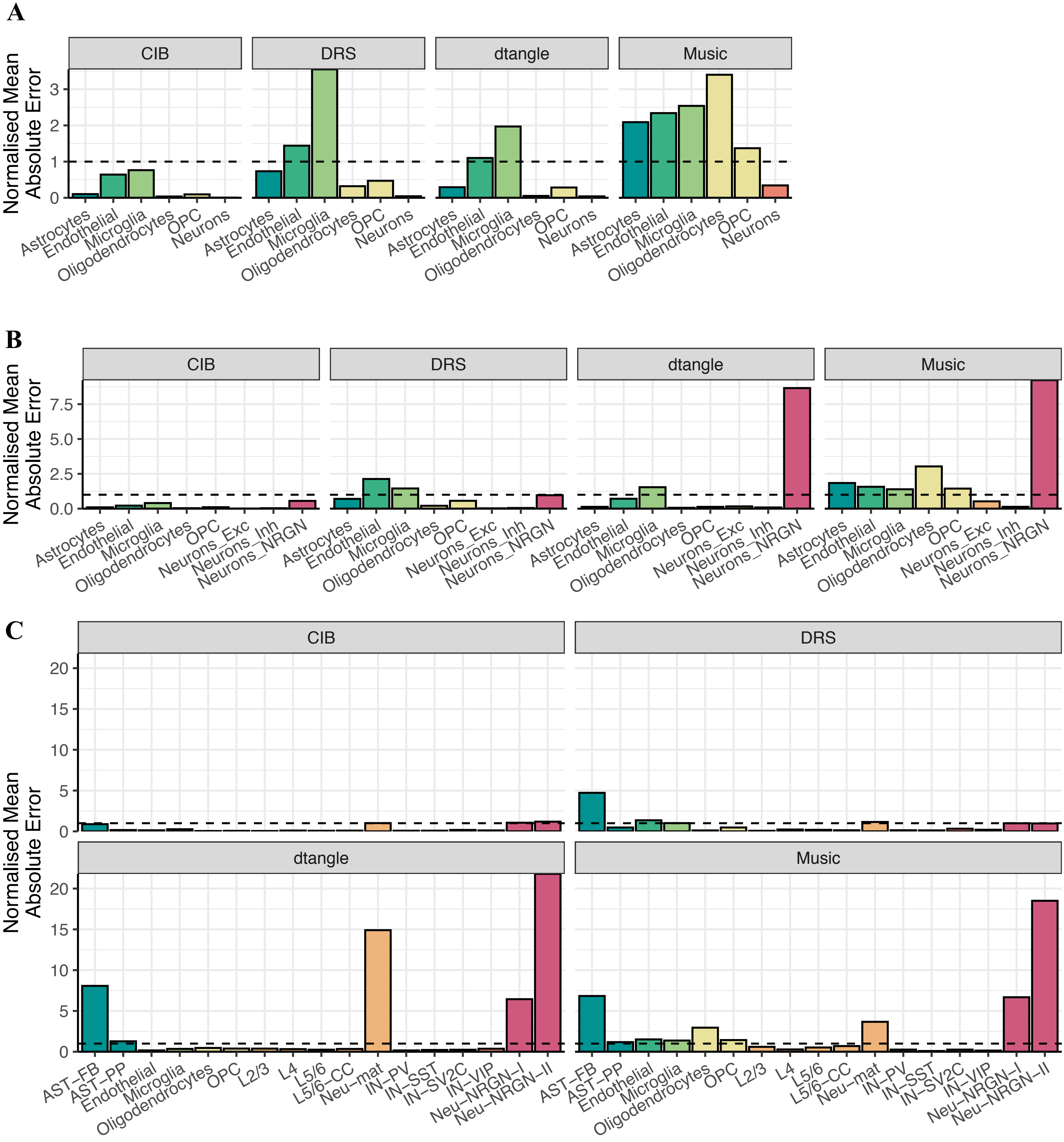


**Supplementary Figure 3.** Barplots of normalised mean absolute error (*nmae*) between true and deconvolution-estimated proportions in 100 VL-derived *in silico* mixtures. *Nmae* was calculated as the average error divided by the average true value. **A.** Using only major cell-type labels in the signature. **B.** Using a mix of major cell-type and cell-subtype labels in the signature. **C.** Using all cell-subtype labels are the signature. *Dotted black line*: *nmae* = 1.

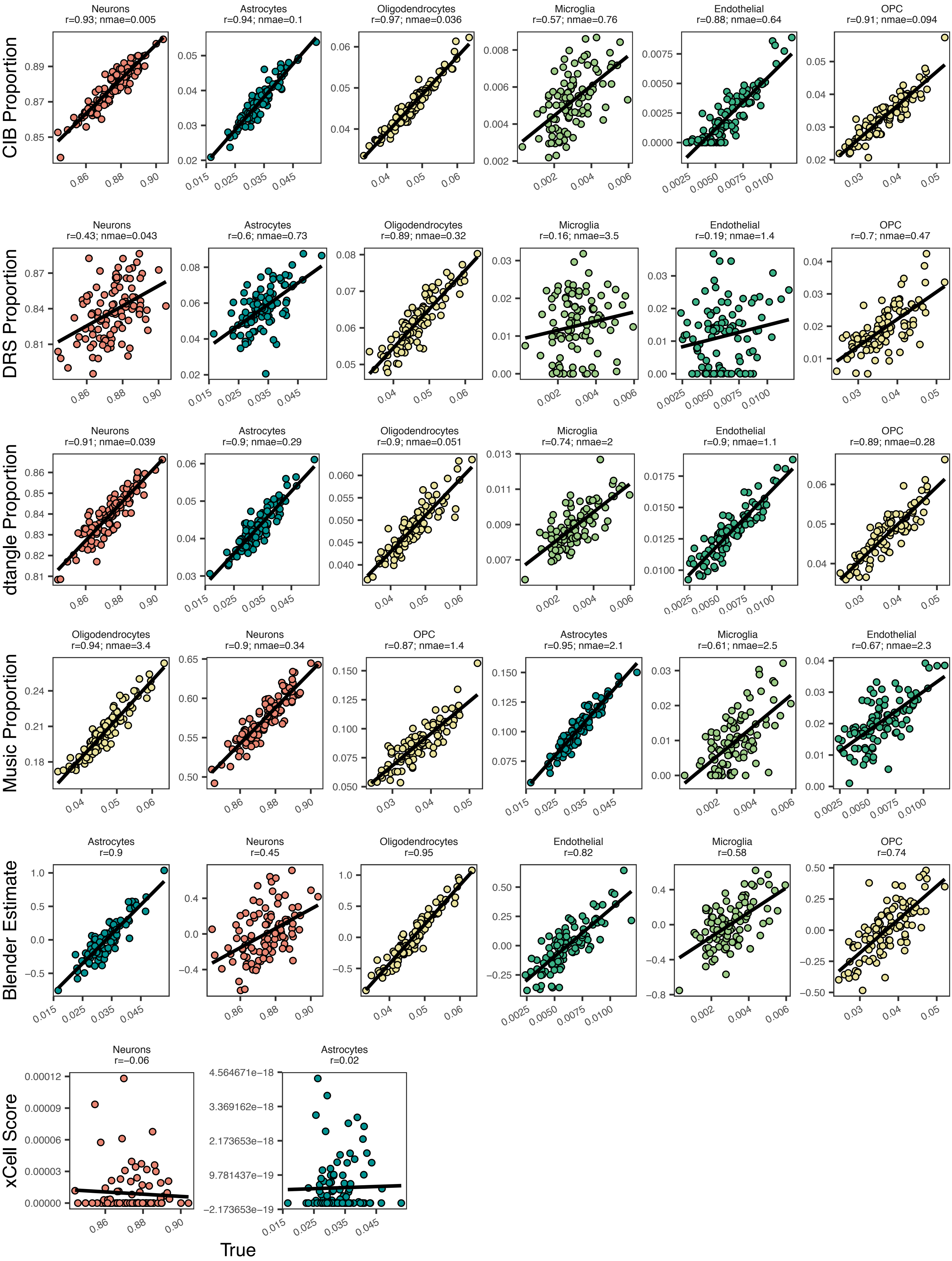


**Supplementary Figure 4.** Scatterplots of true and deconvolution-estimated proportions in 100 VL-derived *in silico* mixtures. Each row represents a different algorithm. The signature used only major cell-types. *Solid black line*: regression line. *Nmae*: normalised mean absolute error. *r*: Pearson correlation coefficient. Note that nmae was not calculated for xCell and Blender as their output is an enrichment score rather than a proportion

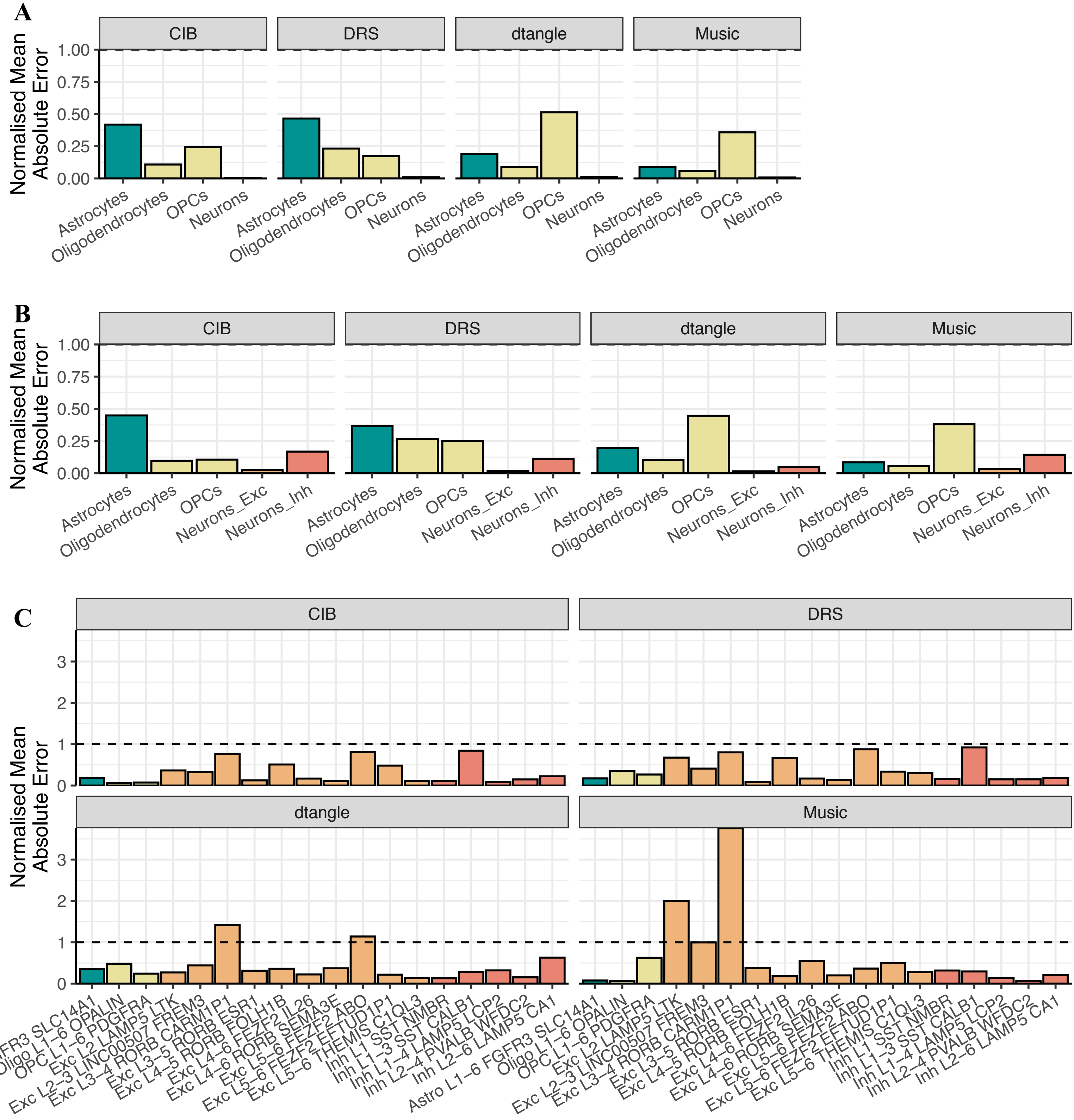


**Supplementary Figure 5.** Barplots of normalised mean absolute error (*nmae*) between true and deconvolution-estimated proportions in 100 CA-derived *in silico* mixtures. *Nmae* was calculated as the average error divided by the average true value. **A.** Using only major cell-type labels in the signature. **B.** Using a mix of major cell-type and cell-subtype labels in the signature. **C.** Using all cell-subtype labels are the signature. *Dotted black line*: *nmae* = 1.

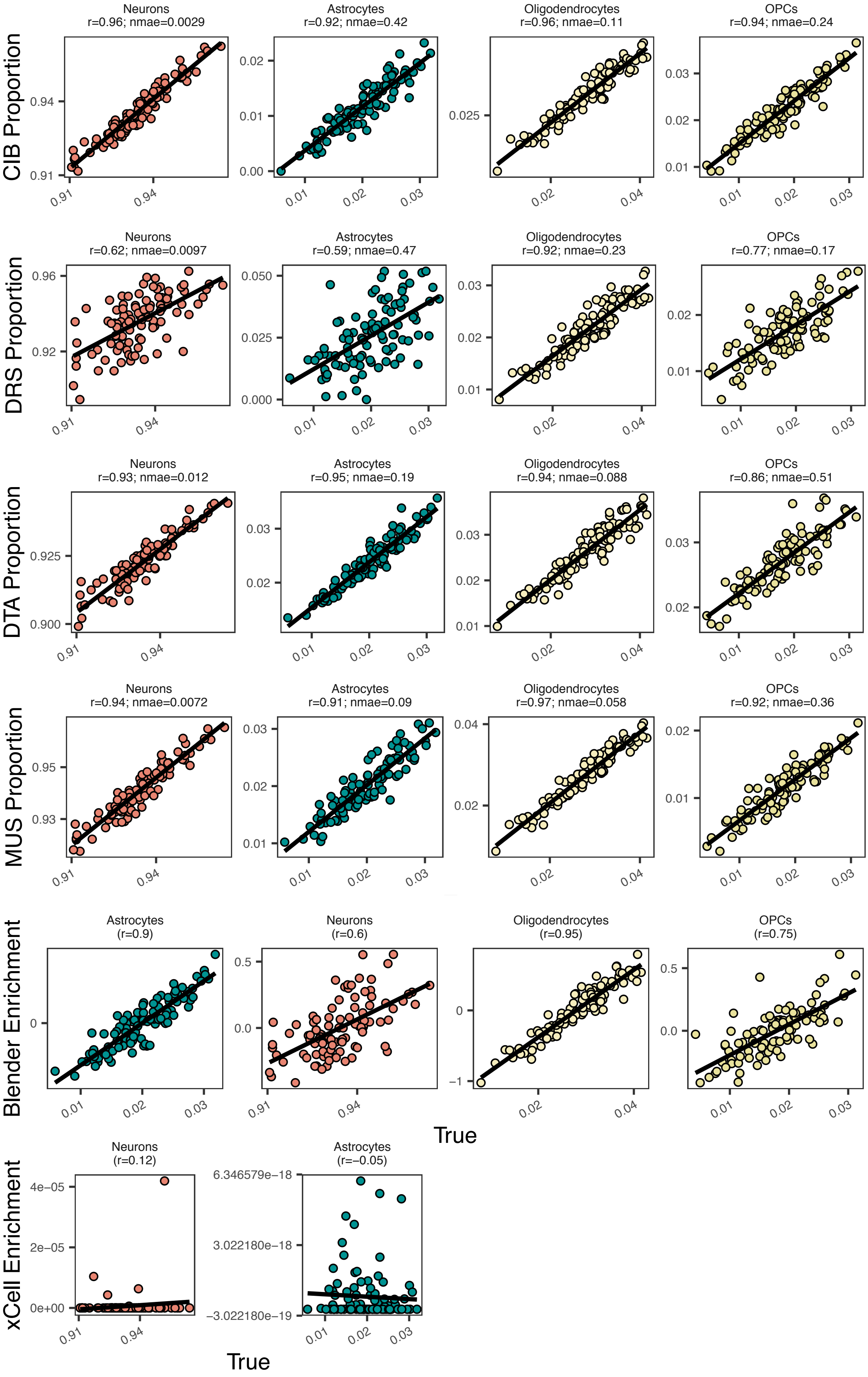


**Supplementary Figure 6.** Scatterplots of true and deconvolution-estimated proportions in 100 CA-derived *in silico* mixtures. Each row represents a different algorithm. The signature used only major cell-types. *Solid black line*: regression line. *Nmae*: normalised mean absolute error. *r*: Pearson correlation coefficient. Note that nmae was not calculated for xCell and Blender as their output is an enrichment score rather than a proportion

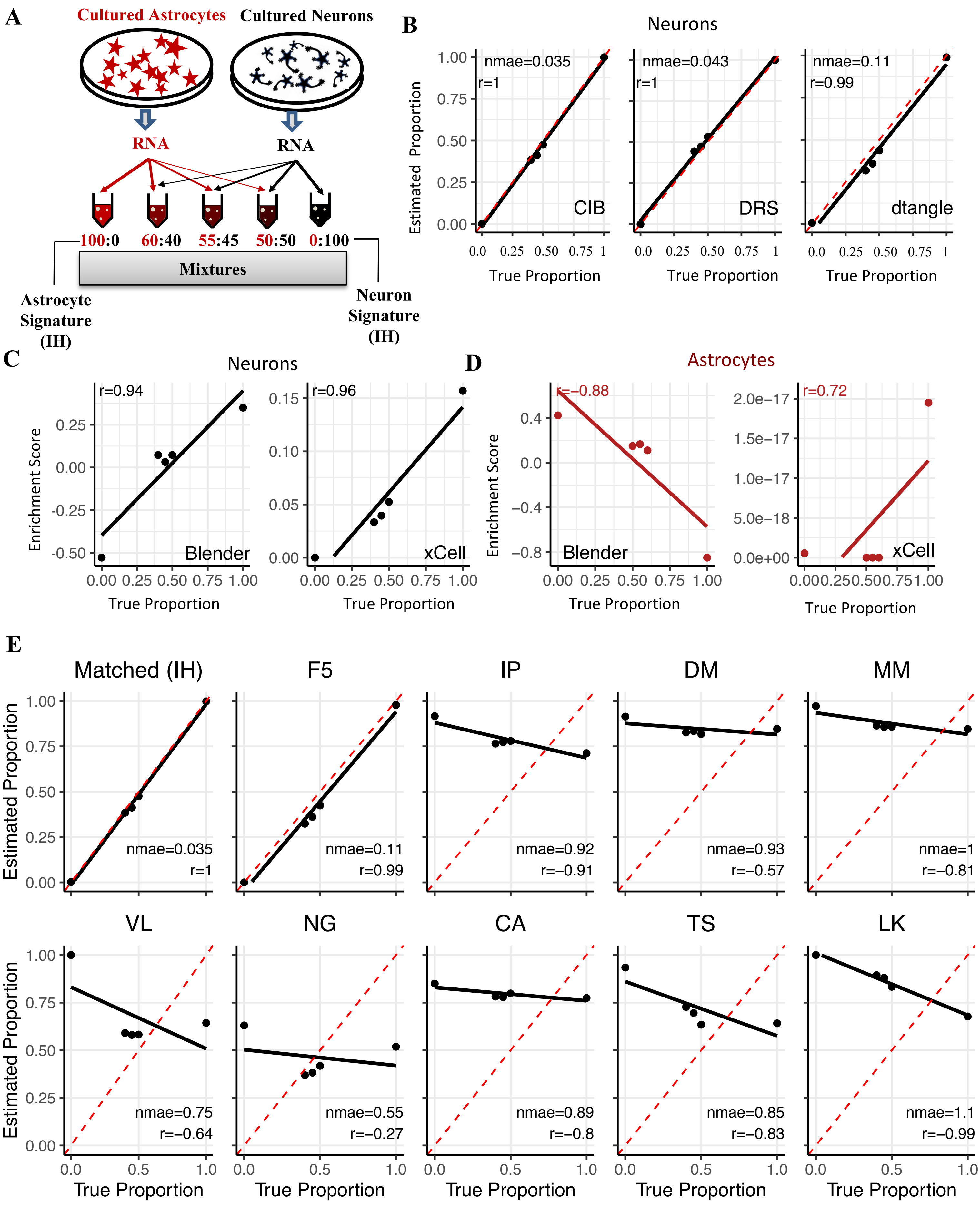


**Supplementary Figure 7. Deconvolving mixtures of RNA from cultured neurons and astrocytes. A.** Outline of RNA mixtures and its corresponding in-house (IH) signature. **B.** Scatterplots of estimated and true proportions of neurons using CIB, DRS and DTA, combined with the matching IH signature. Note that the MUS algorithm was not used, as the algorithm is only compatible with single-cell-level data. **C.** Scatterplots of neuron enrichment scores obtained with Blender (left) and xCell (right). **D.** Scatterplots of astrocyte enrichment scores obtained with Blender (left) and xCell (right). **E.** Scatterplots of true versus estimated neuronal proportion when using CIB and mismatched signatures. All signatures contained just neuronal and astrocyte expression values.

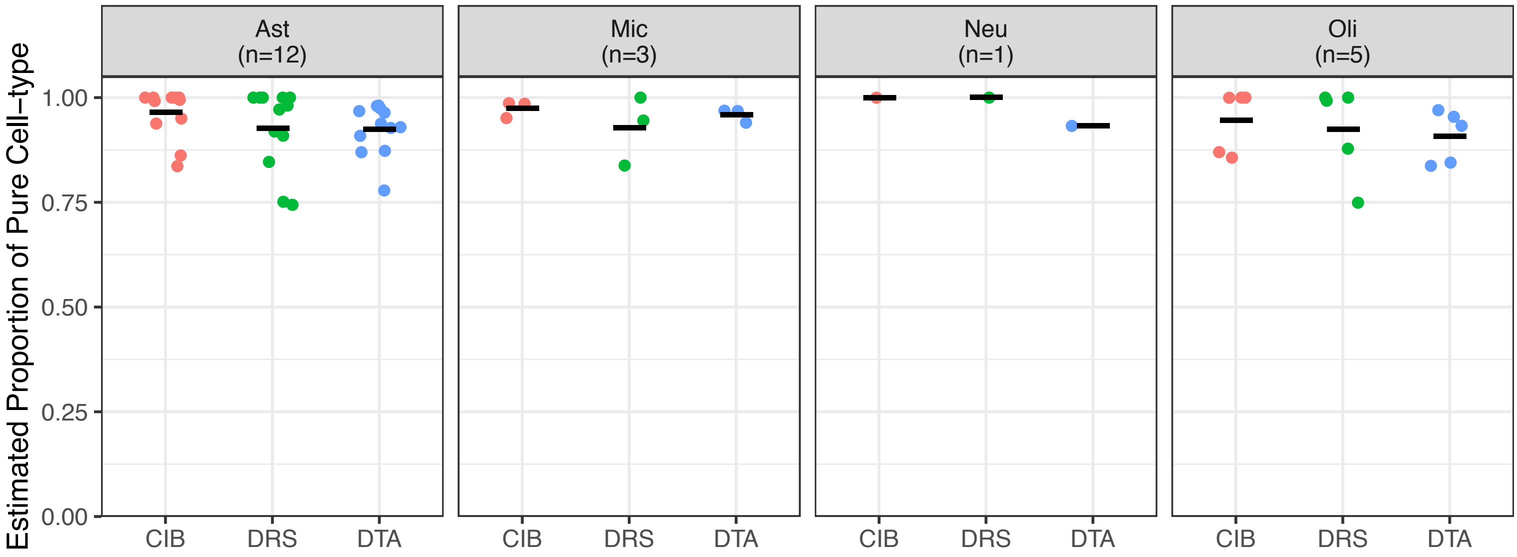


**Supplementary Figure 8.** Estimated proportion in immuno-panned purified brain. *Thick horizontal line*: mean. *Neu*: neurons. *Ast*: astrocytes. *Oli*: oligodendrocytes. *Mic*: microglia. *CIB*: CIBERSORT. *DRS*: DeconRNASeq. *DTA*: dtangle. Note that MuSiC was not applied as it requires single-cell or –nucleus data for its signature.

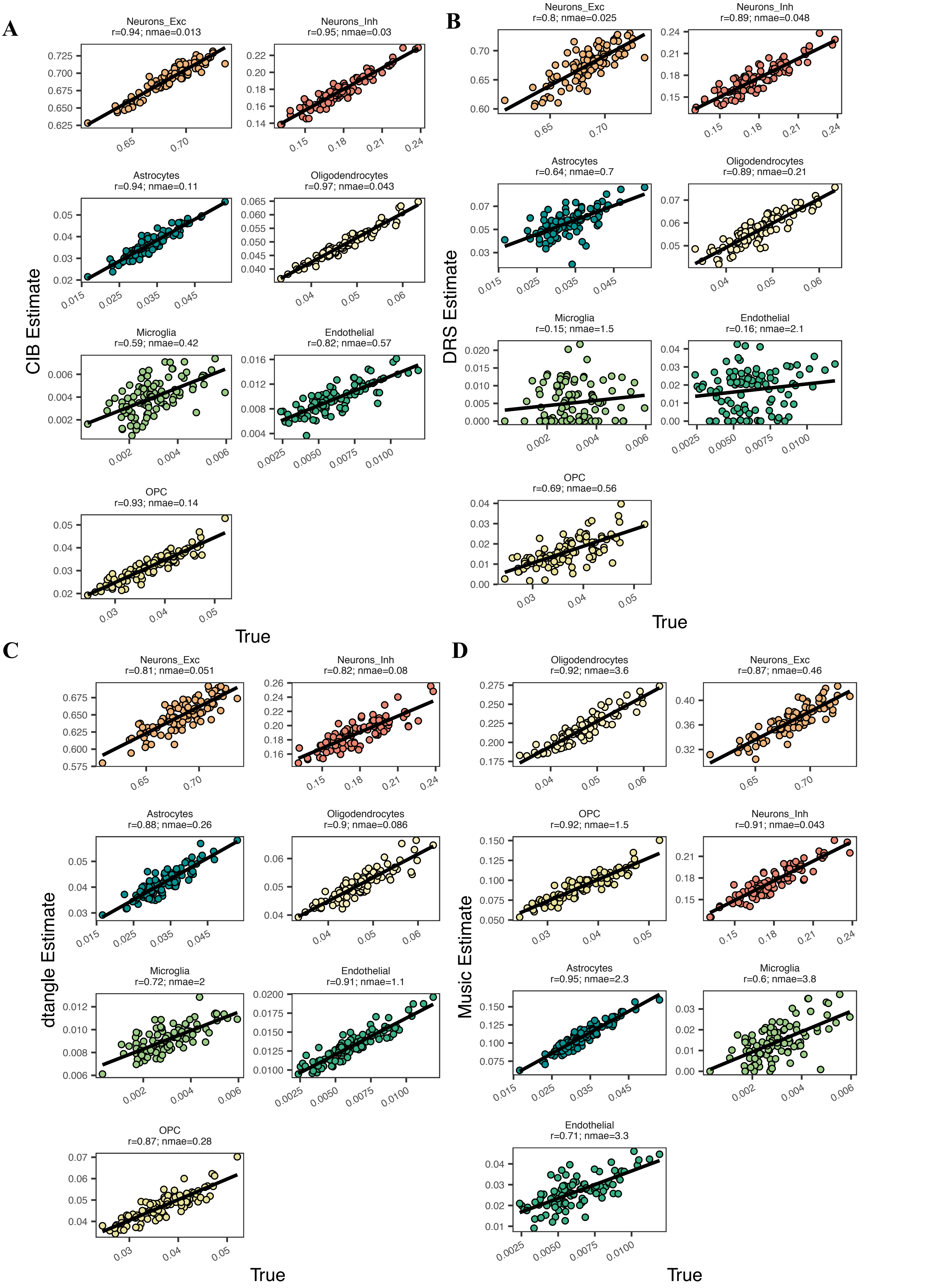


**Supplementary Figure 9. Scatterplots of true and deconvolution-estimated proportions in 100 VL-derived *in silico* mixtures**. The signature used a range of cell-subtypes and major cell-types. **A.** CIBERSORT deconvolution. **B.** DeconRNASeq. **C.** dtangle. **D.** MuSiC. *Solid black line*: regression line. *Nmae*: normalised mean absolute error. *r*: Pearson correlation coefficient. *Neurons_Inh and Neurons_Exc*: Inhibitory and excitatory neurons, respectively.

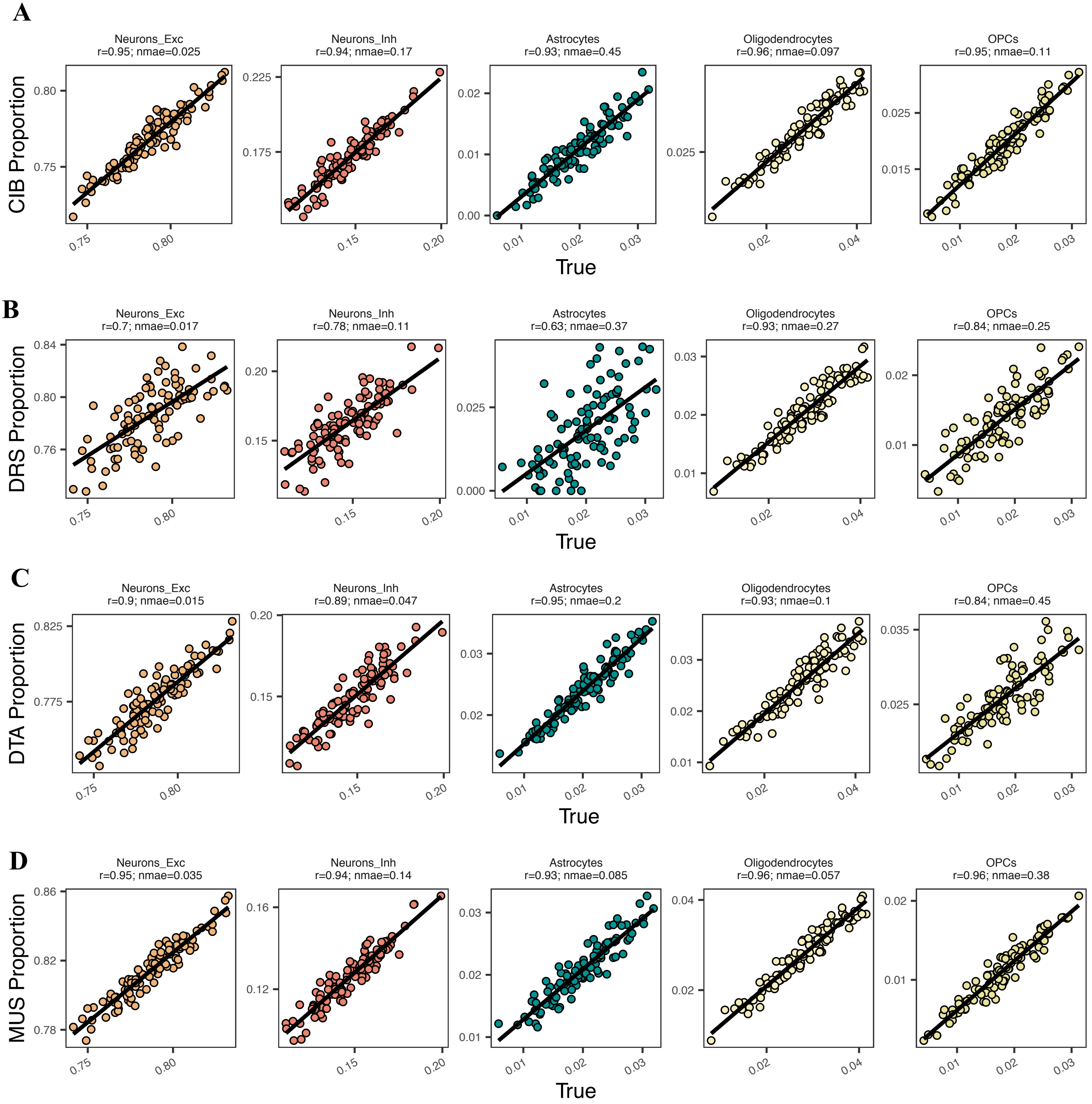


**Supplementary Figure 10.** Scatterplots of true and deconvolution-estimated proportions in 100 CA-*in silico* mixtures. Each row represents a different algorithm. The signature used a range of cell-subtypes and major cell-types. **A.** CIBERSORT deconvolution. **B.** DeconRNASeq. **C.** dtangle. **D.** MuSiC. *Solid black line*: regression line. *Nmae*: normalised mean absolute error. *r*: Pearson correlation coefficient. *Neurons_Inh and Neurons_Exc*: Inhibitory and excitatory neurons, respectively.

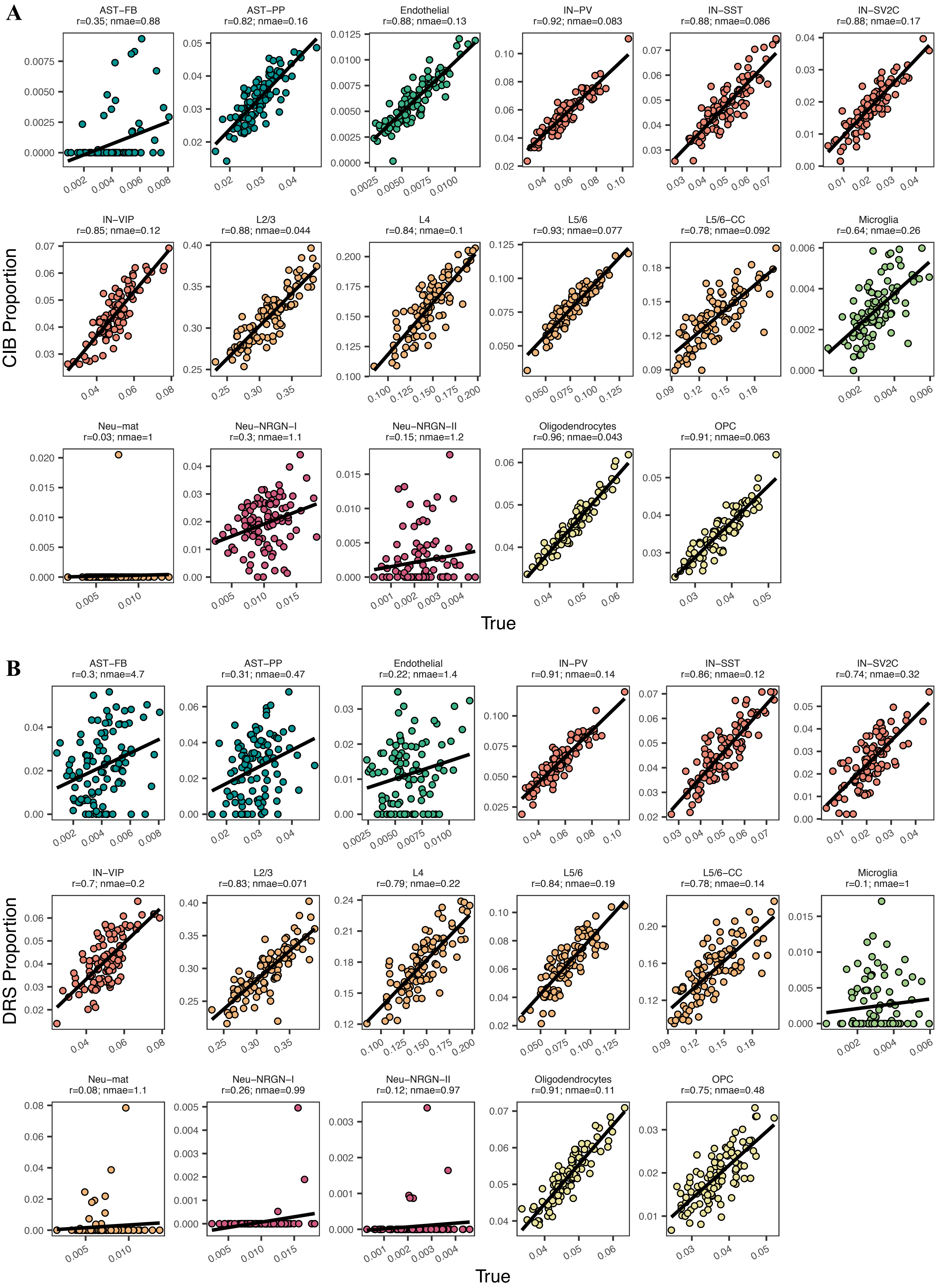


**Supplementary Figure 11. Scatterplots of true and deconvolution-estimated proportions in 100 VL-based *in silico* mixtures.** The signature used all cell-subtypes from the original publication by Velmeshev *et al.* (2019). **A.** CIBERSORT deconvolution. **B.** DeconRNASeq. *Solid black line*: regression line. *Nmae*: normalised mean absolute error. *r*: Pearson correlation coefficient.

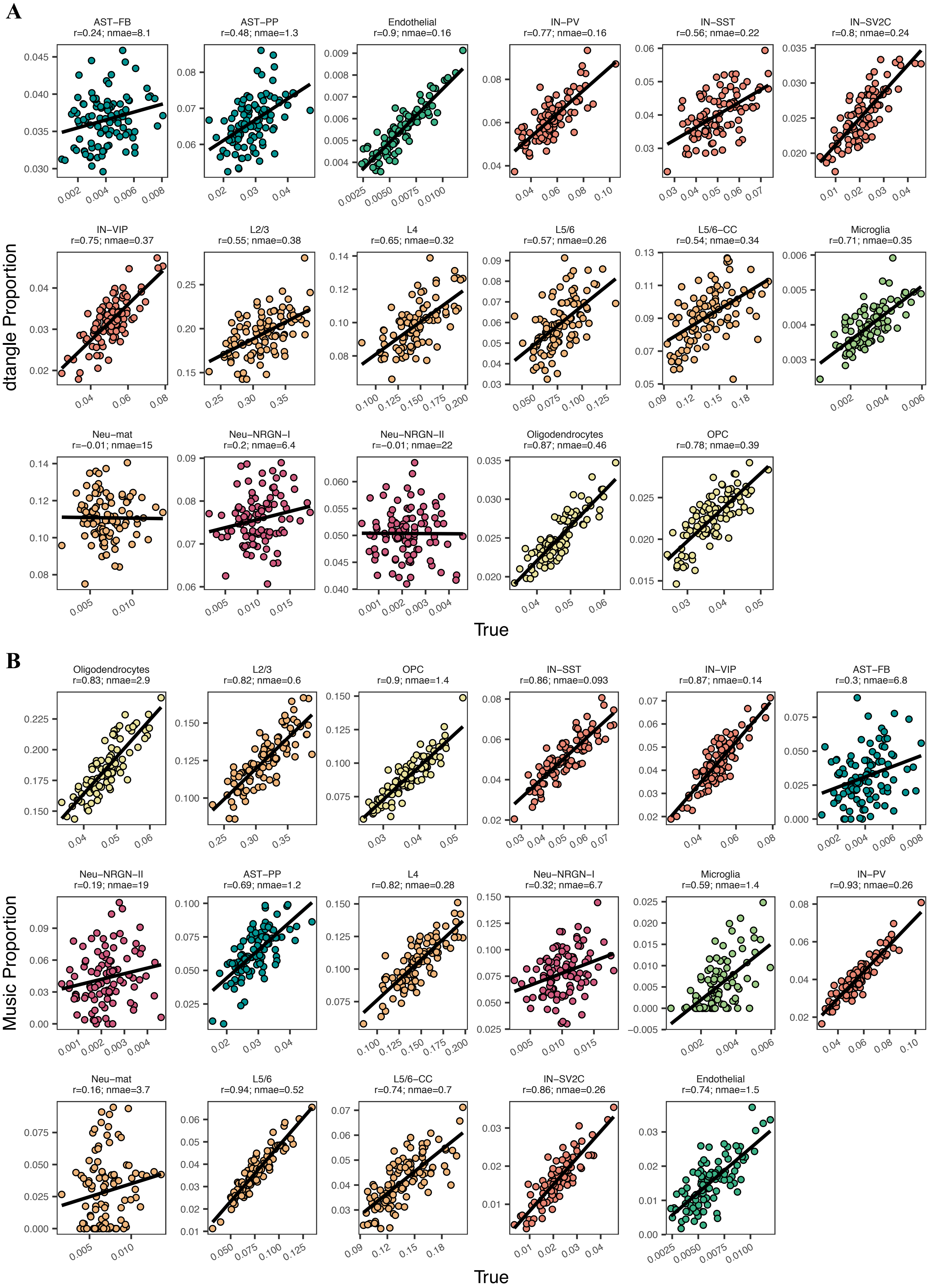


**Supplementary Figure 12. Scatterplots of true and deconvolution-estimated proportions in 100 VL-based *in silico* mixtures.** The signature used all cell-subtypes from the original publication by Velmeshev *et al.* (2019). **A.** dtangle deconvolution. **B.** MuSiC. *Solid black line*: regression line. *Nmae*: normalised mean absolute error. *r*: Pearson correlation coefficient.

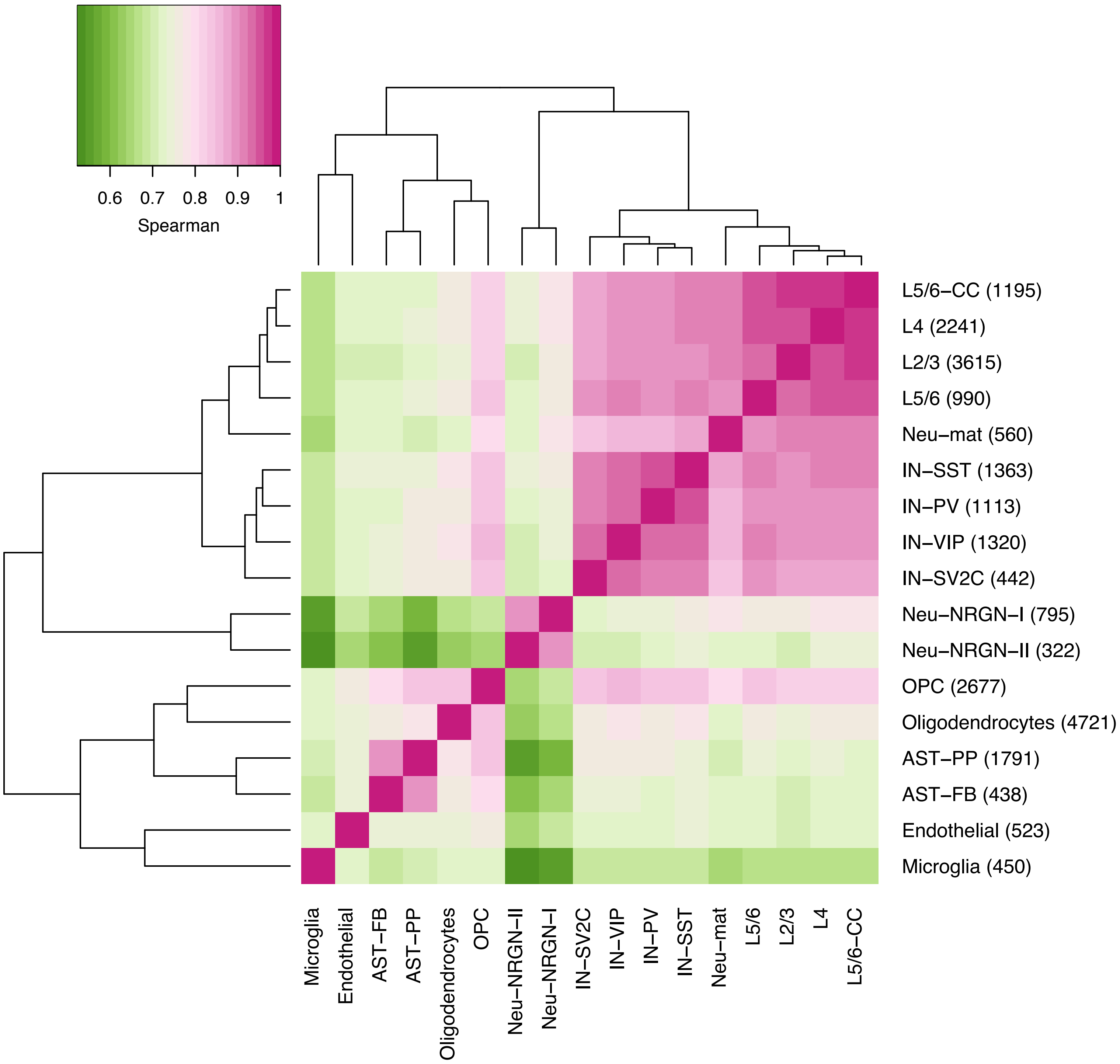


**Supplementary Figure 13. Heatmap of Spearman correlations between cell-subtypes in the VL dataset.** Labels are taken from the original publication**.** Numbers in brackets on the right axis labels indicate the number of nuclei in that class.

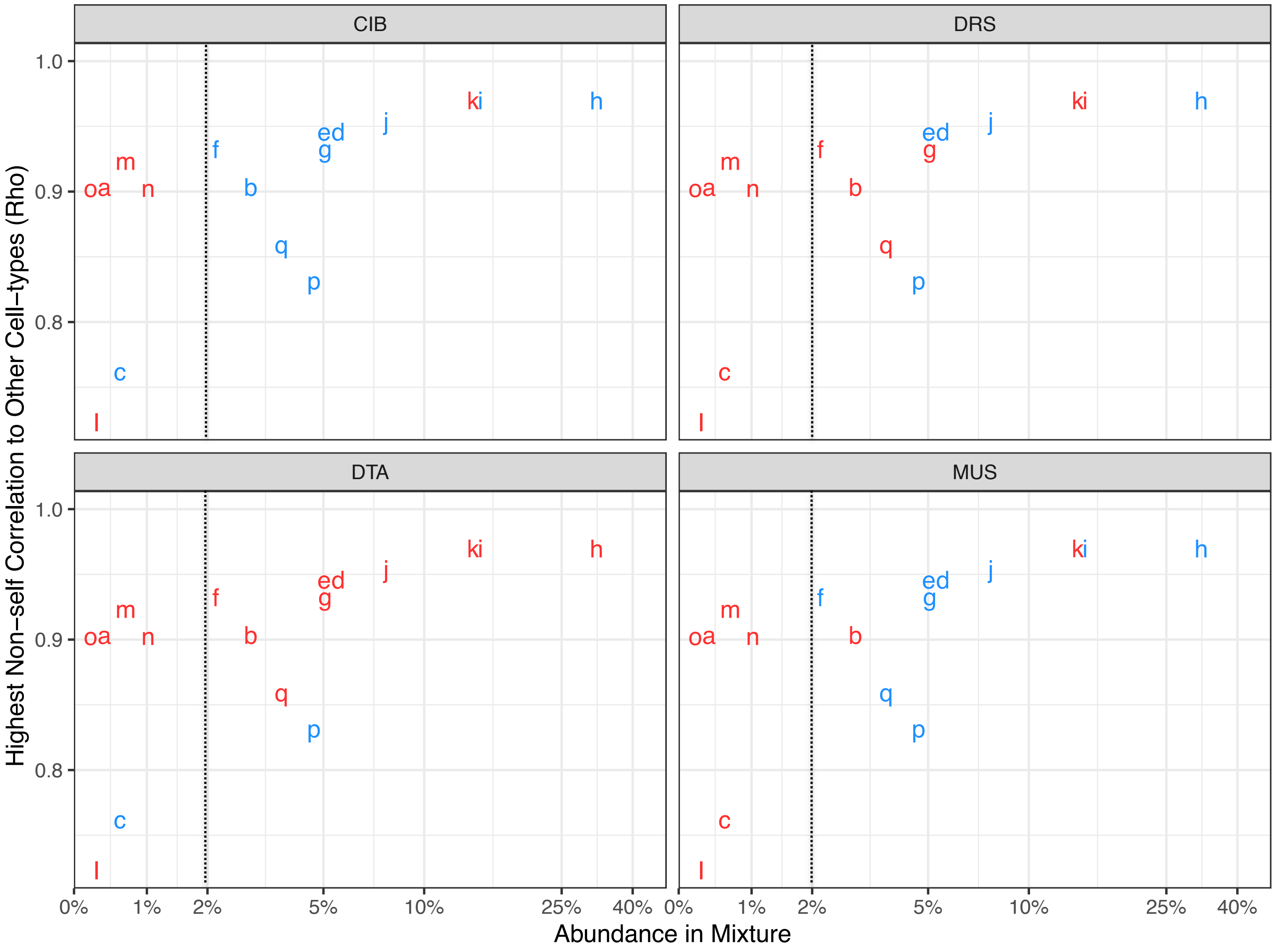


| Label | Celltype |
|---|---|
| a | AST-FB |
| b | AST-PP |
| c | Endothelial |
| d | IN-PV |
| e | IN-SST |
| f | IN-SV2C |
| g | IN-VIP |
| h | L2/3 |
| i | L4 |
| j | L5/6 |
| k | L5/6-CC |
| l | Microglia |
| m | Neu-mat |
| n | Neu-NRGN-I |
| o | Neu-NRGN-II |
| p | Oligodendrocytes |
| q | OPC |

**Deconvolution accuracy**

- Poor (r < 0.8)
- Good (r > 0.8)

**Supplementary Figure 14. Effect of cell-type abundance and collinearity on deconvolution accuracy in VL-based simulations.** Each point represents a cell-subtype in the Velmeshev dataset. Points are labelled by text indicating the cell-subtype classification. Colours represent a binary code for good and poor deconvolution performance. *Rho*: Spearman correlation coefficient. *X axis*: mean abundance across the 100 simulated mixtures. *Y axis*: the highest correlation a cell-subtype has to any of the other cell-subtypes in the dataset, indicating collinearity. Note that “o” and “a” are partially overlapping at x=0.5, y = 0.9, as are “k” and “I” at x=15 and y = 0.96.

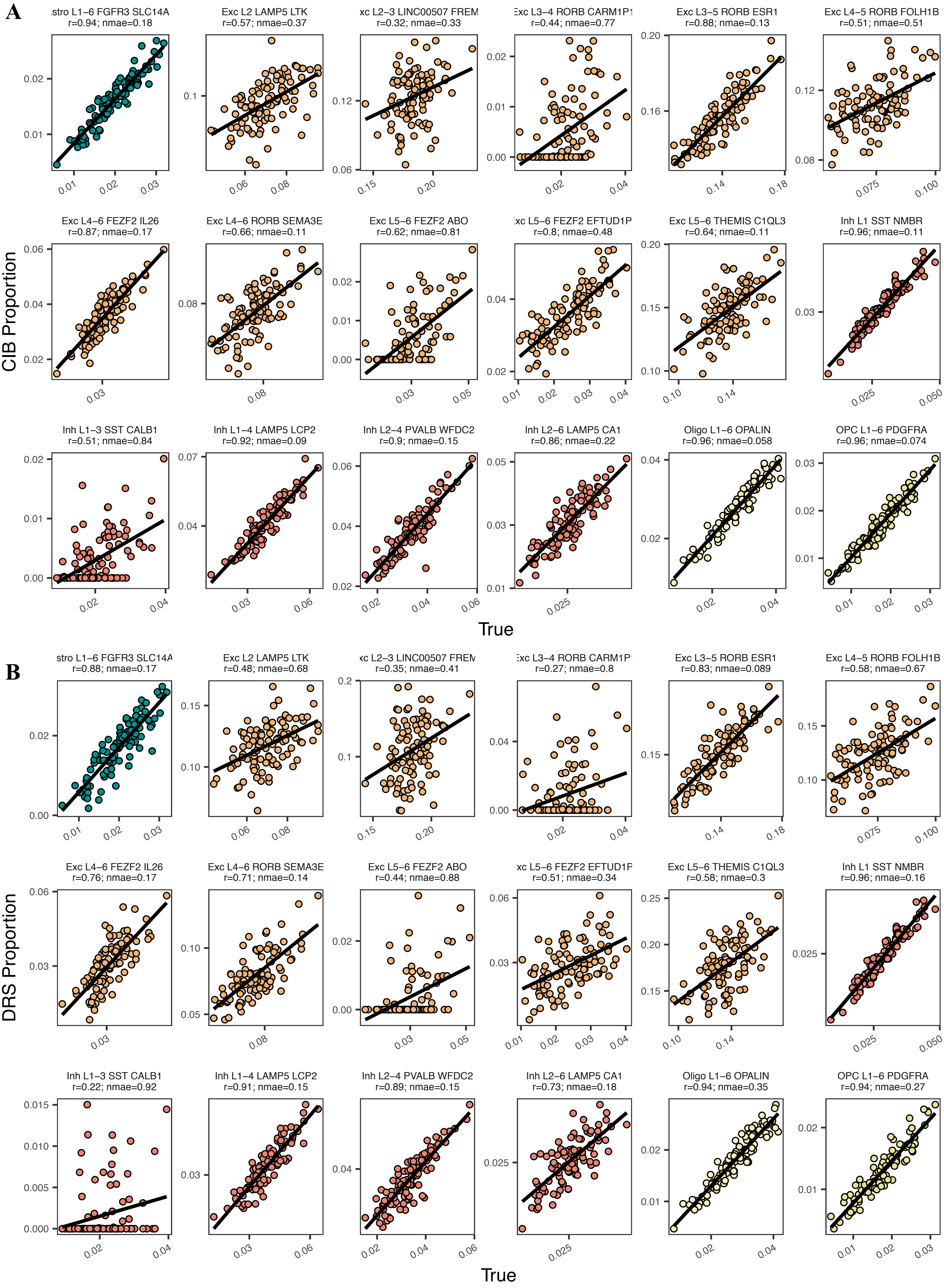


**Supplementary Figure 15. Scatterplots of true and deconvolution-estimated proportions in 100 CA-based *in silico* mixtures.** The signature used all cell-subtypes from the original publication by Hodge *et al.* (2019). **A.** CIBERSORT deconvolution. **B.** DeconRNASeq. *Solid black line*: regression line. *Nmae*: normalised mean absolute error. *r*: Pearson correlation coefficient.

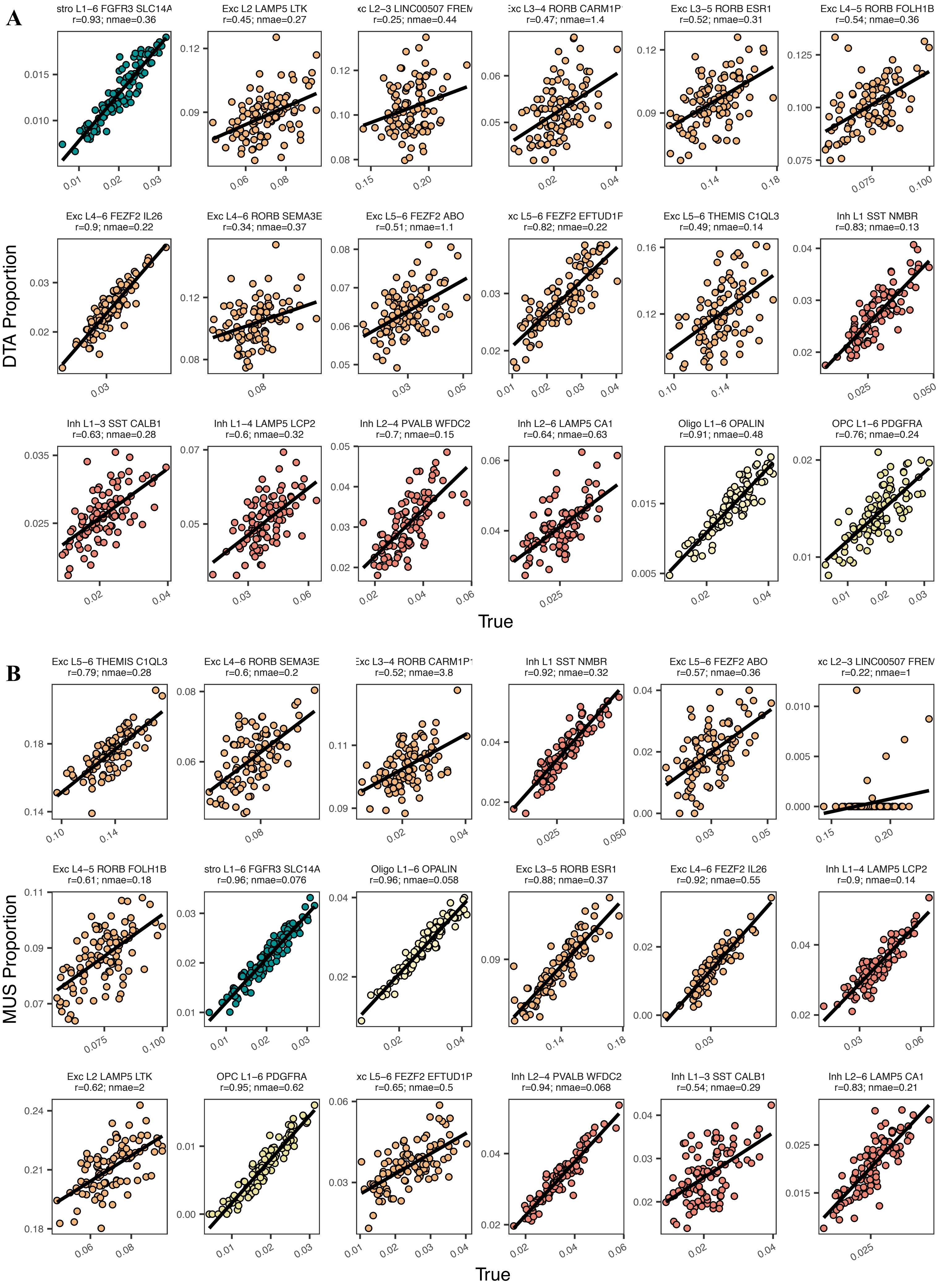


**Supplementary Figure 16. Scatterplots of true and deconvolution-estimated proportions in 100 CA *in silico* mixtures.** The signature used all cell-subtypes from the original publication by Hodge *et al.* (2019). **A.** dtangle deconvolution. **B.** MuSiC deconvolution. *Solid black line*: regression line. *Nmae*: normalised mean absolute error. *r*: Pearson correlation coefficient.

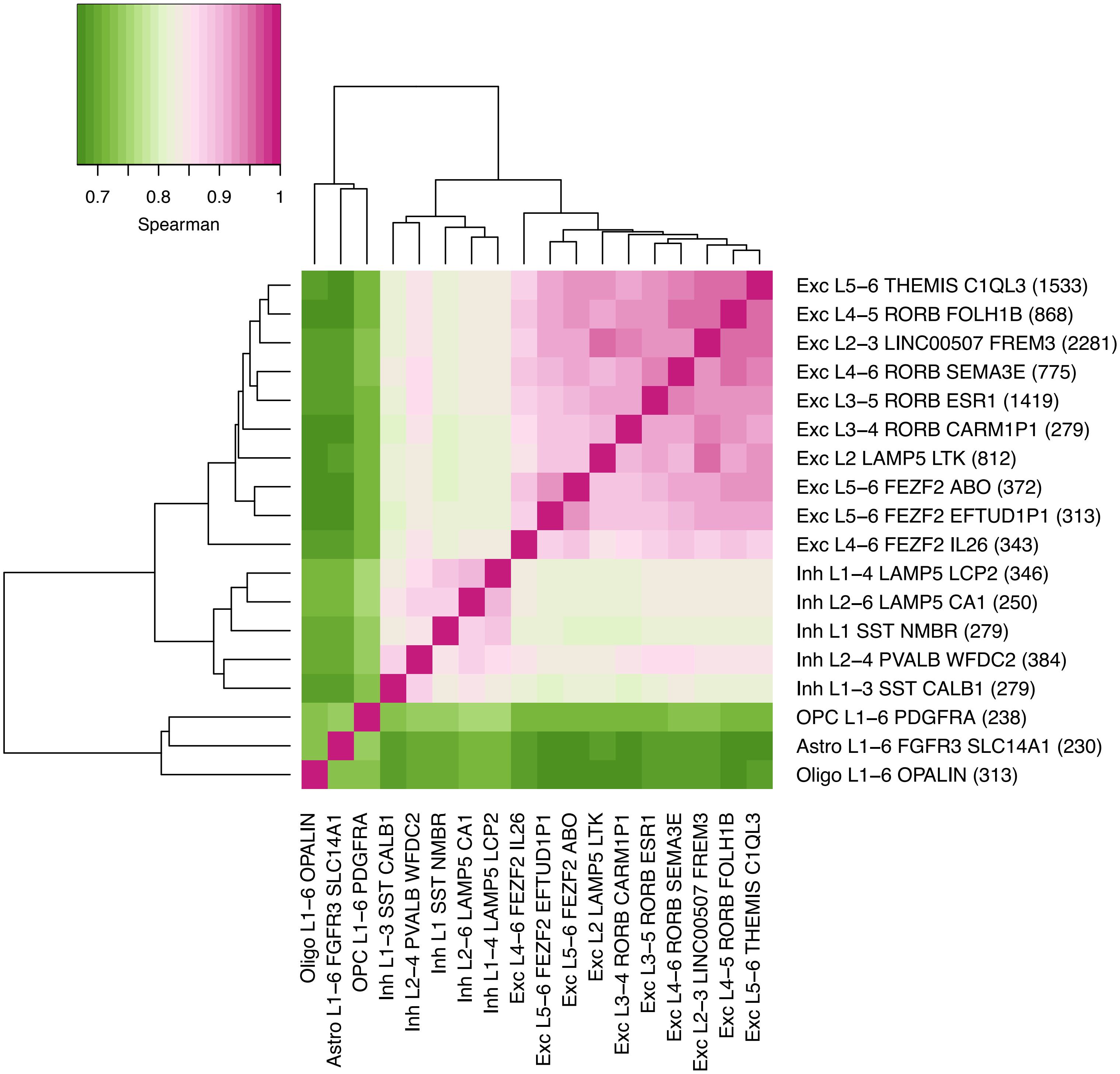


**Supplementary Figure 17. Heatmap of Spearman correlations between cell-subtypes in the CA dataset**. Labels are taken from the original publication. Numbers in brackets on the right axis labels indicate the number of nuclei in that class.

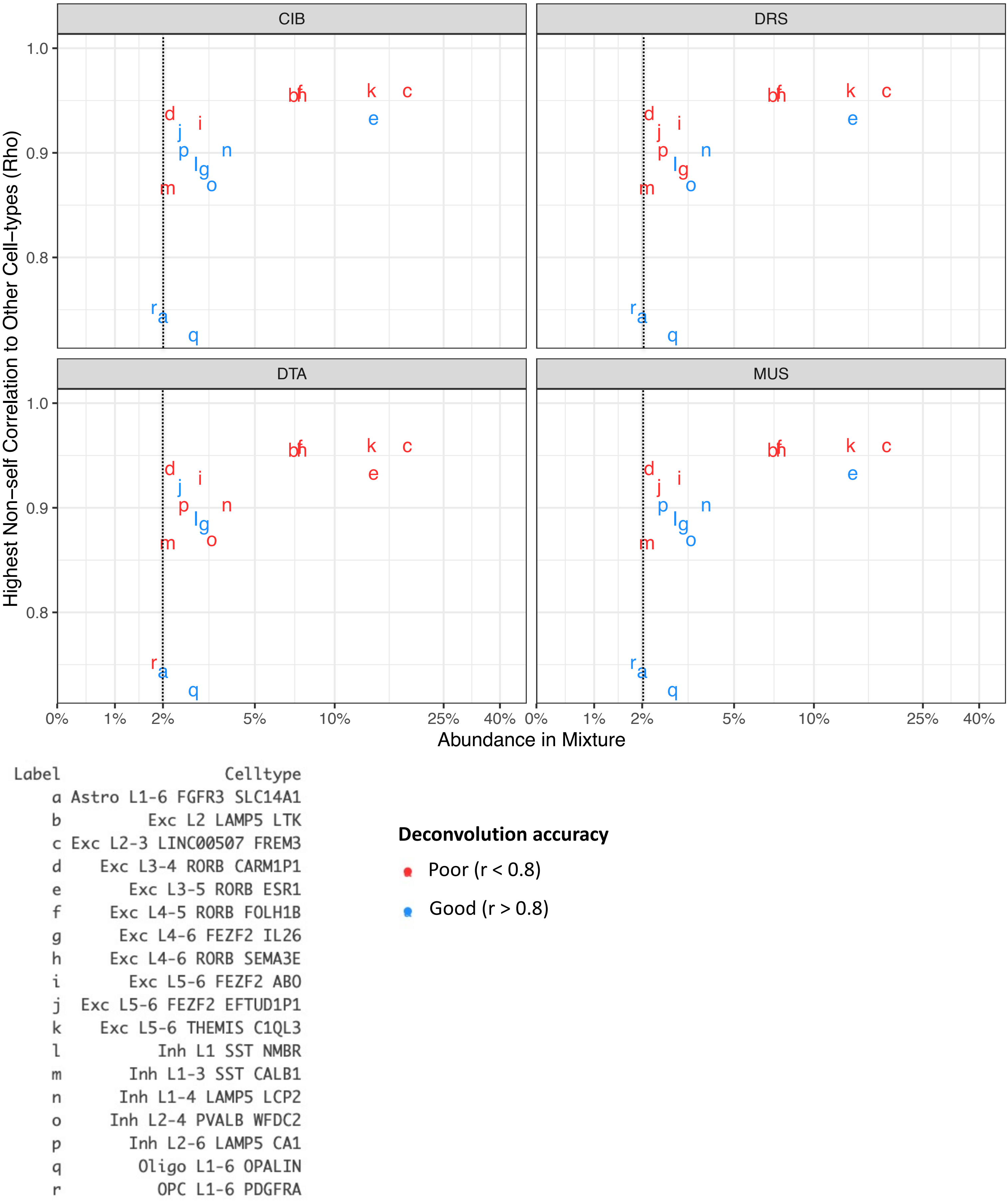


**Supplementary Figure 18. Effect of cell-type abundance and collinearity on deconvolution accuracy in CA-based simulations.** Each point represents a cell-subtype in the CA dataset. Points are labelled by text indicating the cell-subtype classification. Colours represent a binary code for good and poor deconvolution performance. *Rho*: Spearman correlation coefficient. *X axis*: mean abundance across the 100 simulated mixtures. *Y axis*: the highest correlation a cell-subtype has to any of the other cell-subtypes in the dataset, indicating collinearity. Note that the following labels are partially overlapping: "l", "g", and "o" at x=3, y=0.87; and "b", "f", and "h" at x=7, y=0.95

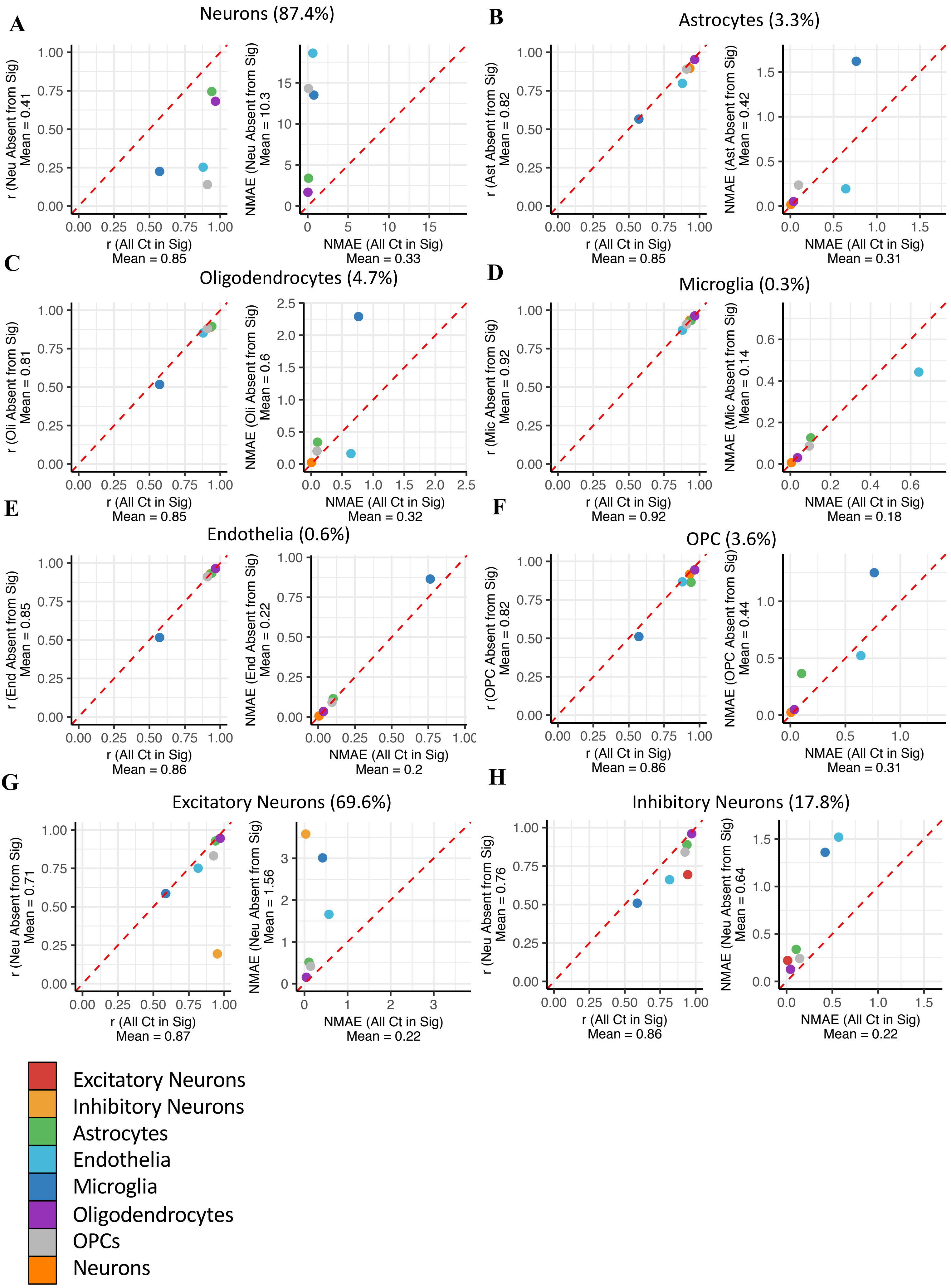


**Supplementary Figure 19. Effect of removing cell-types or cell-subtypes from the signature matrix.** For each cell-type, its mean abundance in the mixtures is shown in backets, and scatterplots display the deconvolution accuracy when all cell types are present in the signature (x-axis) vs. when the cell type is absent from signature (y-axis). Accuracy is measured as either *r* correlation coefficient (left panel) or normalised mean absolute error (right panel). Calculations of mean *r* and mean NMAE for the x-axis label do not include the absent cell-type, and thus differ across plots. *Dotted red line:* y = x.

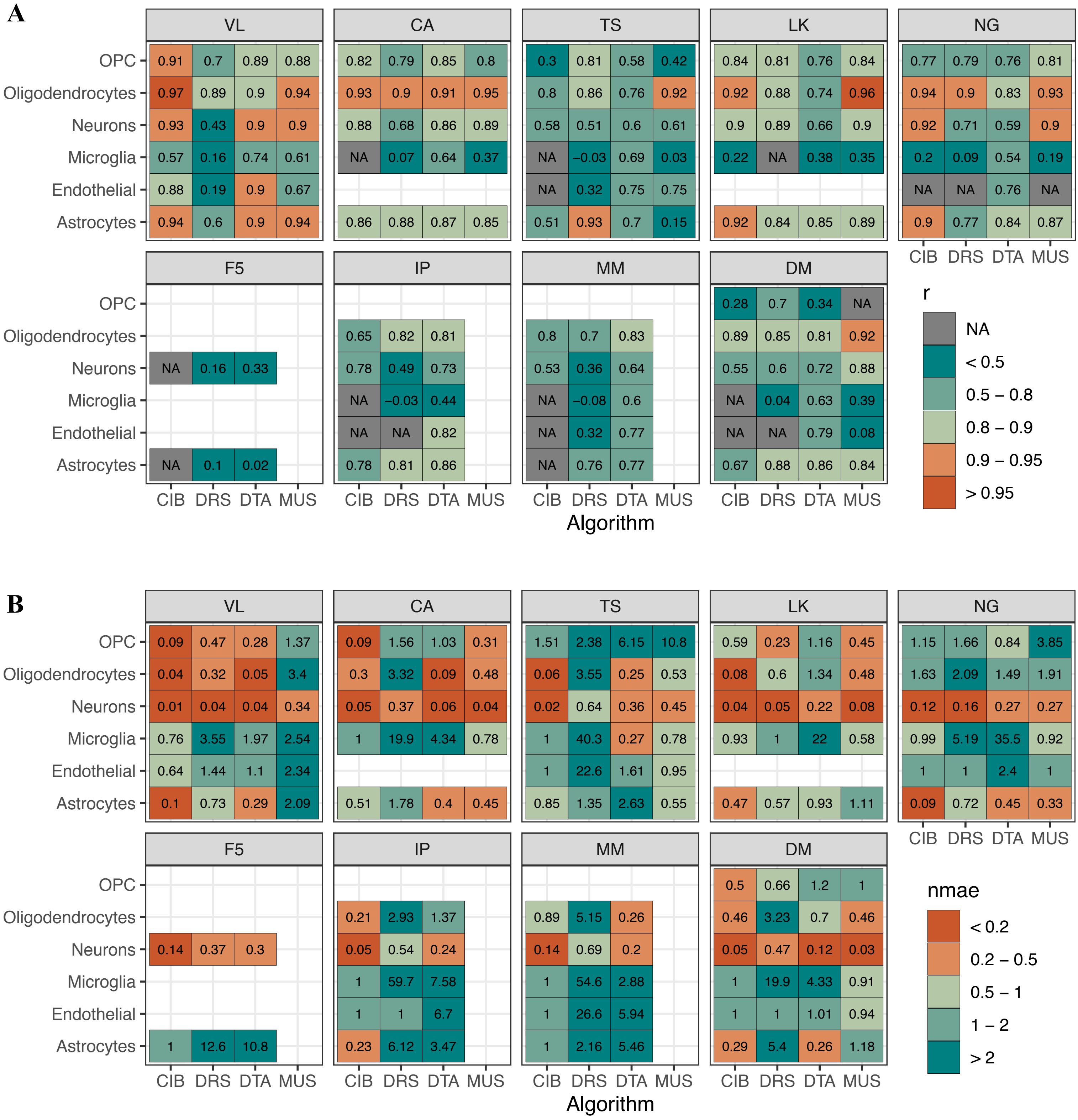


**Supplementary Figure 20. Effect of varying the signature in VL-based simulated mixtures**. Heatmaps of Pearson correlation (*r*; **A.**) and normalised mean absolute error (nmae; **B.**) for estimated versus true proportion when varying the reference signature. The mixtures are 100 *in silico* VL simulations. Signatures only included a pan-neuronal expression profile, rather than excitatory or inhibitory sub-types. Blank squares indicate that the cell-type was not present in the signature, and thus no statistic was calculated. Grey squares indicate NA, indicating that the cell-type was present in the signature but the statistic could not be calculated; for *r*, this means there was no variance in the composition estimates, typically meaning all 100 samples' estimates were 0 or 1. For more details about signature characteristics, see methods.

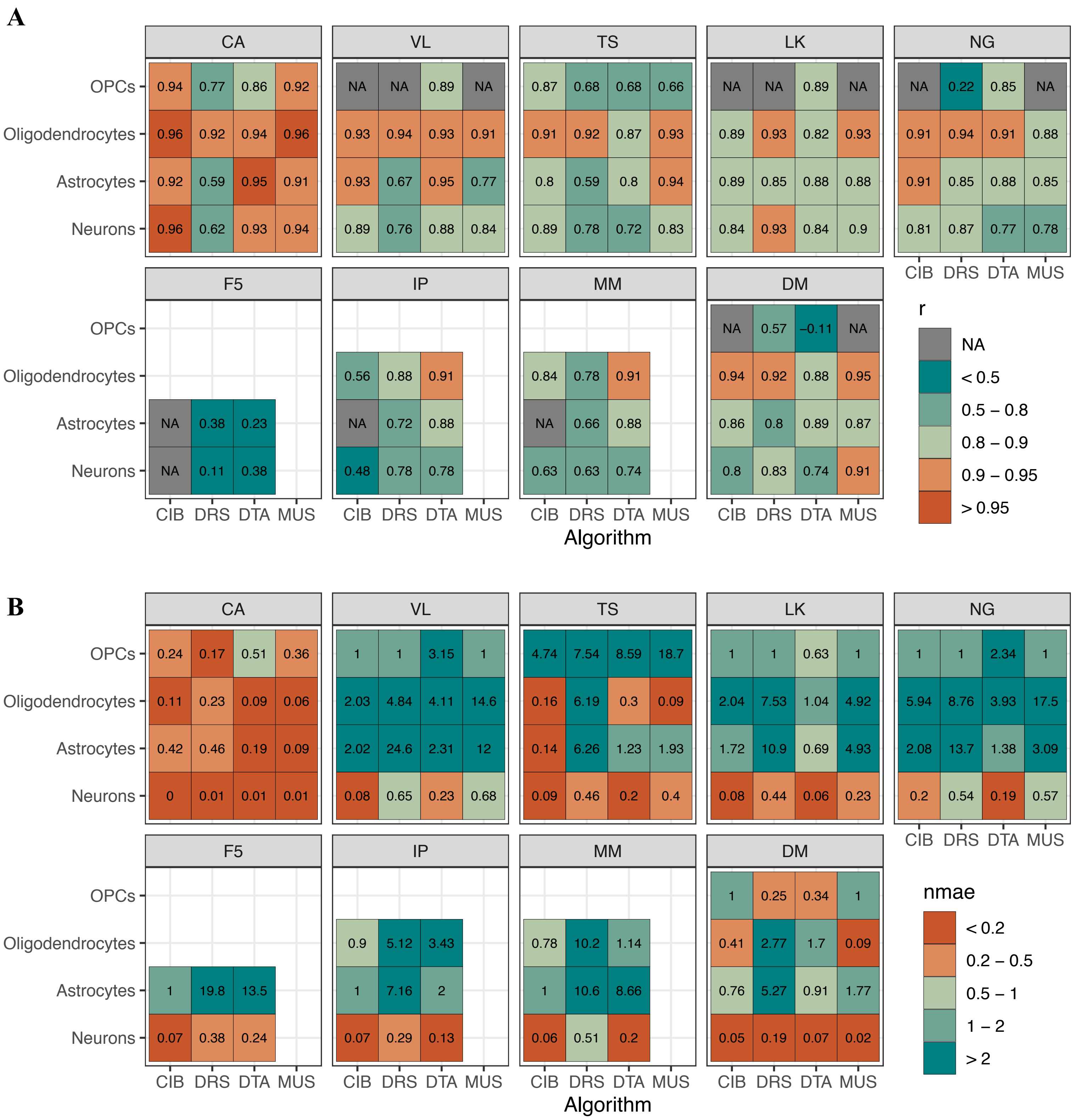


**Supplementary Figure 21. Effect of varying the signature in CA-based simulated mixtures**. Heatmaps of Pearson correlation (*r*; **A.**) and normalised mean absolute error (nmae; **B.**) for estimated versus true proportion when varying the reference signature. The mixtures are 100 *in silico* CA simulations. Signatures only included a pan-neuronal expression profile, rather than excitatory or inhibitory sub-types. Blank squares indicate that the cell-type was not present in the signature, and thus no statistic was calculated. Grey squares indicate NA, indicating that the cell-type was present in the signature but the statistic could not be calculated; for *r*, this means there was no variance in the composition estimates, typically meaning all 100 samples' estimates were 0 or 1. For more details about signature characteristics, see methods.

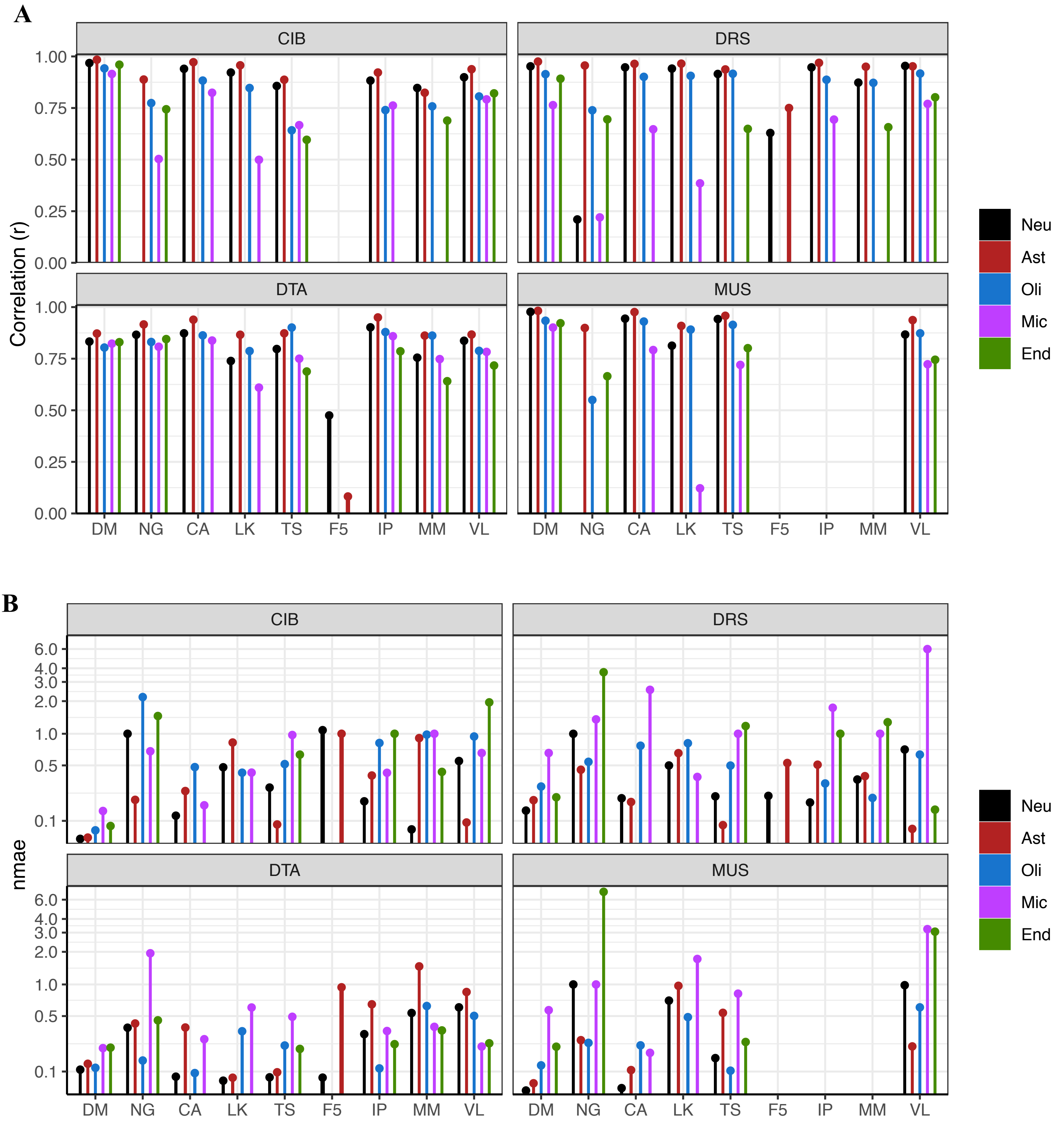


**Supplementary Figure 22. Effect of varying the signature in DM-based simulated mixtures.** Heatmaps of Pearson correlation ($r$; top panel) and normalised mean absolute error (nmae; bottom panel) for estimated versus true proportion when varying the reference signature. Signatures only included a pan-neuronal expression profile, rather than excitatory or inhibitory sub-types. *Neu*: Neurons. *Ast*: Astrocytes. *Oli*: Oligodendrocytes. *Mic*: Microglia. *End*: Endothelia. For more details about signature characteristics, see methods.

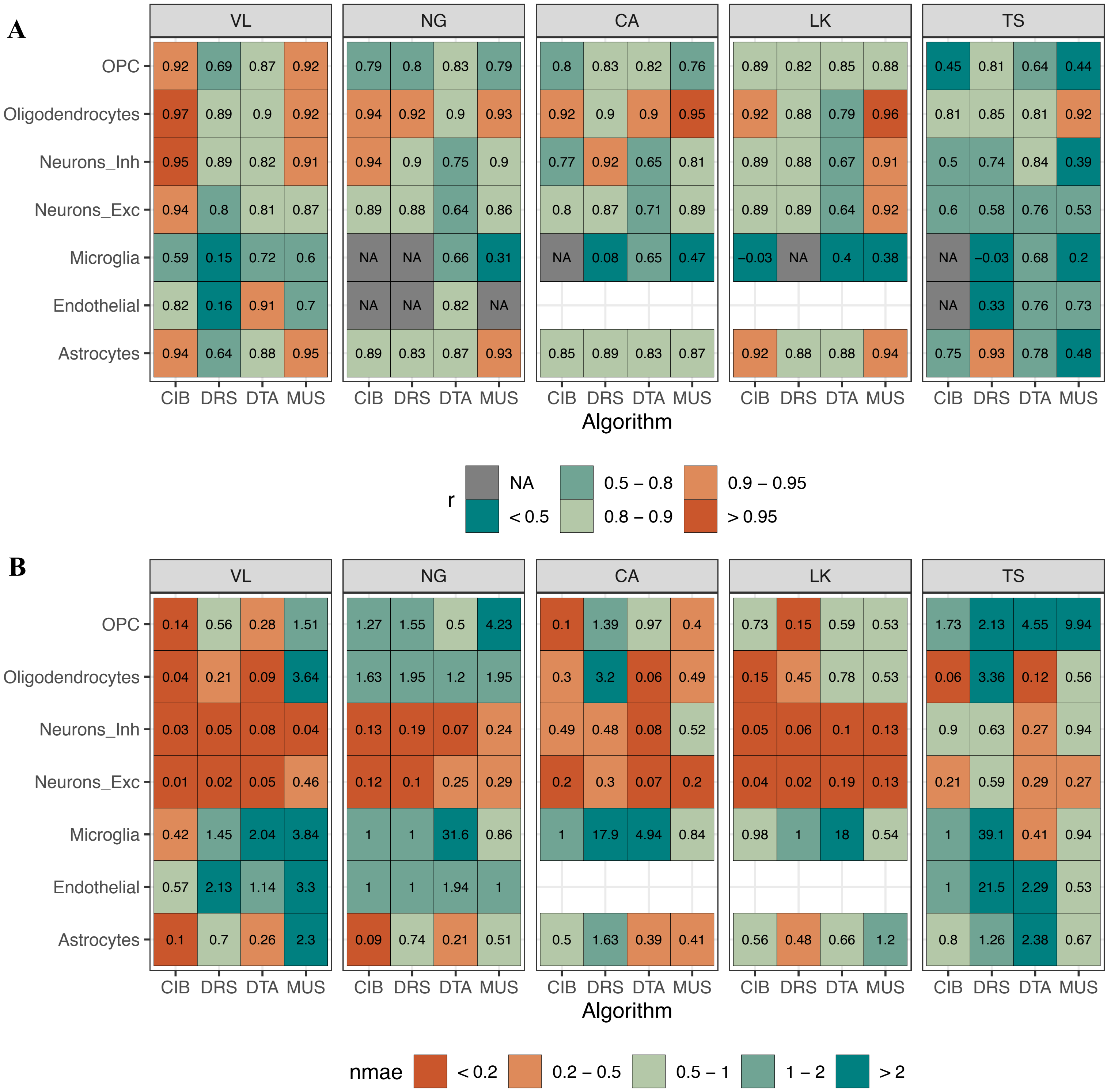


**Supplementary Figure 23. Effect of varying the signature while including neuronal subtypes in VL-based simulated mixtures.** Heatmaps of Pearson correlation (*r*; **A.**) and normalised mean absolute error (nmae; **B.**) for estimated versus true proportion when varying the reference signature. The mixtures are 100 *in silico* VL simulations. All signatures here include information about the broad neuronal subtype (excitatory or inhibitory). Blank squares indicate that the cell-type was not present in the signature, and thus no statistic was calculated. Grey squares indicate NA, indicating that the cell-type was present in the signature but the statistic could not be calculated; for *r*, this means there was no variance in the composition estimates, typically meaning all 100 samples' estimates were 0 or 1. For more details about signature characteristics, see methods.

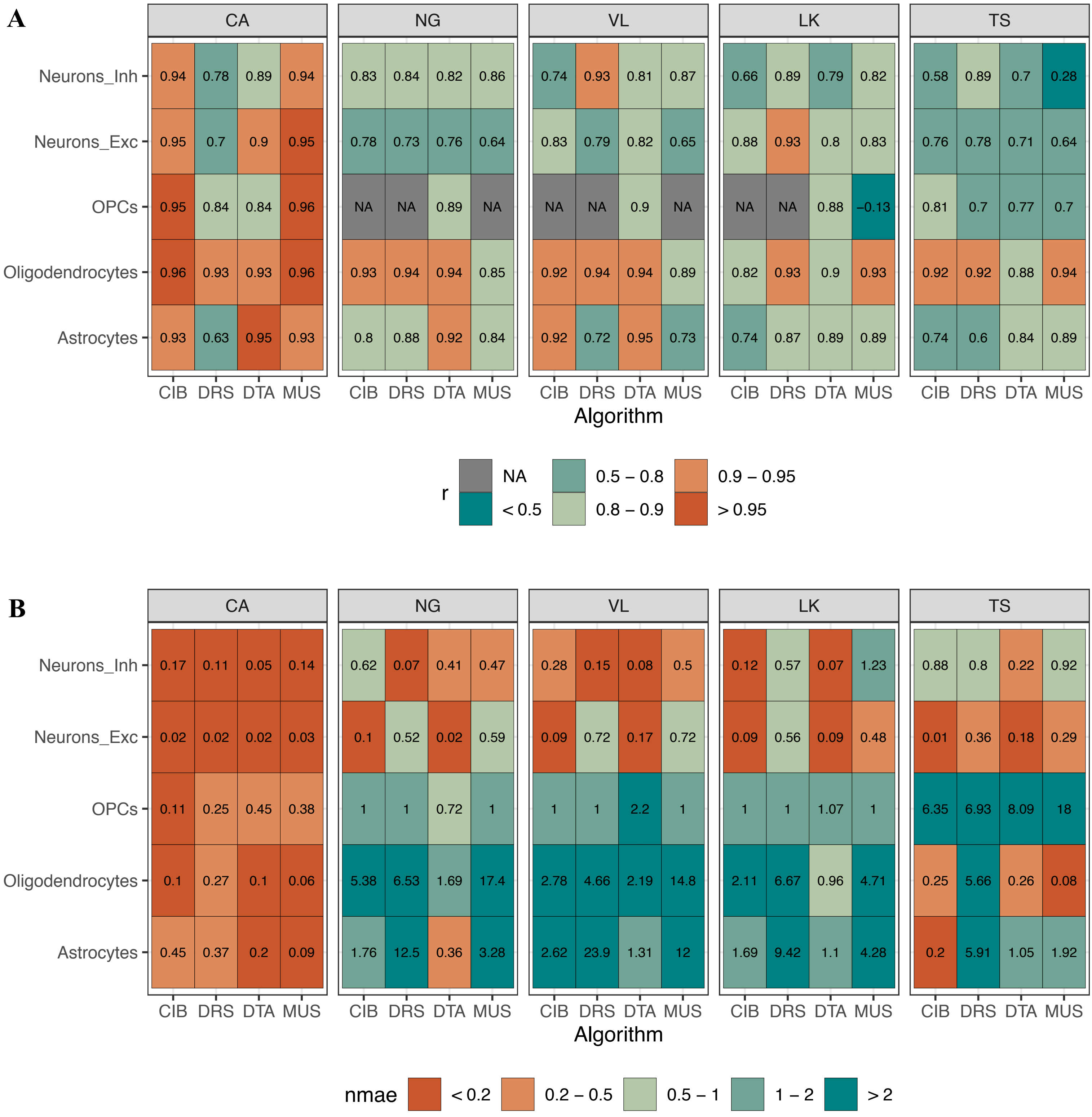


**Supplementary Figure 24. Effect of varying the signature while including neuronal subtypes in CA-based simulated mixtures.** Heatmaps of Pearson correlation (*r*; **A.**) and normalised mean absolute error (nmae; **B.**) for estimated versus true proportion when varying the reference signature. The mixtures are 100 *in silico* CA simulations. All signatures here include information about the broad neuronal subtype (excitatory or inhibitory). Blank squares indicate that the cell-type was not present in the signature, and thus no statistic was calculated. Grey squares indicate NA, indicating that the cell-type was present in the signature but the statistic could not be calculated; for *r*, this means there was no variance in the composition estimates, typically meaning all 100 samples' estimates were 0 or 1. For more details about signature characteristics, see methods.

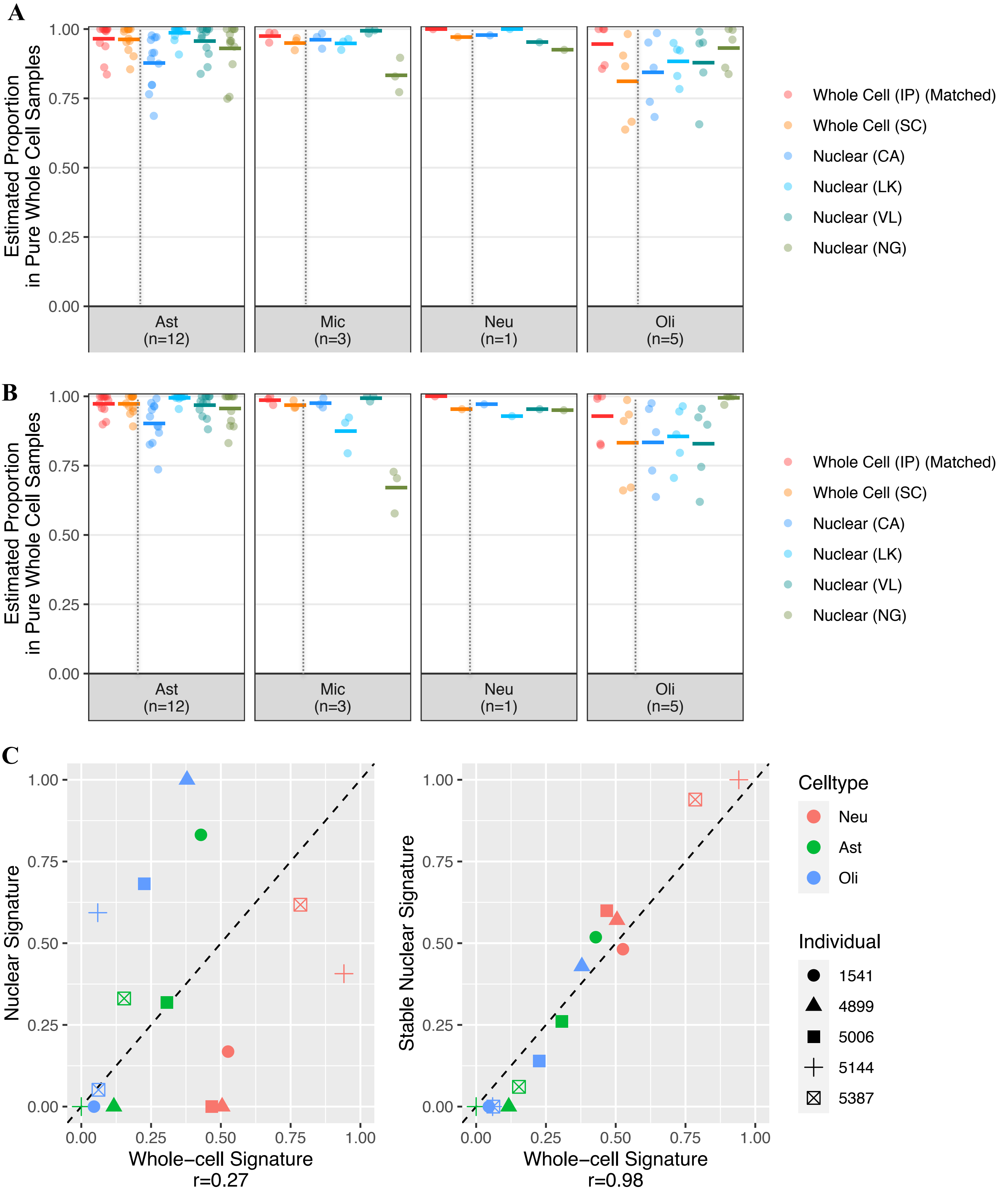


**Supplementary Figure 25. The role of compartment-specific genes when using snRNA-seq signatures. A and B:** Estimated proportions for pure samples of immuno-panned cell-types, using whole-cell and snRNA-seq signatures. (A) all genes were included in the signature. (B) compartment-specific genes were filtered-out. *Thick horizontal line*: mean. *Dotted vertical line*: separates whole-cell from snRNA-seq signatures. *Neu*: neurons. *Ast*: astrocytes. *Oli*: oligodendrocytes. *Mic*: microglia. **C.** Scatterplot of estimated proportions for bulk brain samples using the RNA-seq signature, IP (x-axis) or the snRNA-seq signature derived from the same individual (y-axis). *Left*: all genes were included in the signature; *right*: compartment-specific genes were filtered-out. *Individual*: NICHD brain bank id number. Cell-type proportions were estimated using CIBERSORT.

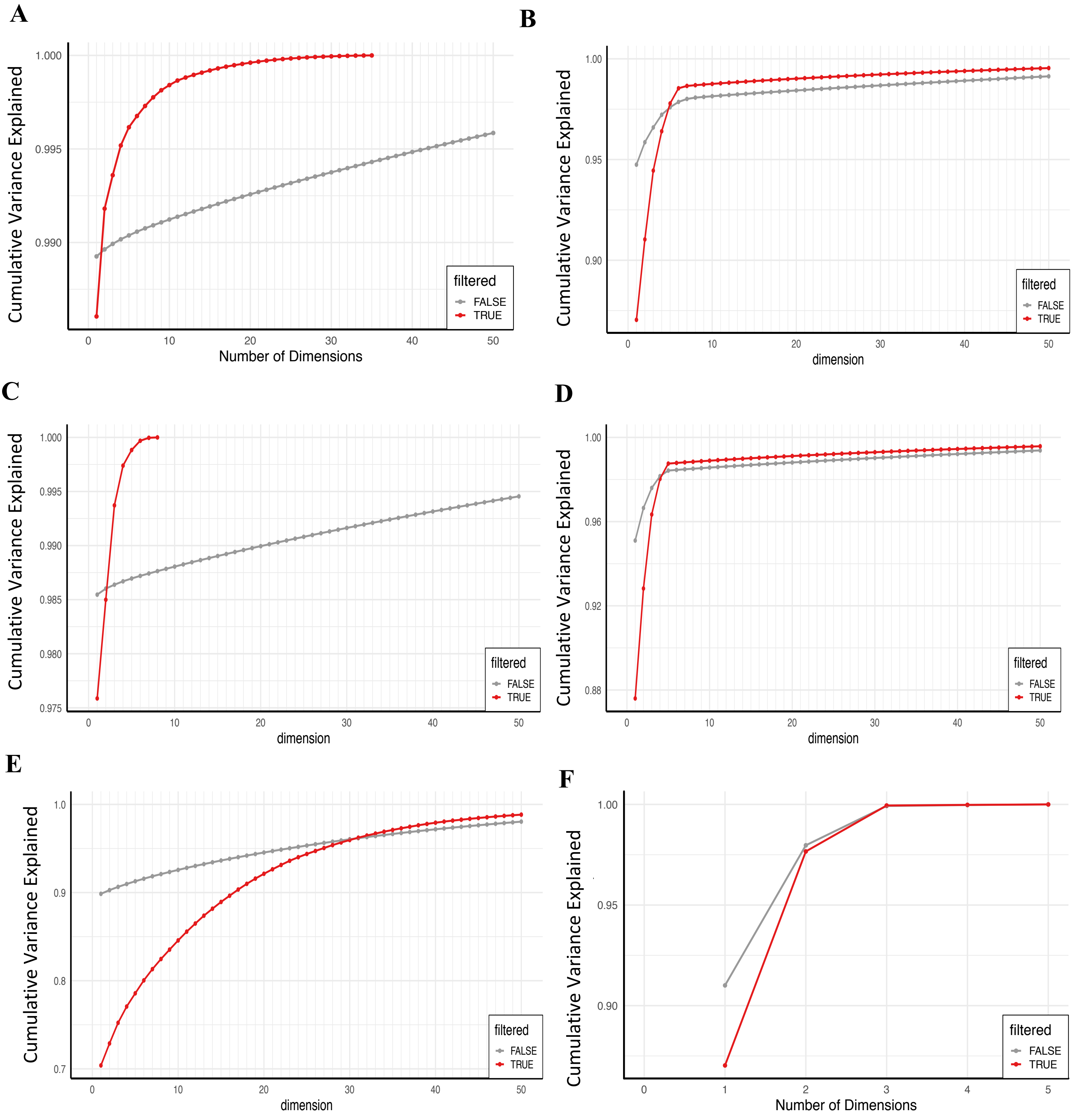


**Supplementary Figure 26. Proportion of variance in gene expression explained by singular-value decomposition.** Linseed proposes that the saturation point of the curve (*i.e.* the number of linearly-independent components that contribute to the mixture) is its number of constituent cell-types. **A-B.** VL-derived mixtures based on random sampling (A) and those simulated with a wide range and variance in cell-type composition (B). **C-D.** CA mixtures based on random sampling (C) and and those simulated with a wide range and variance in cell-type composition (D). **E.** DM mixtures based on random sampling. **F.** RNA mixtures.

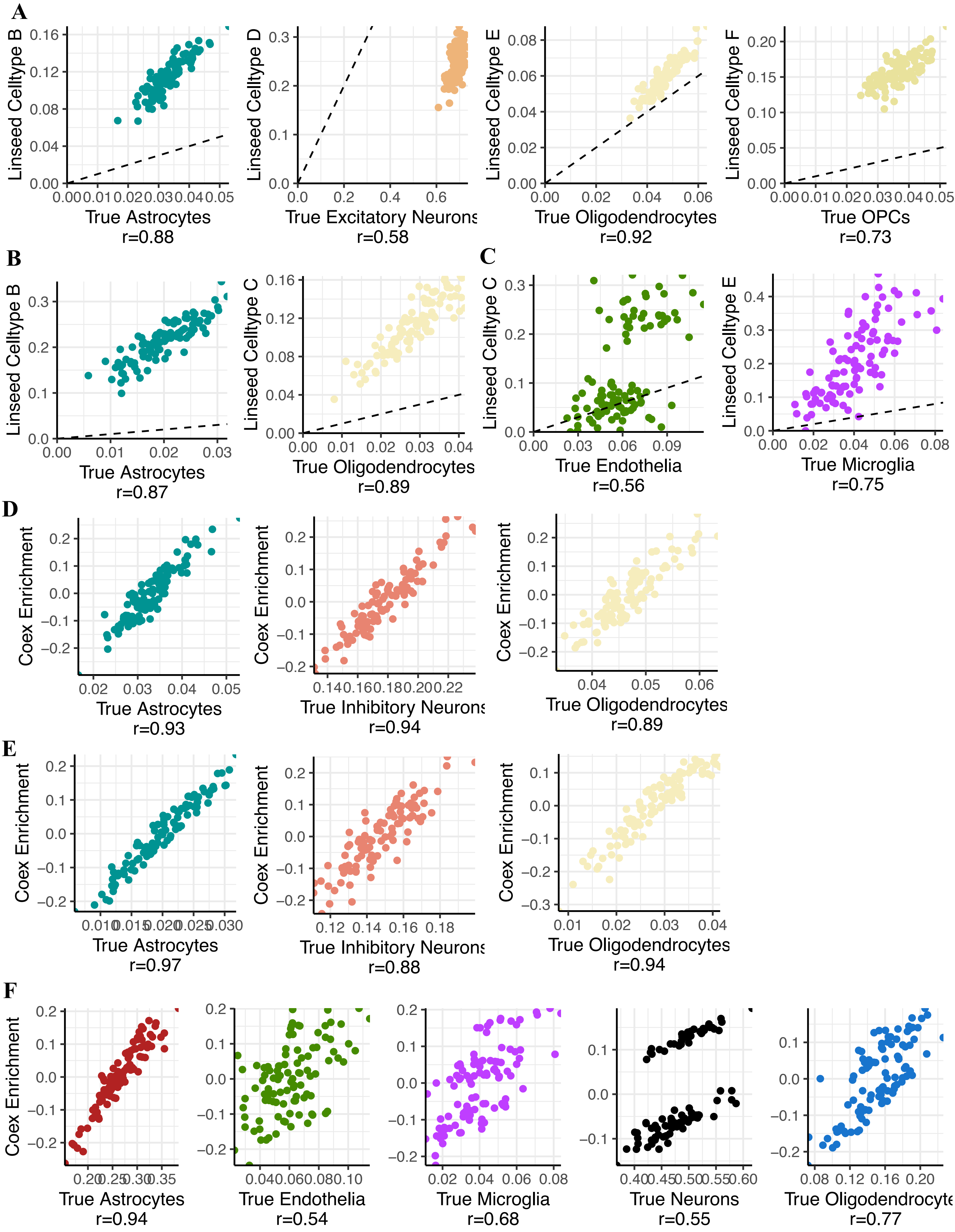


**Supplementary Figure 27. Scatterplots of reference-free deconvolution estimates versus true proportion**. **A-C.** Linseed. Plots are only shown for inferred cell-types correlated to a true cell-type at r > 0.5. A. VL-derived random simulations. B. CA-derived random simulations. C. DM-derived random simulations. *Dotted line*: y = x. **D-F.** Coex. Plots are only shown for inferred cell-types that were assigned to a true cell-type through marker enrichment analysis (Fisher Test, $p<1x10^{-5}$, odds ratio > 5; Methods) . D. VL-derived random simulations. E. CA-derived random simulations. F. DM-derived random simulations.

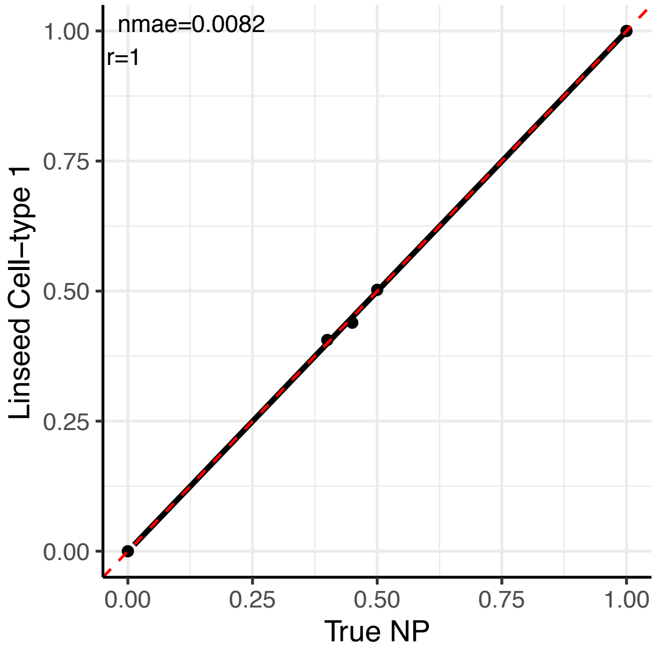


**Supplementary Figure 28.** Scatterplot of proportions estimated by Linseed in the RNA mixtures of neurons and astrocytes setting number of cell types *k*=2. *Black line*: regression line. *Red dotted line*: y=x.

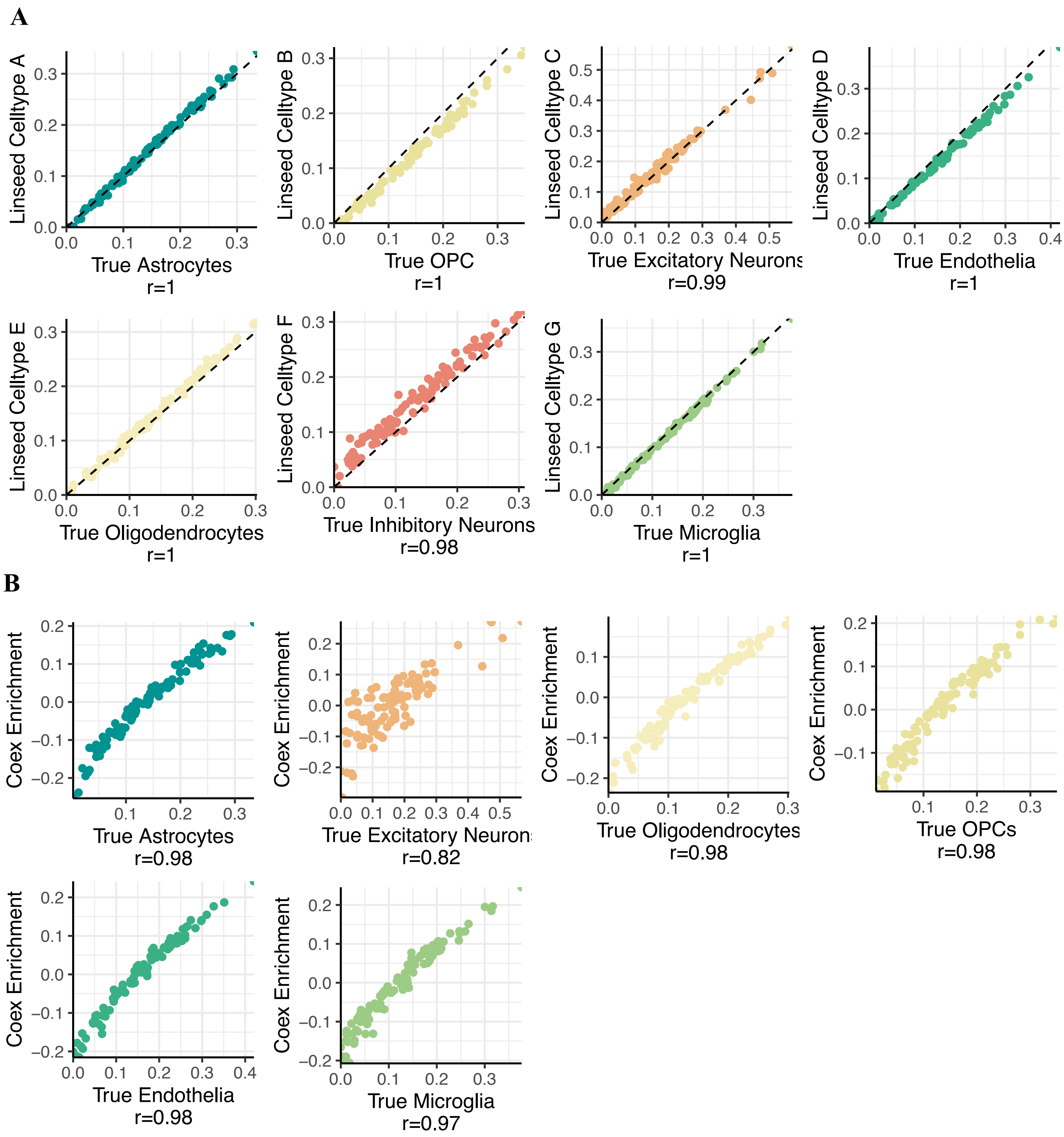


**Supplementary Figure 29.** Scatterplots of reference-free deconvolution estimates versus true proportion in simulations with increased cell-type variance. Simulations were based on VL single-nuclei. **A.** Linseed. **B.** Coex.

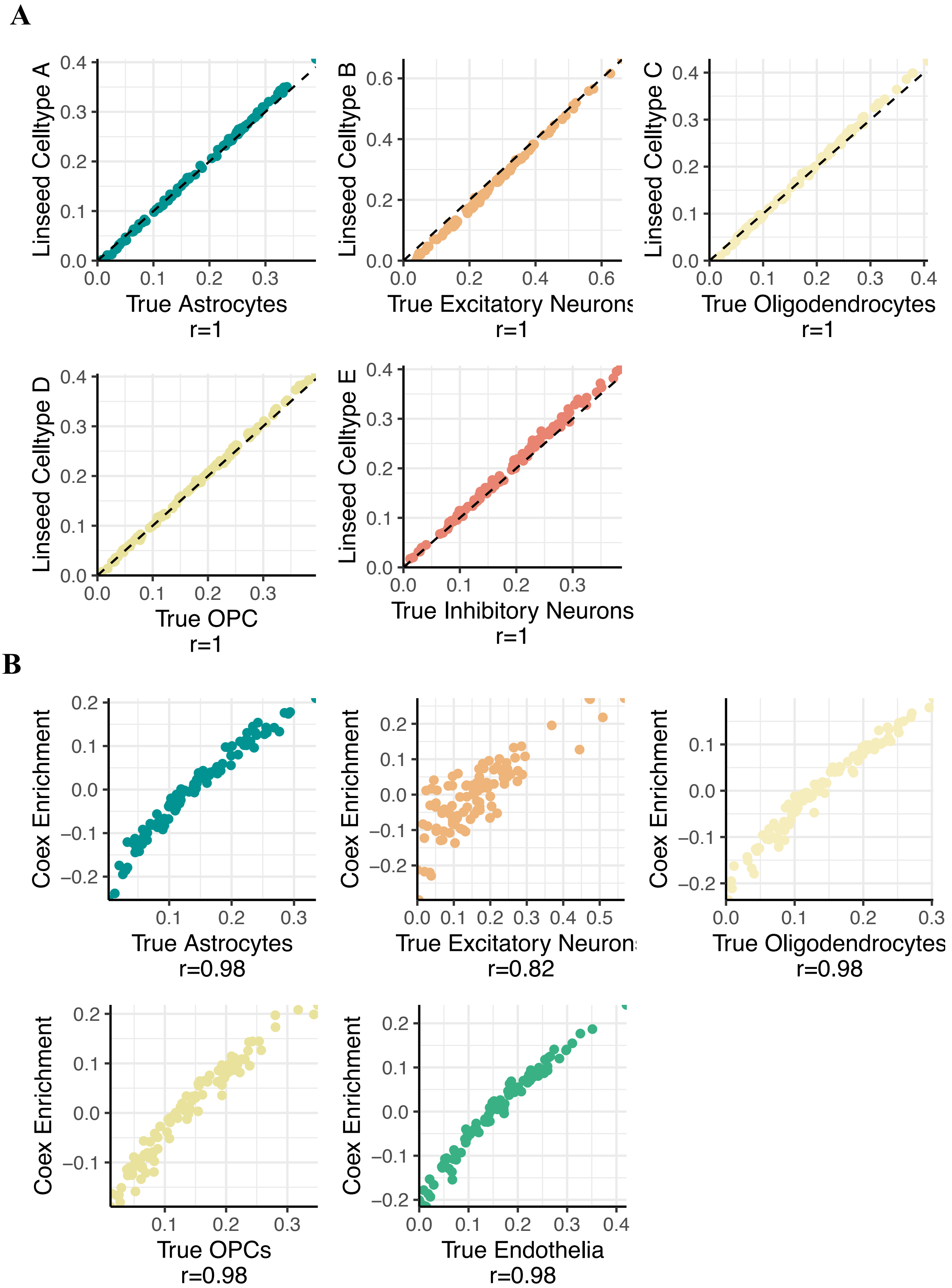


**Supplementary Figure 30.** Scatterplots of reference-free deconvolution estimates versus true proportion in simulations with increased cell-type variance. Simulations were based on CA single-nuclei. **A.** Linseed. **B.** Coex.

A

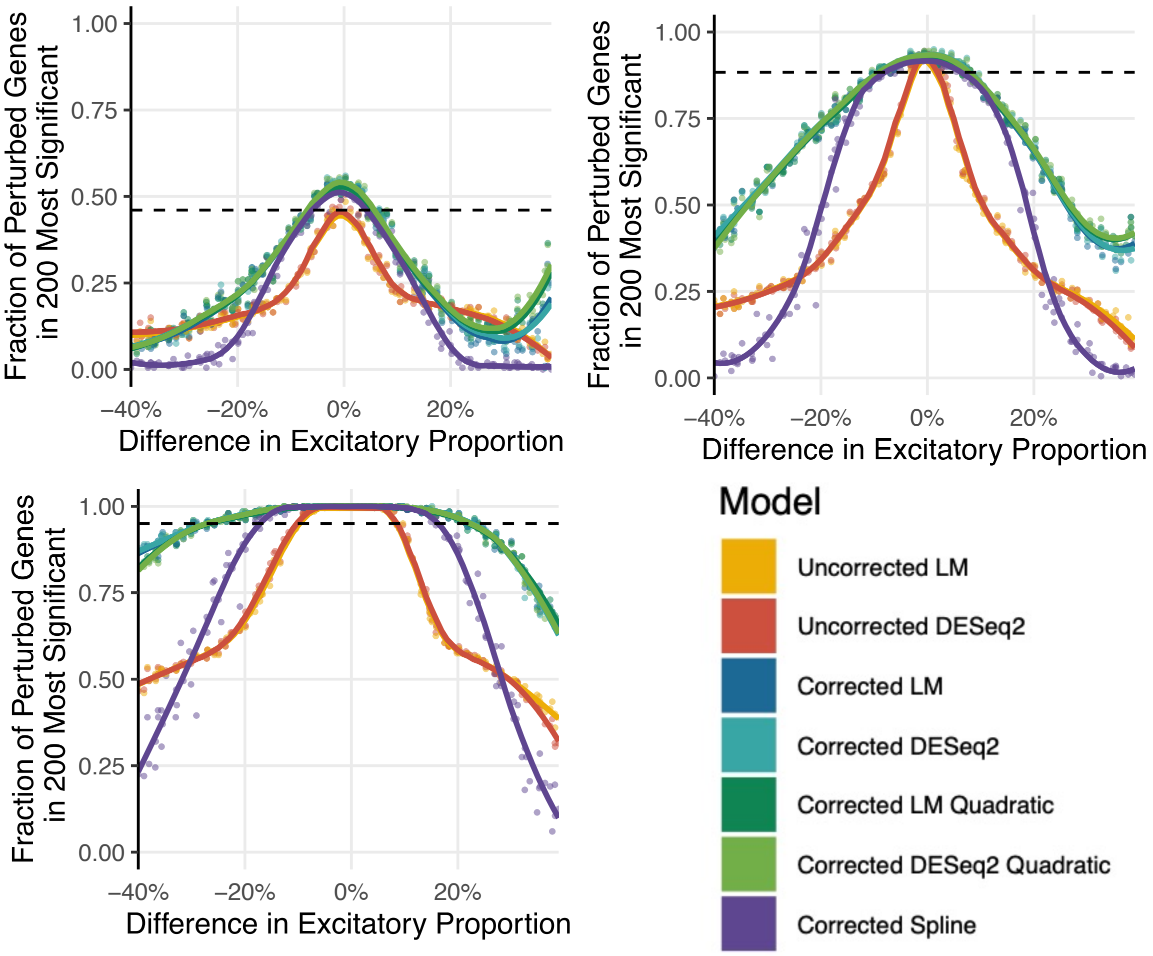


B

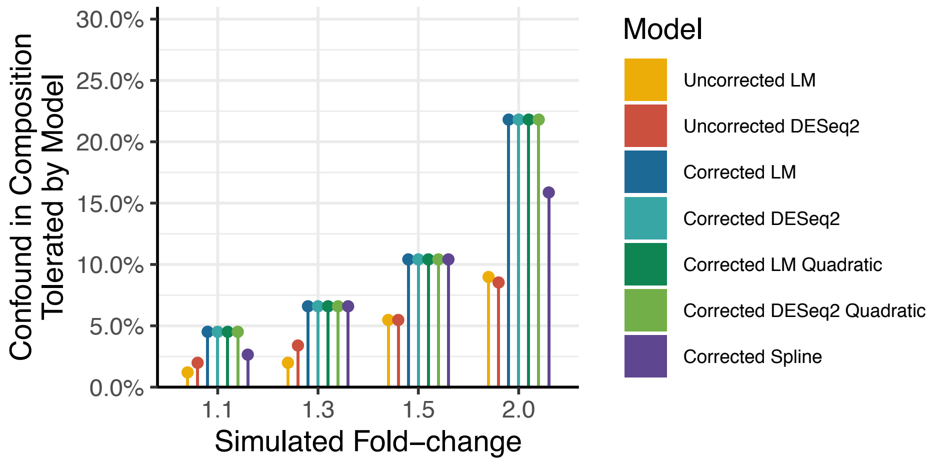


**Supplementary Figure 31. Interplay between confounds in excitatory cell-type proportion and differential expression for true up- or down-regulated genes**. **A.** Scatterplots show the relationship between the confound in excitatory proportion within a simulation, and the discriminatory ability (fraction of known perturbed genes in the 200 genes with the smallest p-value) (left), the true positive rate for marker genes of excitatory neurons (middle), and the true positive rate for non-marker genes (right). *Top-left*: true fold-change of 1.1. *Top-right*: true fold-change of 1.3. *Bottom-left*: true fold-change of 2. *Coloured lines*: local regression line. *Dotted line in the left* panel: 0.95 times the discriminatory ability for LM when x = 0. **B.** Model robustness to composition confounds. Barplots show the smallest negative composition change where discriminatory ability fell below 0.95 of the baseline (*i.e.* that from an uncorrected linear model on a simulation with no composition confound). Labels are per A.

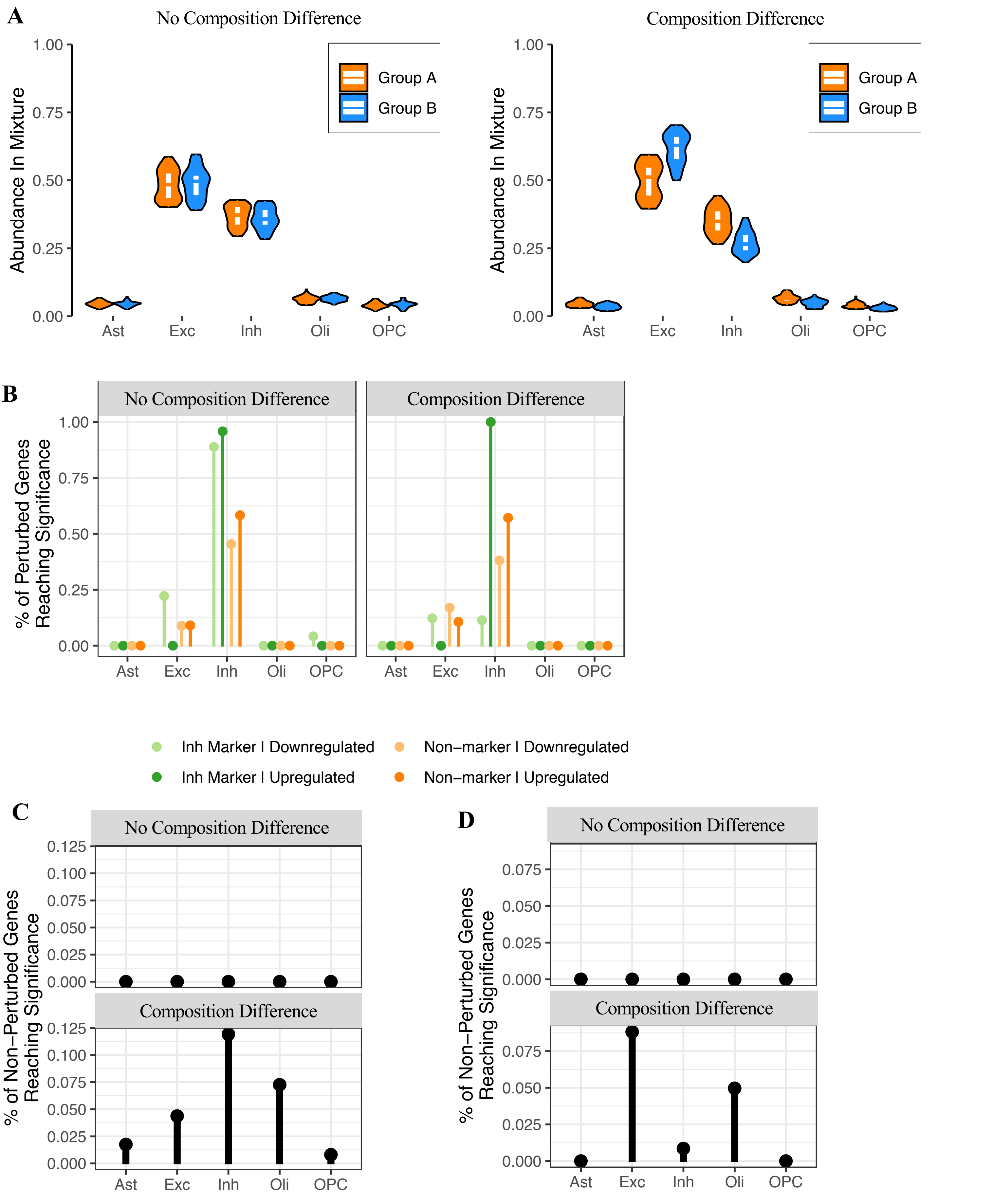


**Supplementary Figure 32. Cell-type-specific differential expression analysis using CIBERSORTx. A.** Composition distribution of simulated datasets. *Left*: simulated data without a composition difference between the two groups. *Right*: simulated data with a composition difference between the two groups. Each group contained 50 samples. **B.** Gene expression was perturbed 1.5-fold in Group B in inhibitory neurons for 100 non-marker genes plus 100 inhibitory neuronal marker genes. CIBERSORTx was used to extract cell-type-specific expression (Methods), with a linear model then run to assess differential expression in each cell-type. The plot displays the fraction of the true perturbed genes with an FDR < 0.05. Note that the fraction was calculated using only the subset of perturbed genes which were detected in the given cell-type. **C and D.** False positive rate across the different simulations when expression was perturbed in either excitatory neurons (C) or inhibitory neurons (D).

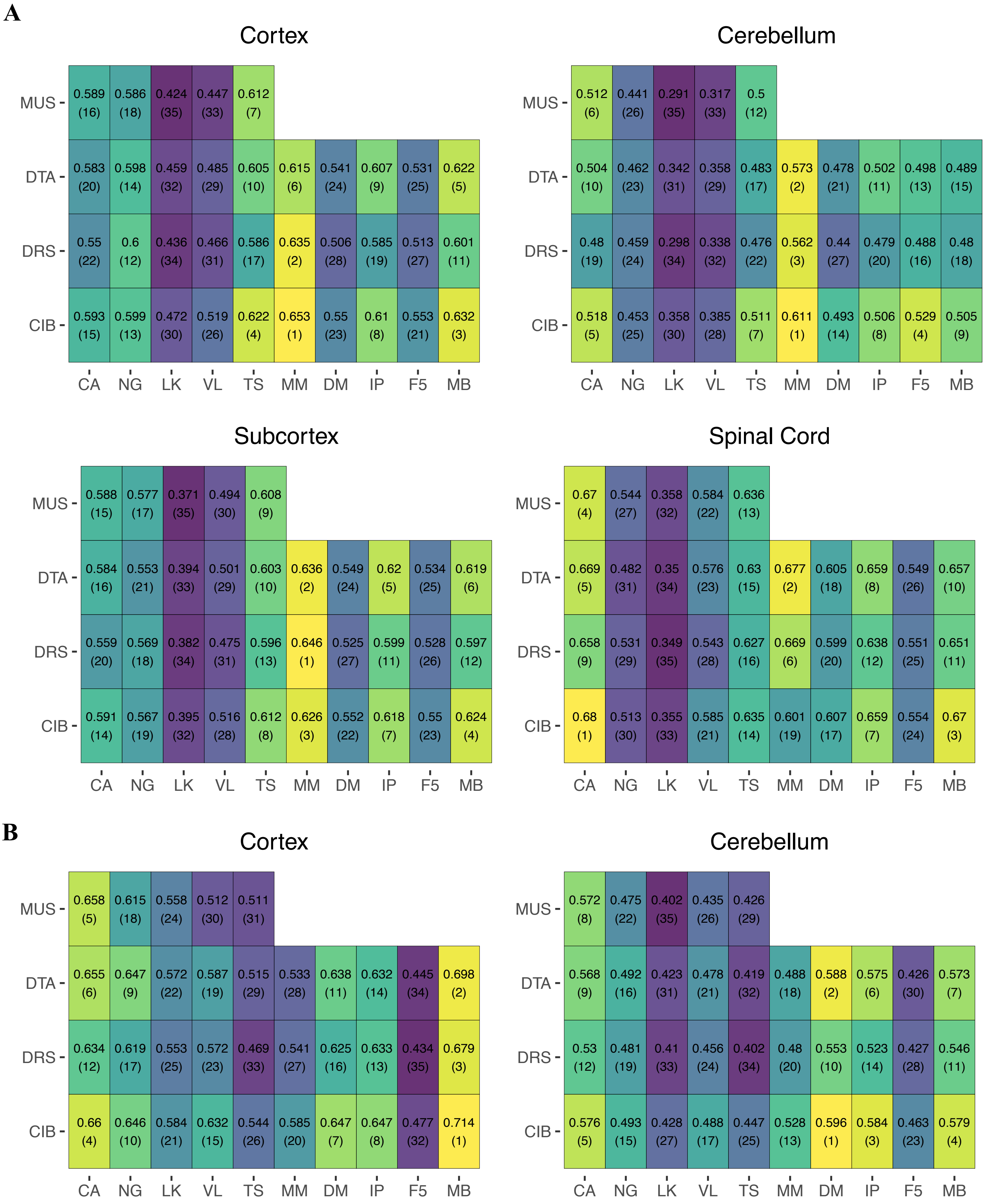


**Supplementary Figure 33. Median goodness-of-fit in large bulk brain RNA-seq datasets across signatures and algorithms. A.** GTEx. **B.** Parikshak *et al.*. Each panel aggregates results from samples from a given region. Rows represent signatures, and columns represent algorithms. Within each cell, the number of top is the median goodness-of-fit, while the number in parentheses below is its rank across all algorithm/signature combinations. Colours represent rank, ranging from purple (worst performance and high rank) to yellow (best performance and low rank)

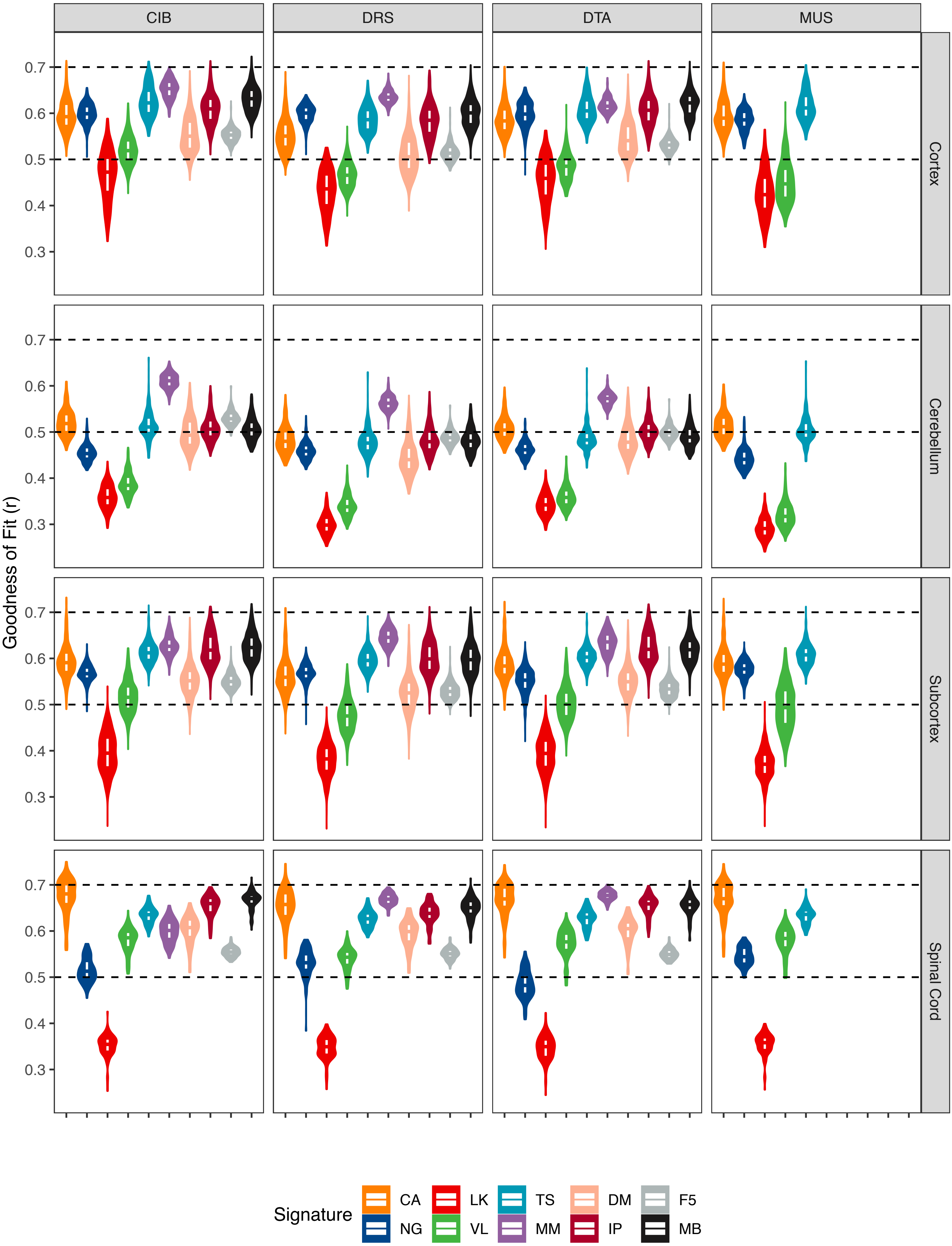


**Supplementary Figure 34. Violin plots of goodness of fit across signatures and regions in GTEx data.** The top, middle, and bottom of the white internal boxes mark the 75th, 50th, and 25th percentiles, respectively. Note that the data presented in the top-left panel (CIB/Cortex) was also shown in Figure 5A.

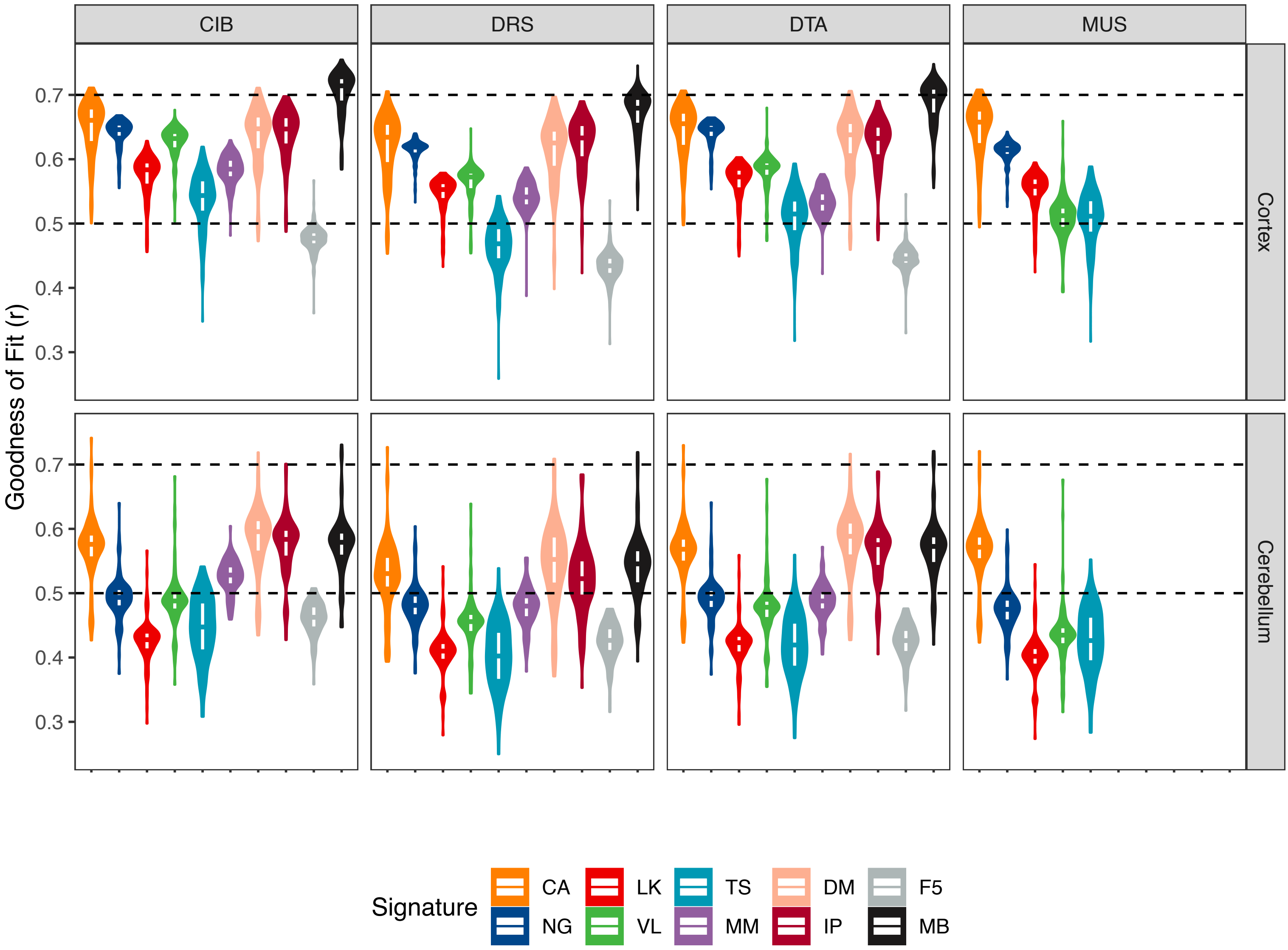


**Supplementary Figure 35. Violin plots of goodness of fit across signatures and regions in the Parikshak dataset.** The top, middle, and bottom of the white internal boxes mark the 75th, 50th, and 25th percentiles, respectively. Note that the data presented in the top-left panel (CIB/Cortex) was also shown in Figure 5B.

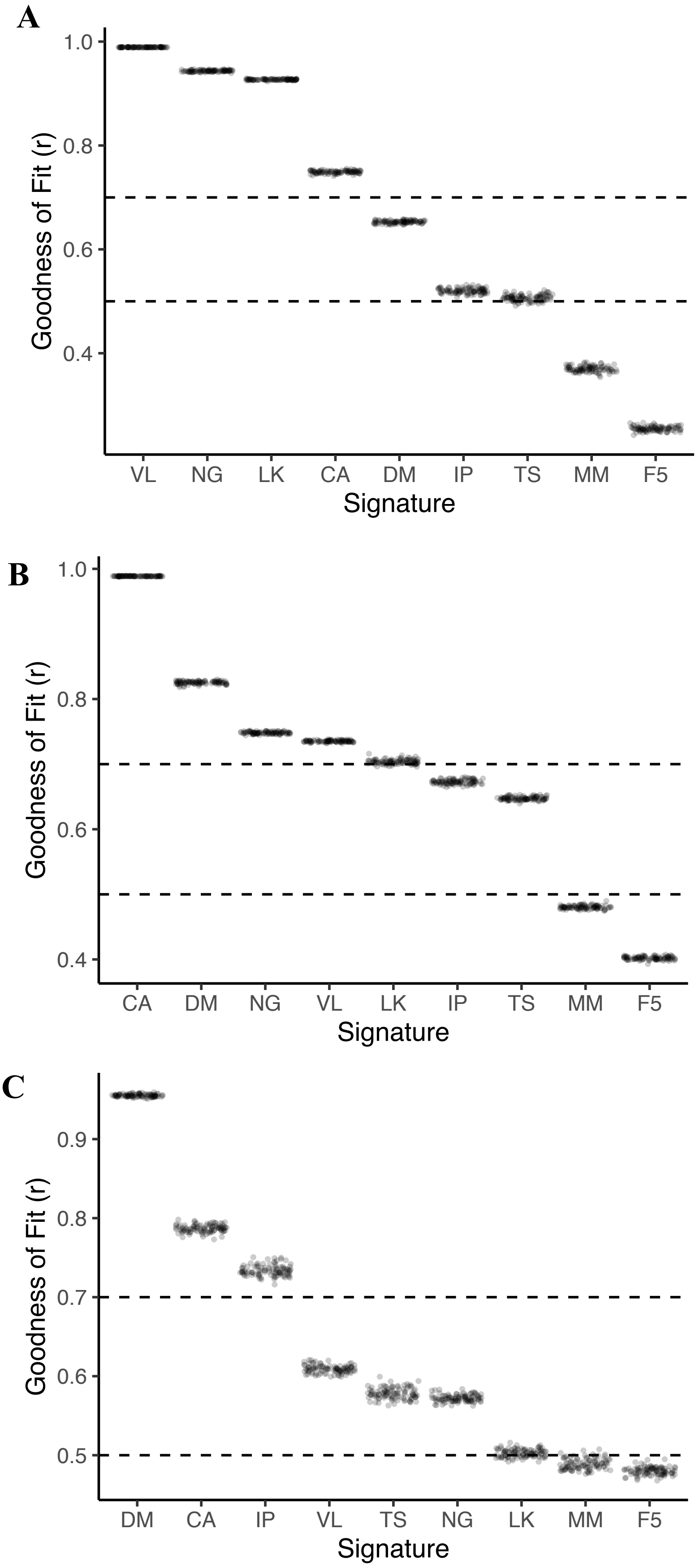


**Supplementary Figure 36. Goodness of fit across signatures in simulated data.** Each point represents one of the hundred mixtures per simulated dataset. **A.** VL-based simulation. **B.** CA-based simulation. **C.** DM-based simulation. Note that the order of signatures along the x-axis is based on median goodness-of-fit in that panel, and therefore differs between panels.

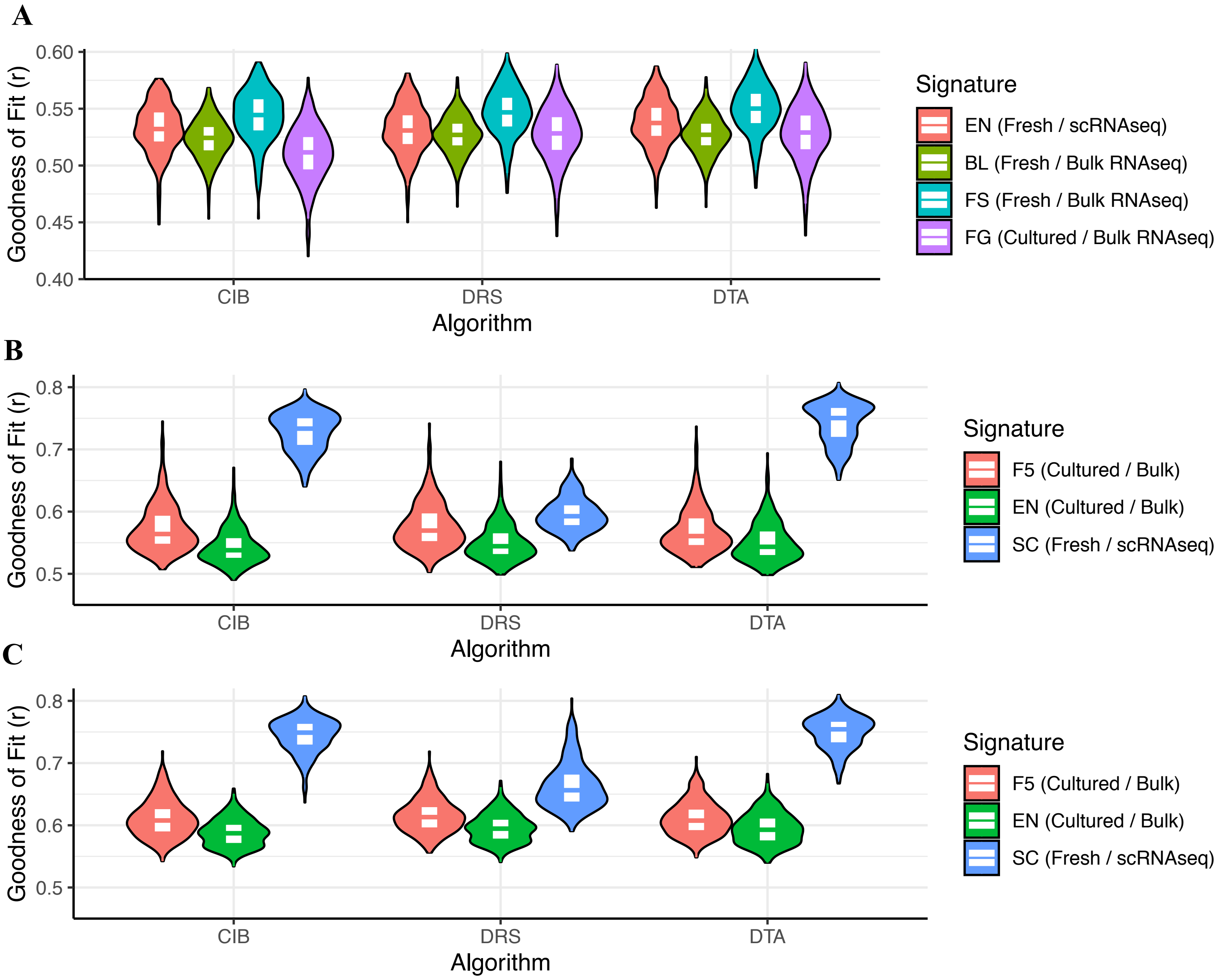


**Supplementary Figure 37. Violin plots of goodness-of-fit in two non-brain tissues in the GTEx dataset**. **A.** Pancreas samples. **B.** Heart left ventricle samples. **C.** Heart atrial appendage samples. *Fresh:* signature derived from freshly-processed human tissue. *Cultured:* signature derived from cultured cells. The bottom, middle, and top of the white boxes mark the first, second, and third quantiles, respectively.

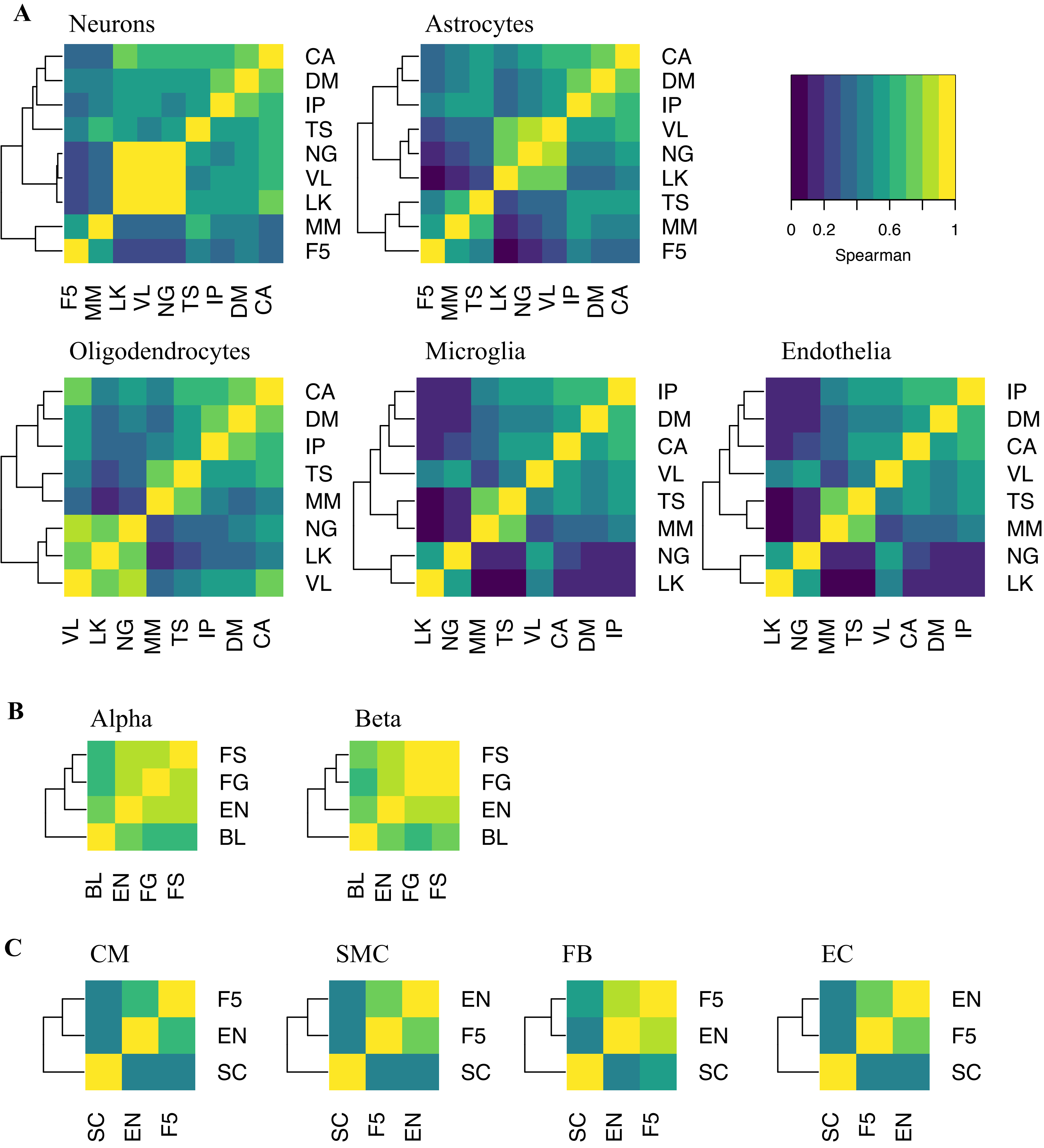


**Supplementary Figure 38. Heatmaps of Spearman correlations across signatures**. **A.** Across nine brain signatures. *Top left*: Neurons. *Top middle*: Astrocytes. *Top right*: legend. *Bottom left*: Oligodendrocytes. *Bottom middle*: Microglia. *Bottom right*: Endothelia. **B.** Across four pancreas signatures. *Left*: alpha cells. R*ight*: beta cells. Legend is per the top right panel of A. **C.** Across three heart signatures. *From left to right*: cardiomyocytes, smooth muscle cells, fibroblasts, and endothelial cells. Legend is per the top right panel of A.

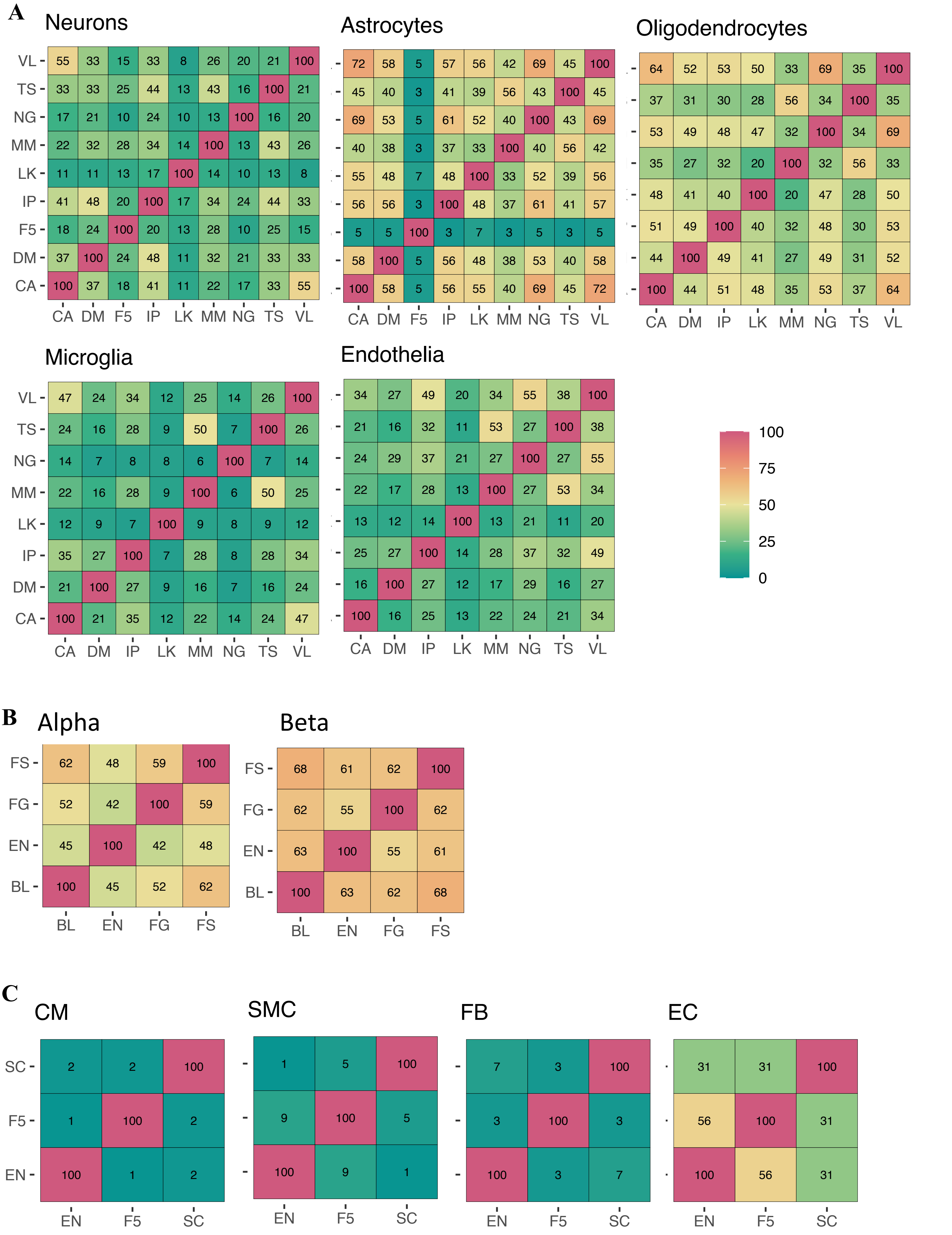


**Supplementary Figure 39. Heatmaps of intersection between the top 100 cell-type marker genes across signatures**. **A.** Across nine brain signatures. **B.** Across four pancreas signatures. Legend is per the bottom right panel of A. **C.** Across three heart signatures. *CM*: cardiomyocytes. *SMC*: smooth muscle cells. *FB*: fibroblasts. *EC*: endothelial cells. Legend is per the bottom right panel of A.

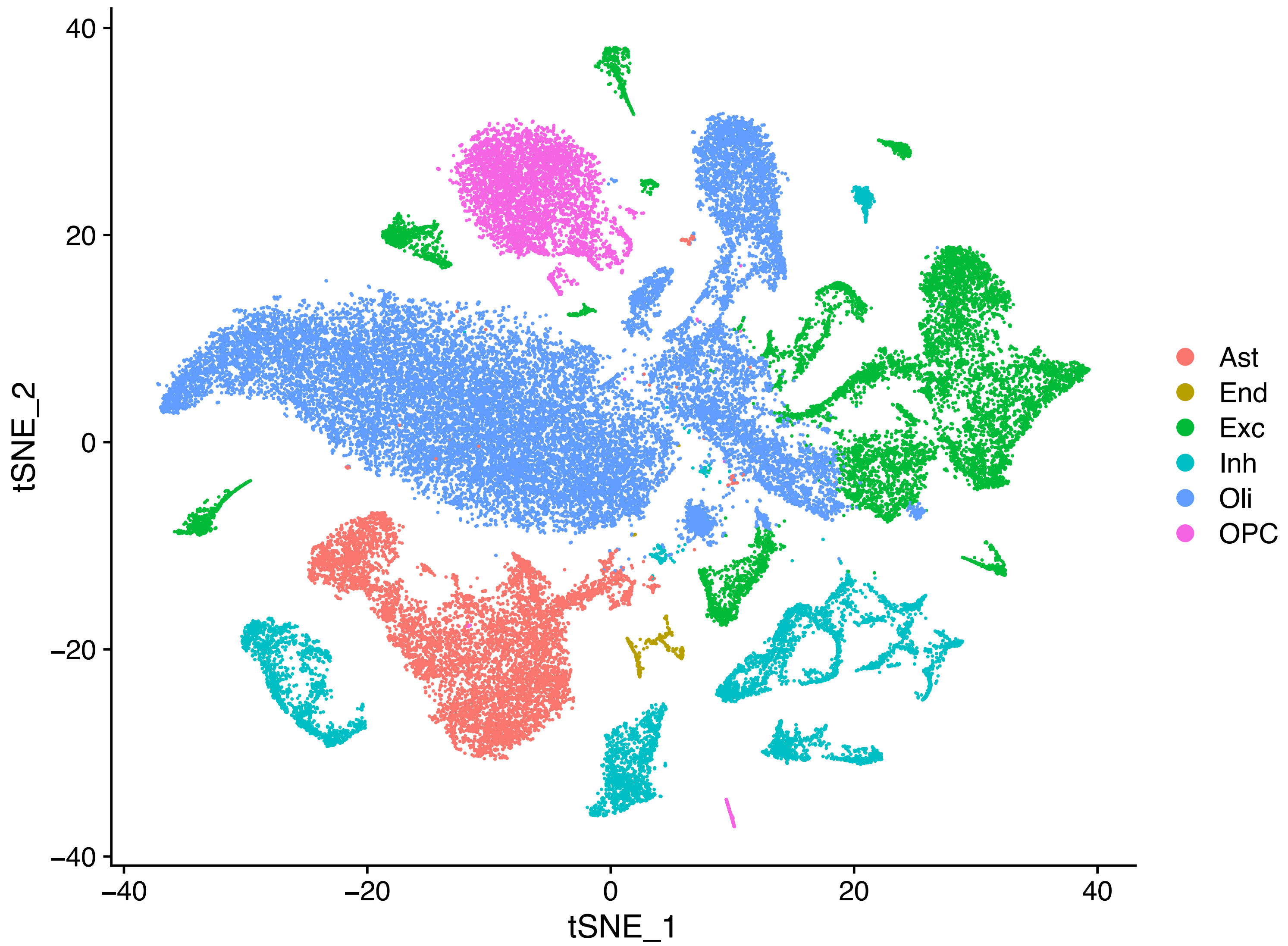


**Supplementary Figure 40. tSNE dimensionality reduction plot of snRNA-seq data generated as part of the present study**. Nuclei were annotated using the SingleR package to transfer labels from the NG signature. *Ast*: astrocytes. *End*: endothelia. *Exc*: excitatory neurons. *Inh*: inhibitory neurons. *Oli*: oligodendrocytes. *OPC*: oligodendrocyte precursor cells.

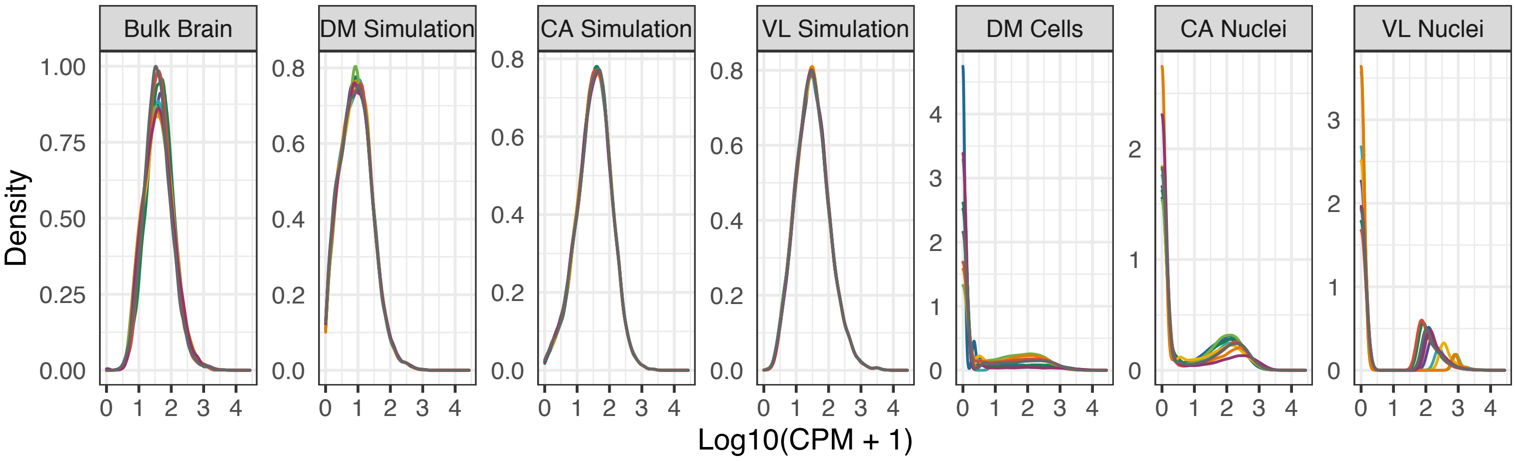


**Supplementary Figure 41. Distribution of gene expression values in real and simulated brain mixtures.** *Brain*: bulk brain RNA-seq from Parikshak et al. (2016). *Simulations*: *in silico* mixtures simulated from the corresponding dataset. *DM*: scRNA-seq from Darmanis *et al.*. *CA*: snRNA-seq data from the Human Cell Atlas. *VL*: snRNA-seq data from Velmeshev *et al.* (2019). Simulations contained 500 cells/mixtures for CA and VL, and 100 cells/mixture for DM. *Nuclei, Cells*: single-nuclei and single-cells from the corresponding dataset. Ten samples were randomly for each plot to minimise overplotting. Note: DM simulation is in units of RPKM rather than CPM.

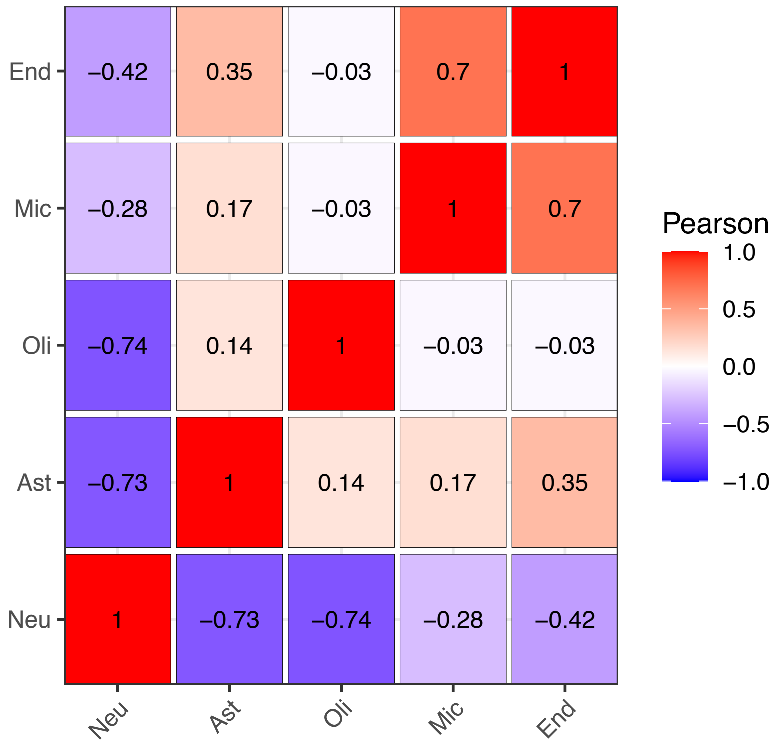


**Supplementary Figure 42. Heatmap of Pearson correlations for cell-ty[e proportion in samples used for ASD analyses**. Proportions were estimated using CIBERSORT and the Multibrain signature.

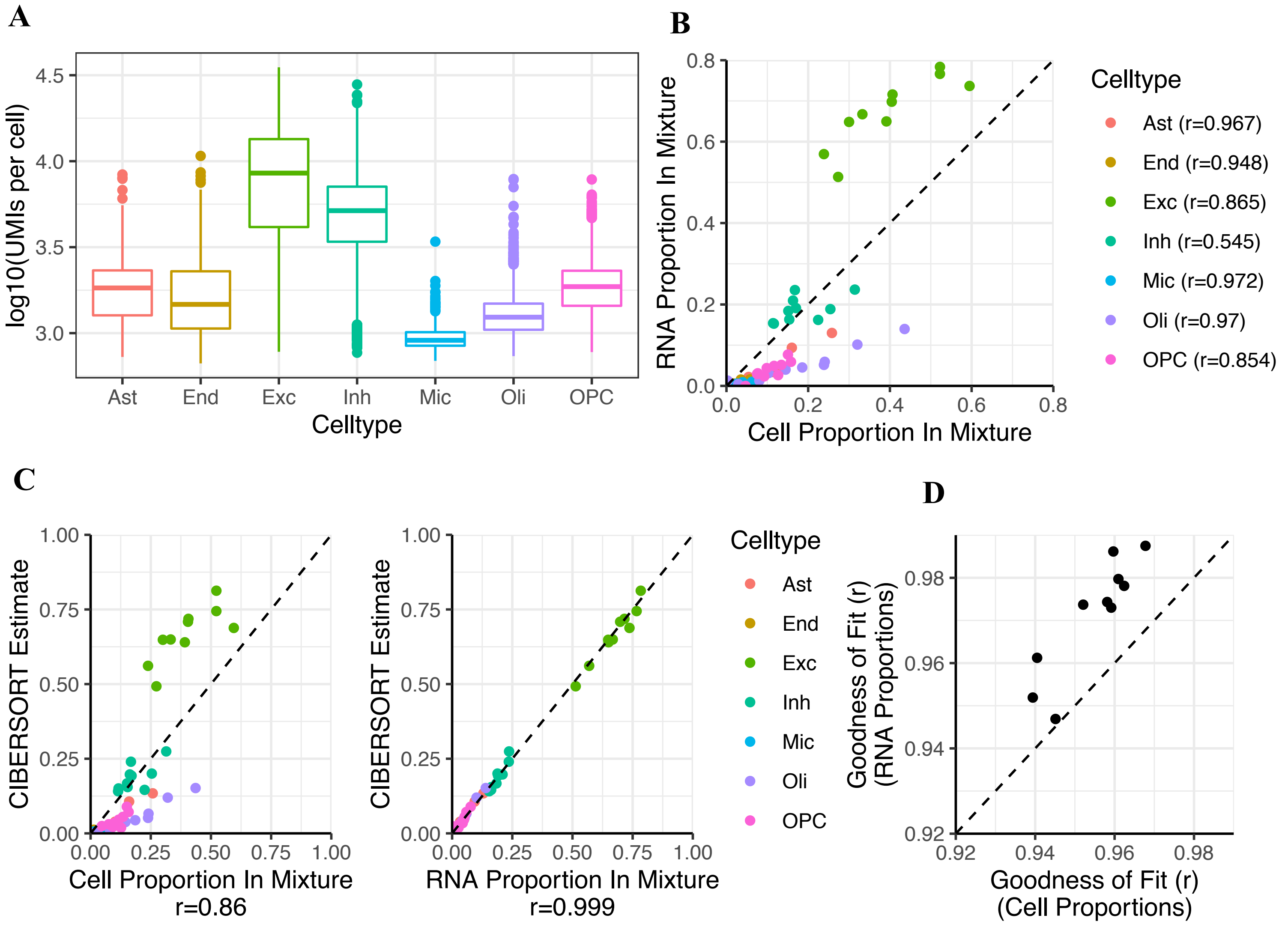


**Supplementary Figure 43. The relationship between deconvolution estimates, RNA proportions (pRNA) and cell-type proportion (pCt). A.** Boxplots of RNA content per cell, reflected in the number of unique molecular identifiers (UMIs) per-cell across cell types in the VL dataset. *Ast:* astrocytes. *End:* Endothelia. *Exc:* Excitatory Neurons. Inh: Inhibitory Neurons. *Mic*: Microglia. *Oli*: Oligodendrocytes. *OPC*: Oligodendrocyte Precursor Cells. **B.** Scatterplot of pCt vs. pRNA in pseudo-bulk samples from 10 individuals. **C.** Scatterplot of Estimated proportion (y-axis) versus true pCt (left) or pRNA (right)**. D.** Scatterplot of goodness-of-fit when reconstructing gene expression using pCt or pRNA. *Dotted black lines:* y=x

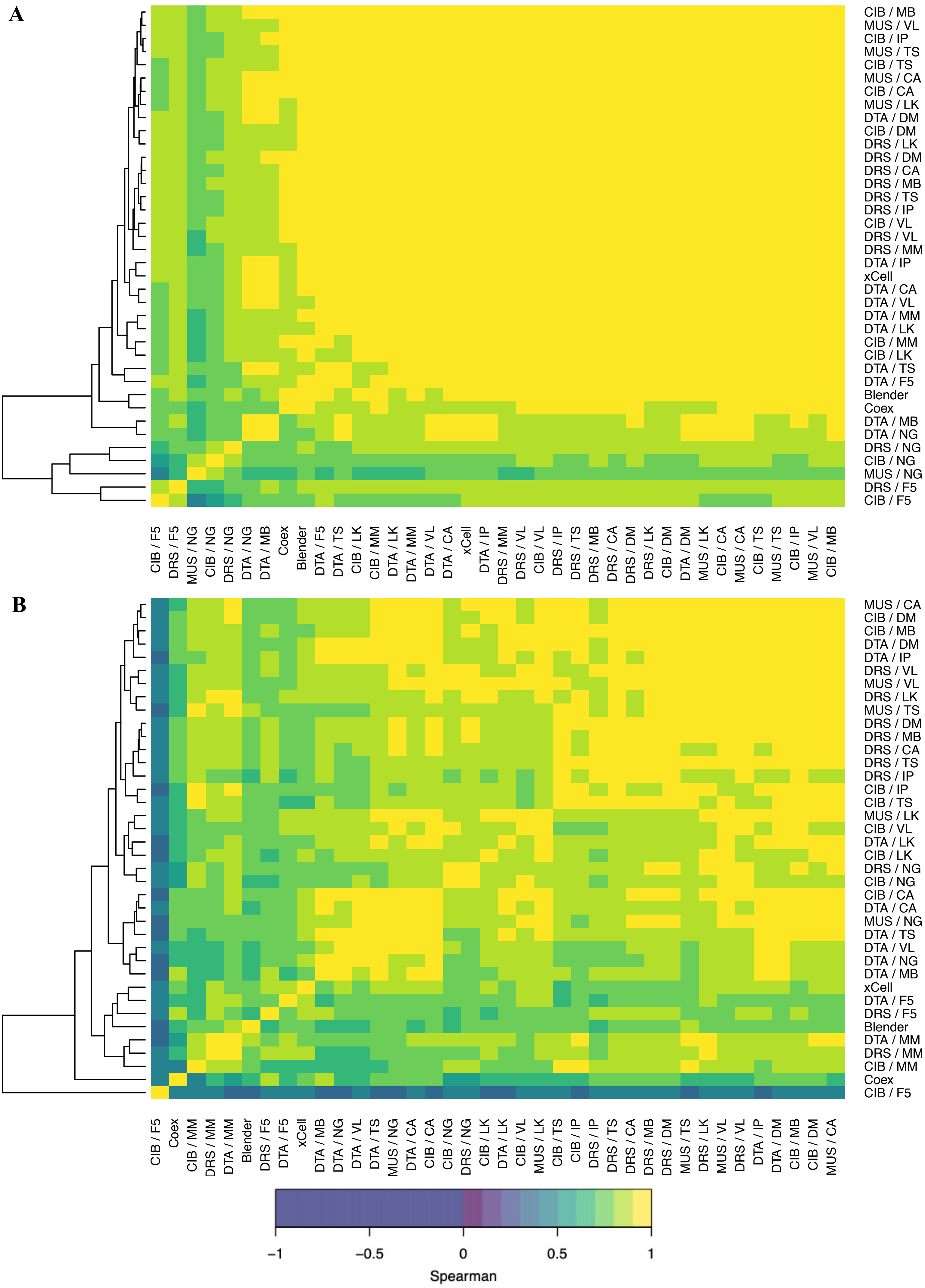


**Supplementary Figure 44. Heatmap of correlations in neuronal estimates across signatures and algorithms in bulk brain datasets. A.** GTEx. **B.** Parikshak *et al.*. Black squares represent NA, where the cell-type estimate had a variance of 0 (typically all estimates being all 0 or all 1). *Dotted black line:* y=0.5.

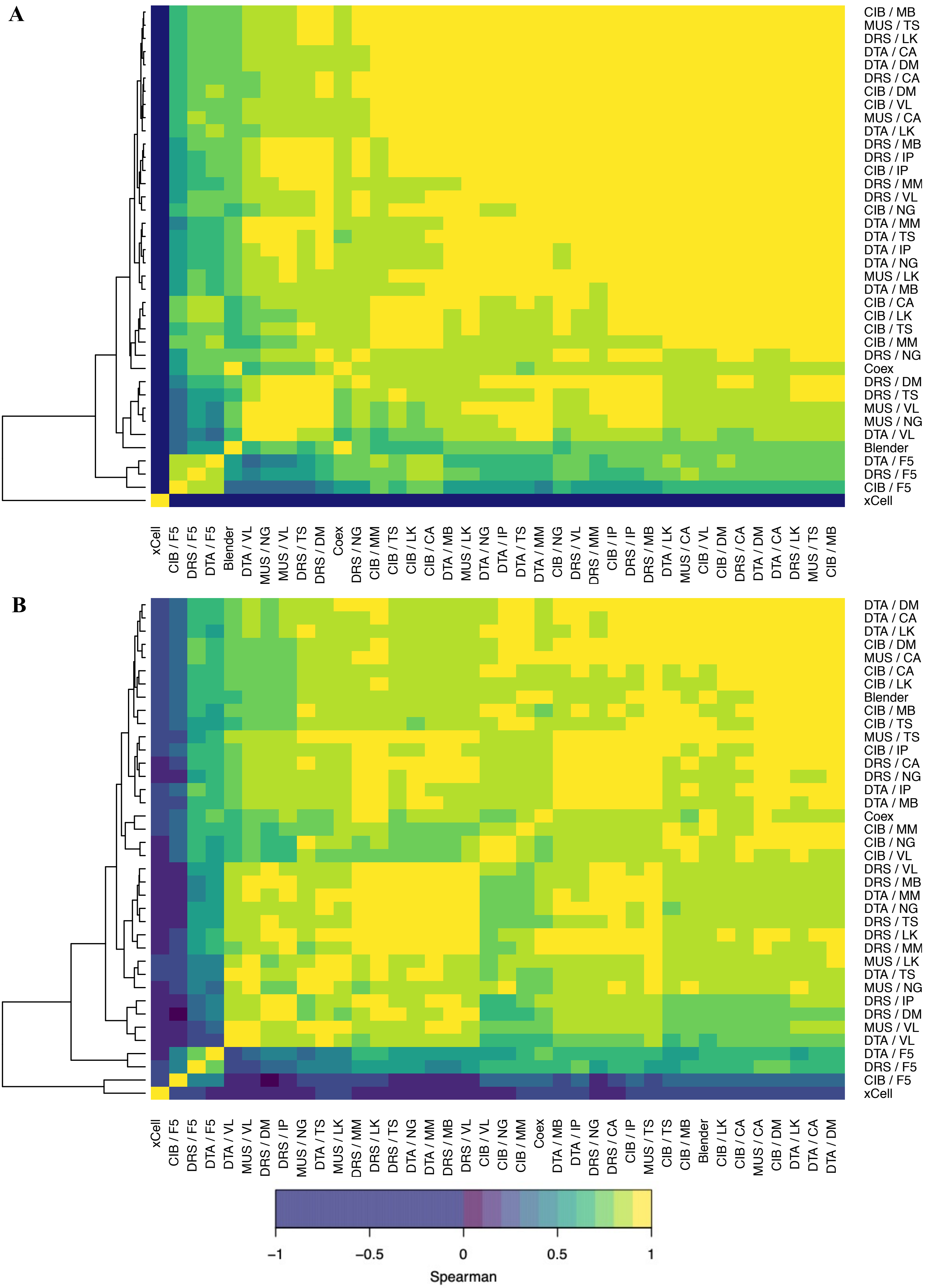


**Supplementary Figure 45. Heatmap of correlations in astrocyte estimates across signatures and algorithms in bulk brain datasets. A.** GTEx. **B.** Parikshak *et al.*. Black squares represent NA, where the cell-type estimate had a variance of 0 (typically all estimates being all 0 or all 1). *Dotted black line:* y=0.5.

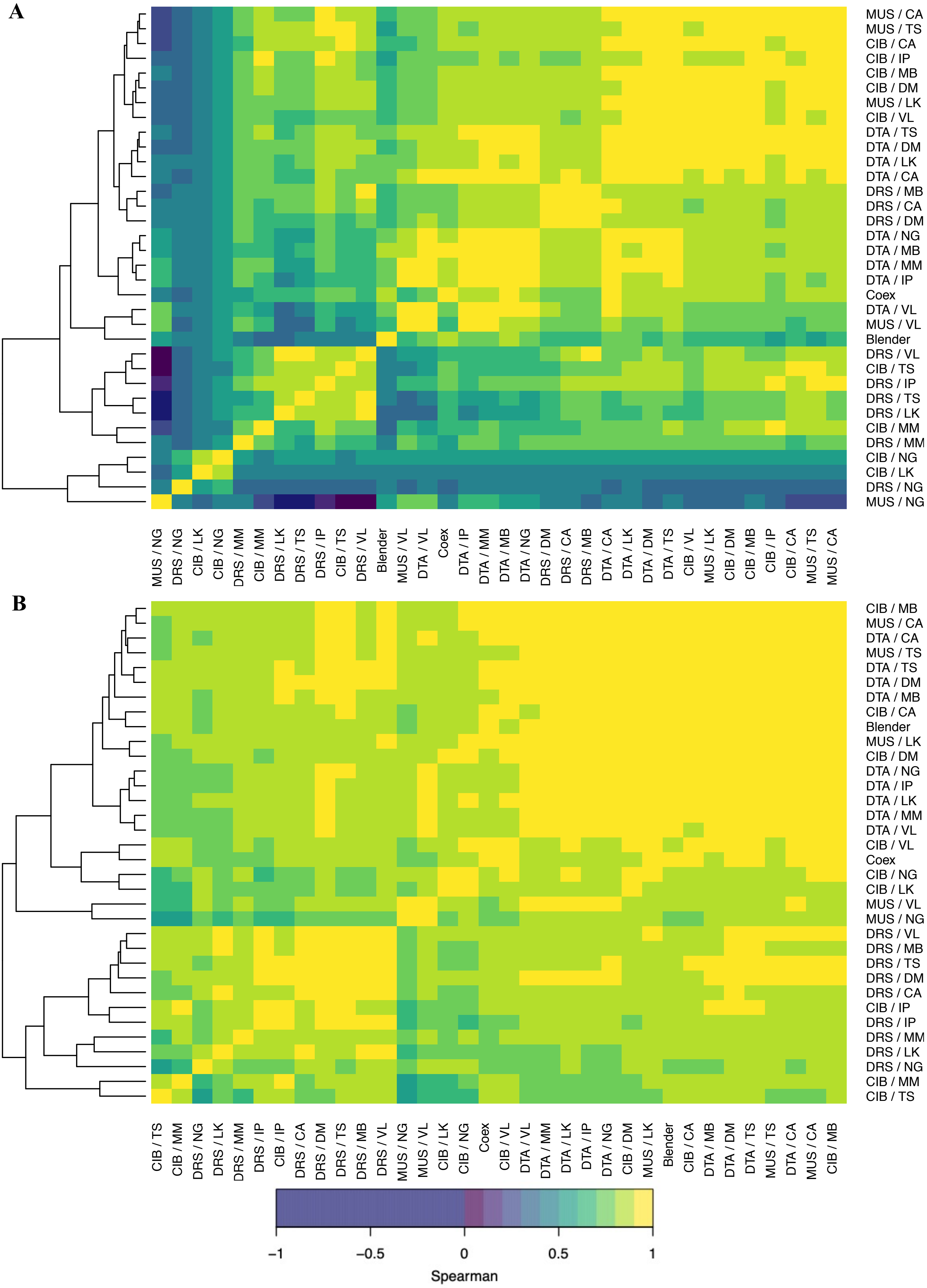


**Supplementary Figure 46. Heatmap of correlations in oligodendrocyte estimates across signatures and algorithms in bulk brain datasets. A.** GTEx. **B.** Parikshak *et al.*. Black squares represent NA, where the cell-type estimate had a variance of 0 (typically all estimates being all 0 or all 1). *Dotted black line:* y=0.5.

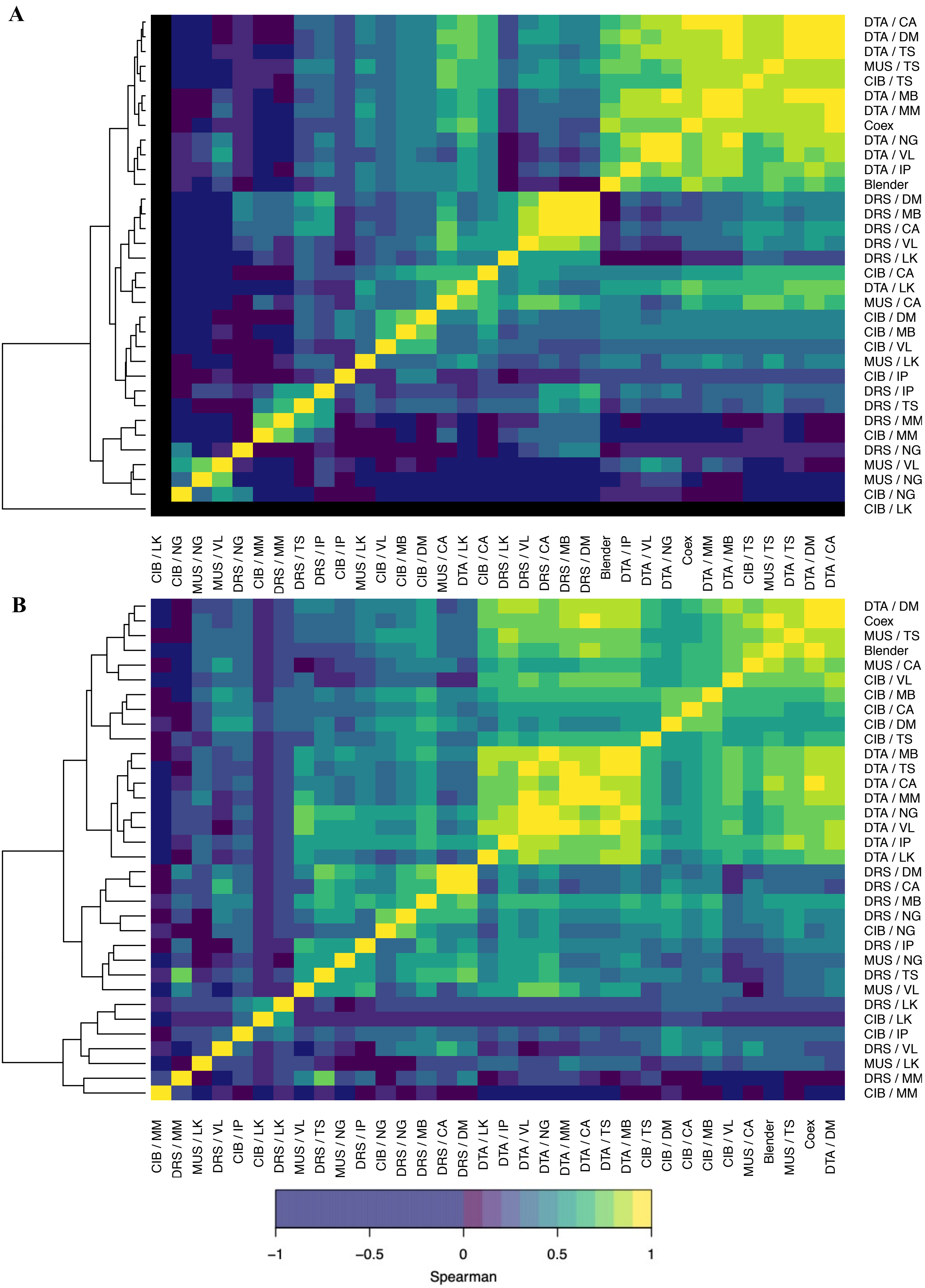


**Supplementary Figure 47. Heatmap of correlations in microglial estimates across signatures and algorithms in bulk brain datasets. A.** GTEx. **B.** Parikshak *et al.*. Black squares represent NA, where the cell-type estimate had a variance of 0 (typically all estimates being all 0 or all 1). *Dotted black line:* y=0.5.

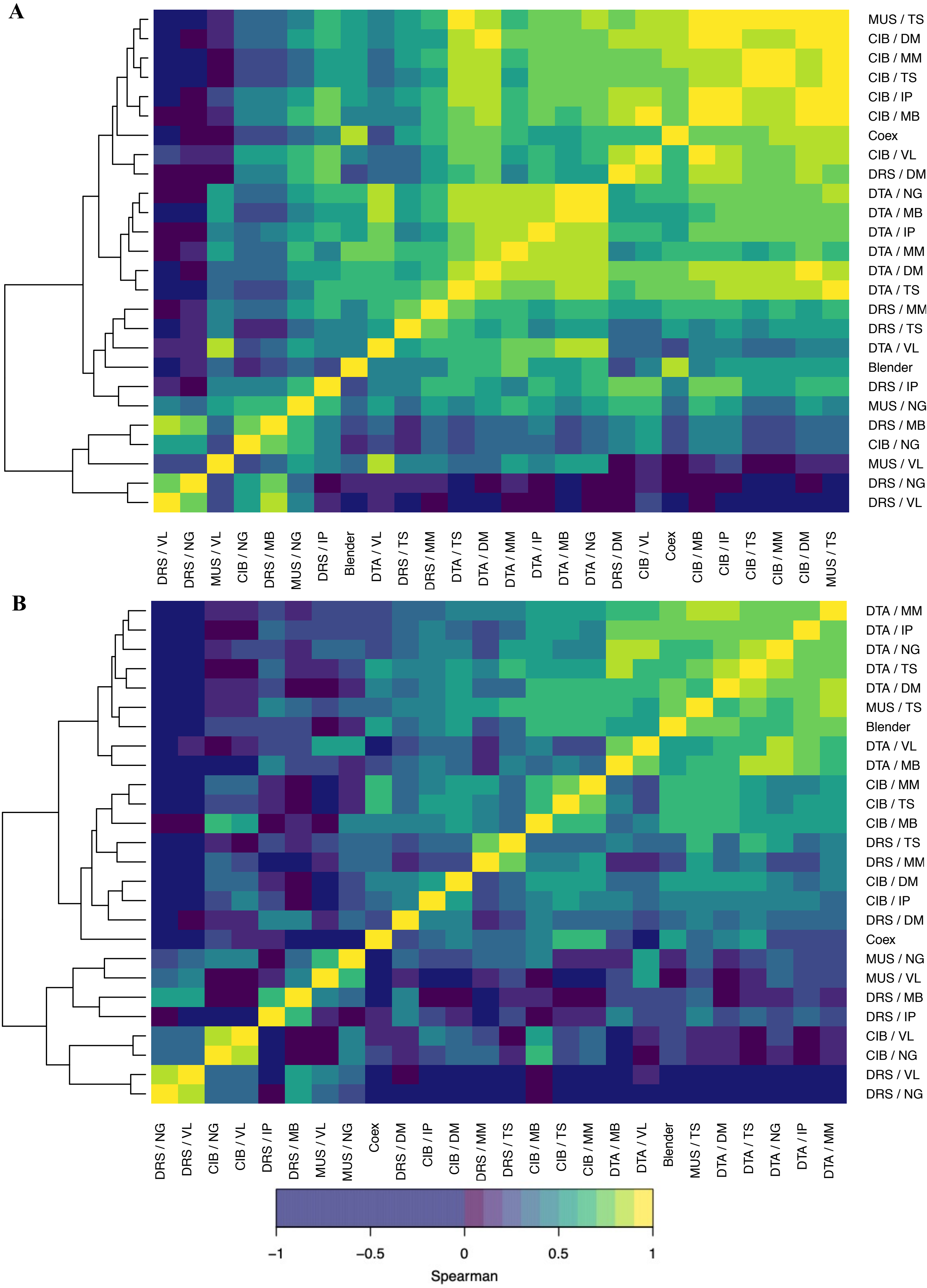


**Supplementary Figure 48. Heatmap of correlations in endothelial estimates across signatures and algorithms in bulk brain datasets. A.** GTEx. **B.** Parikshak *et al.*. Black squares represent NA, where the cell-type estimate had a variance of 0 (typically all estimates being all 0 or all 1). *Dotted black line:* y=0.5.

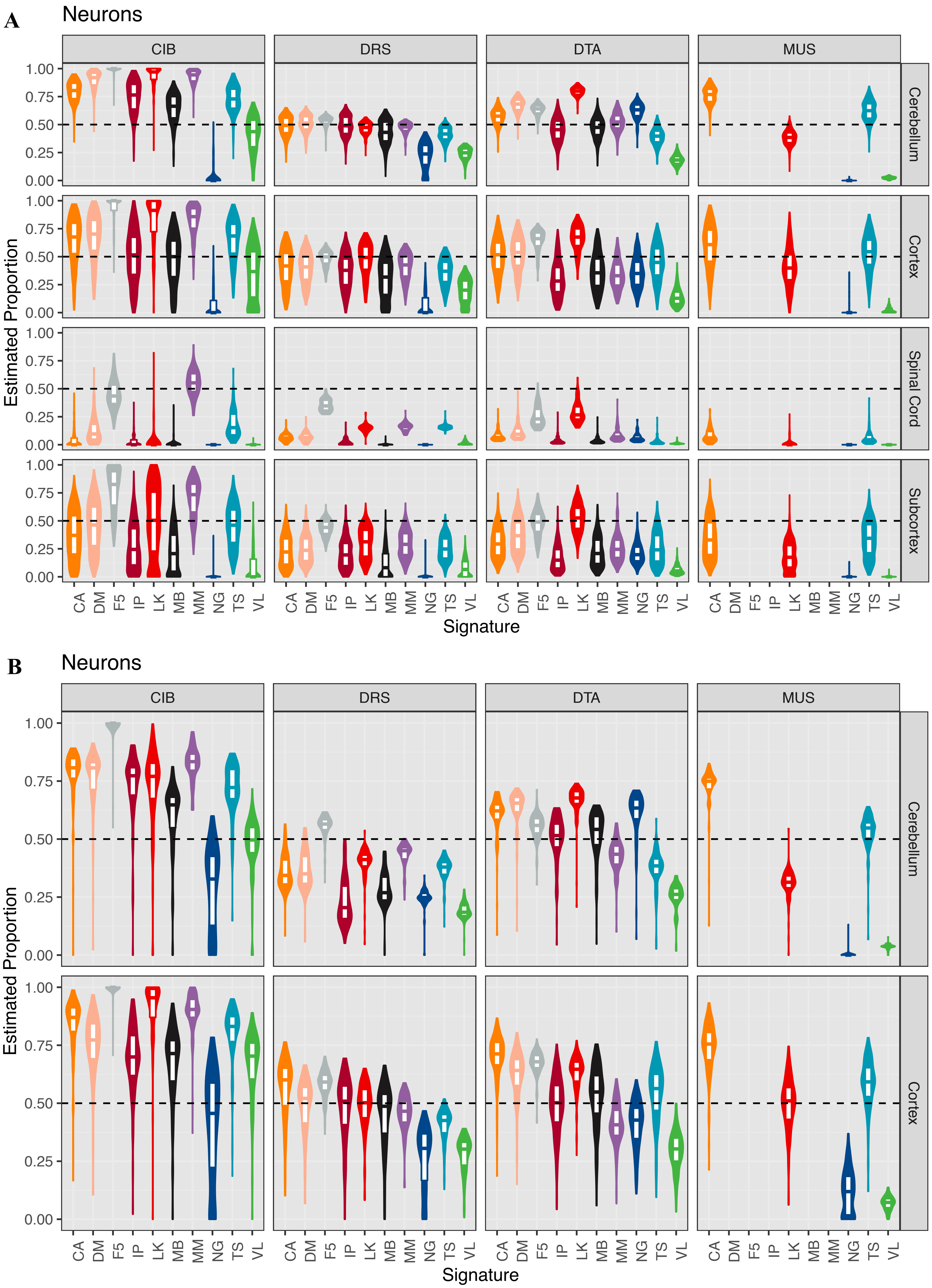


**Supplementary Figure 49. Distribution of neuronal deconvolutions estimates in bulk brain datasets. A.** GTEx. **B.** Parikshak *et al.*. The top, middle, and bottom of the white internal boxes mark the 75th, 50th, and 25th percentiles, respectively. *Dotted black line:* y=0.5.

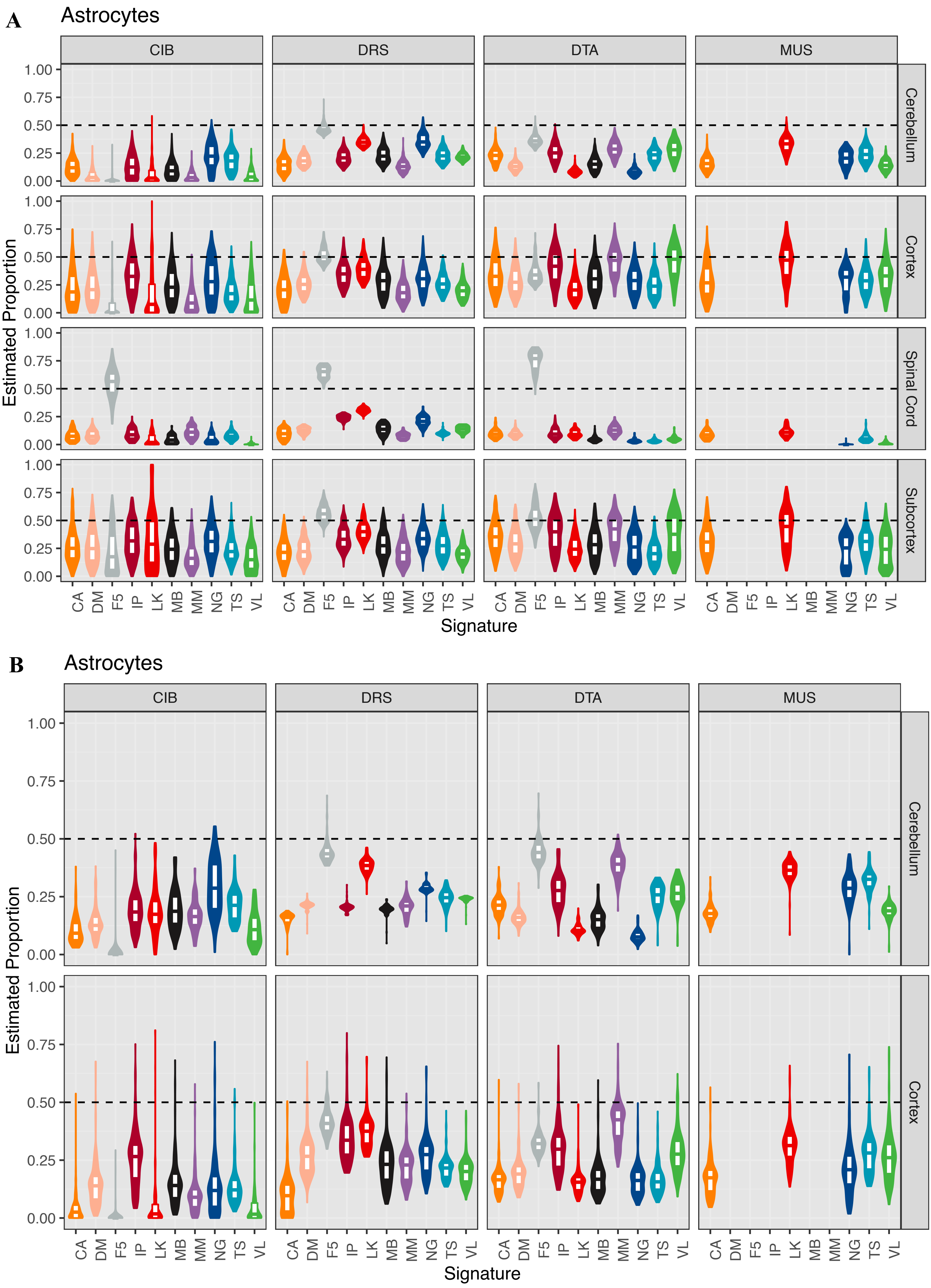


**Supplementary Figure 50. Distribution of astrocyte deconvolutions estimates in bulk brain datasets. A.** GTEx. **B.** Parikshak *et al.*. The top, middle, and bottom of the white internal boxes mark the 75th, 50th, and 25th percentiles, respectively. *Dotted black line:* y=0.5.

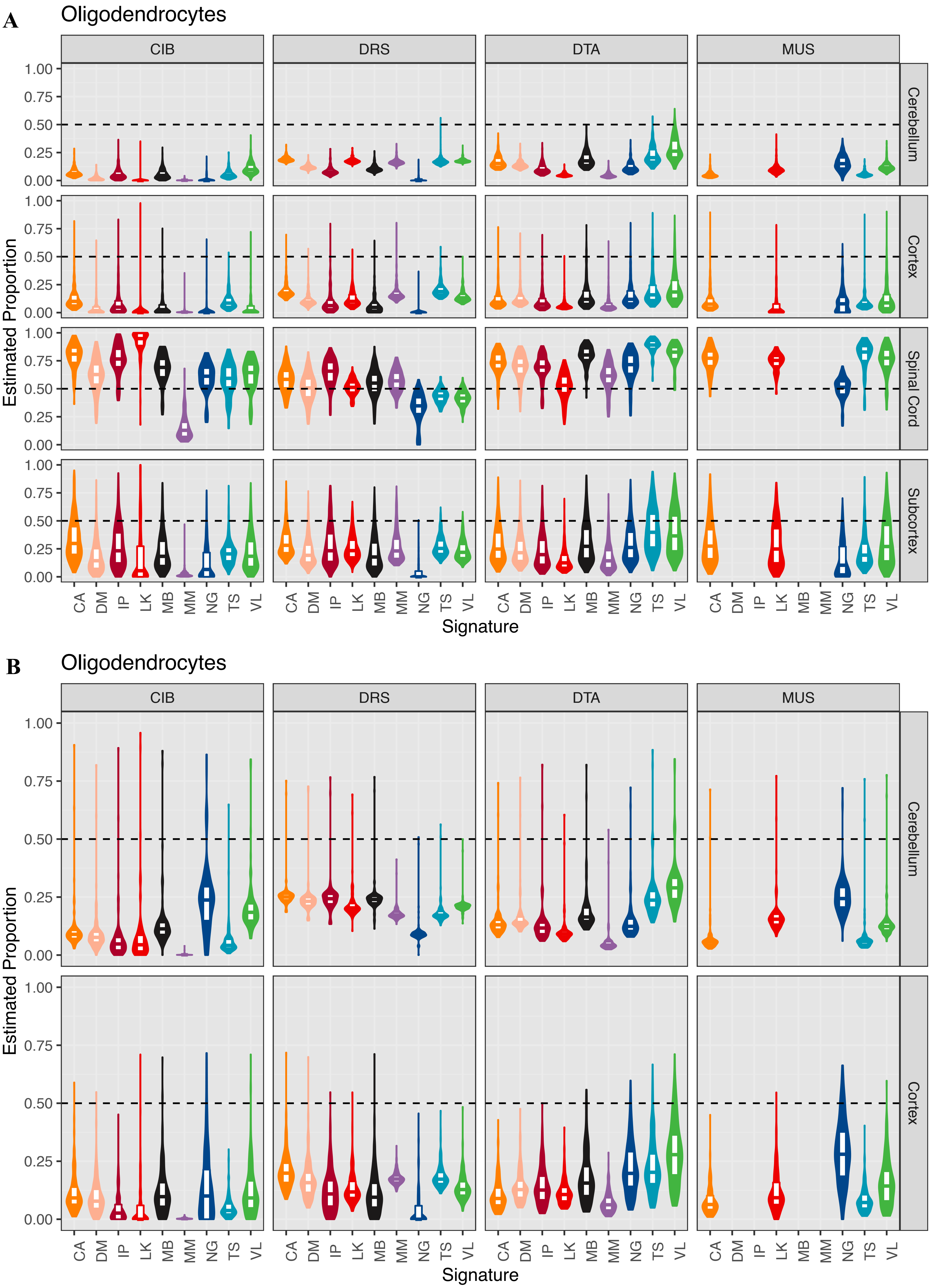


**Supplementary Figure 51. Distribution of oligodendrocyte deconvolutions estimates in bulk brain datasets. A.** GTEx. **B.** Parikshak *et al.*. The top, middle, and bottom of the white internal boxes mark the 75th, 50th, and 25th percentiles, respectively. *Dotted black line:* y=0.5.

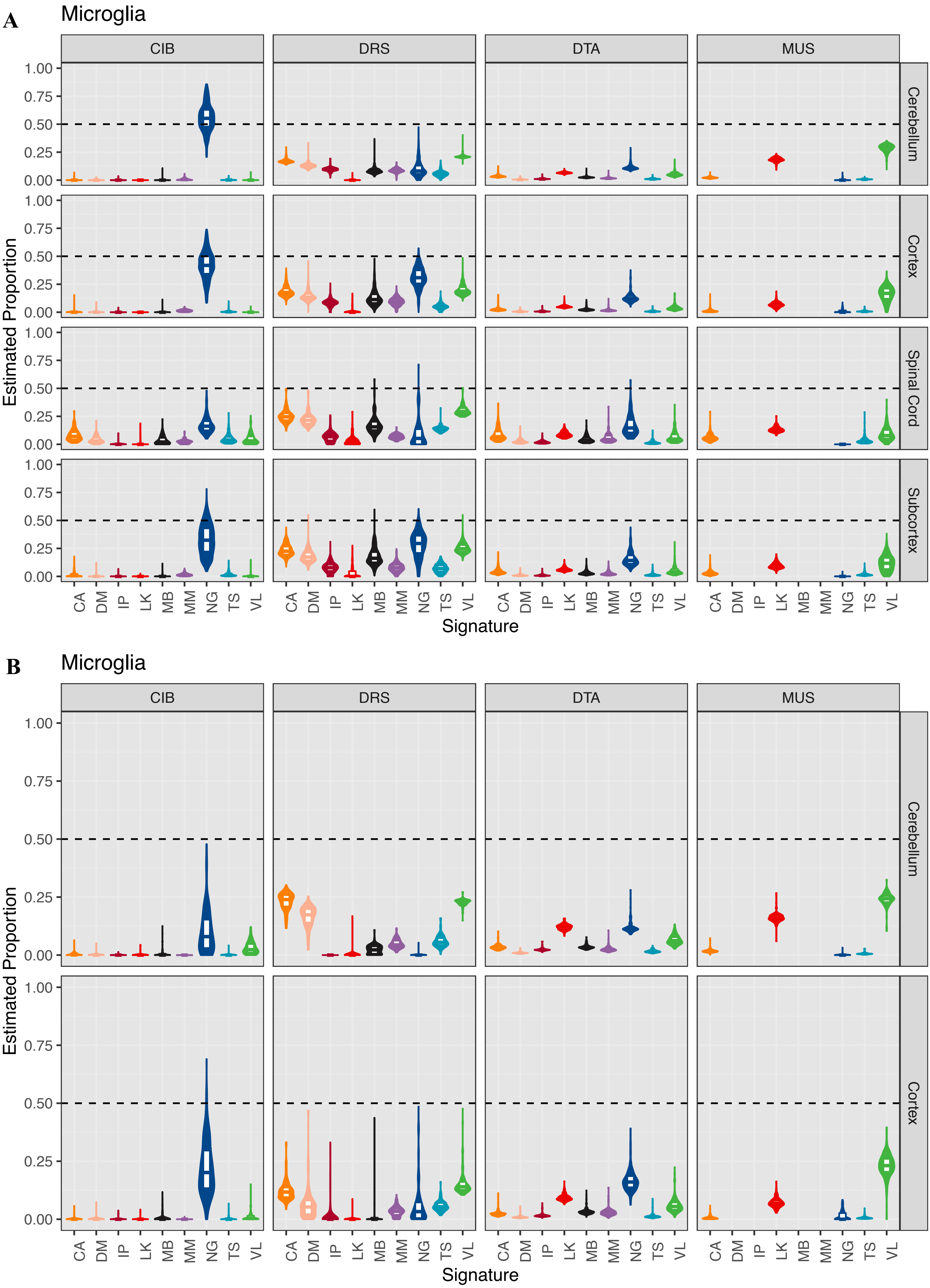


**Supplementary Figure 52. Distribution of microglial deconvolutions estimates in bulk brain datasets. A.** GTEx. **B.** Parikshak *et al.*. The top, middle, and bottom of the white internal boxes mark the 75th, 50th, and 25th percentiles, respectively. *Dotted black line:* y=0.5.

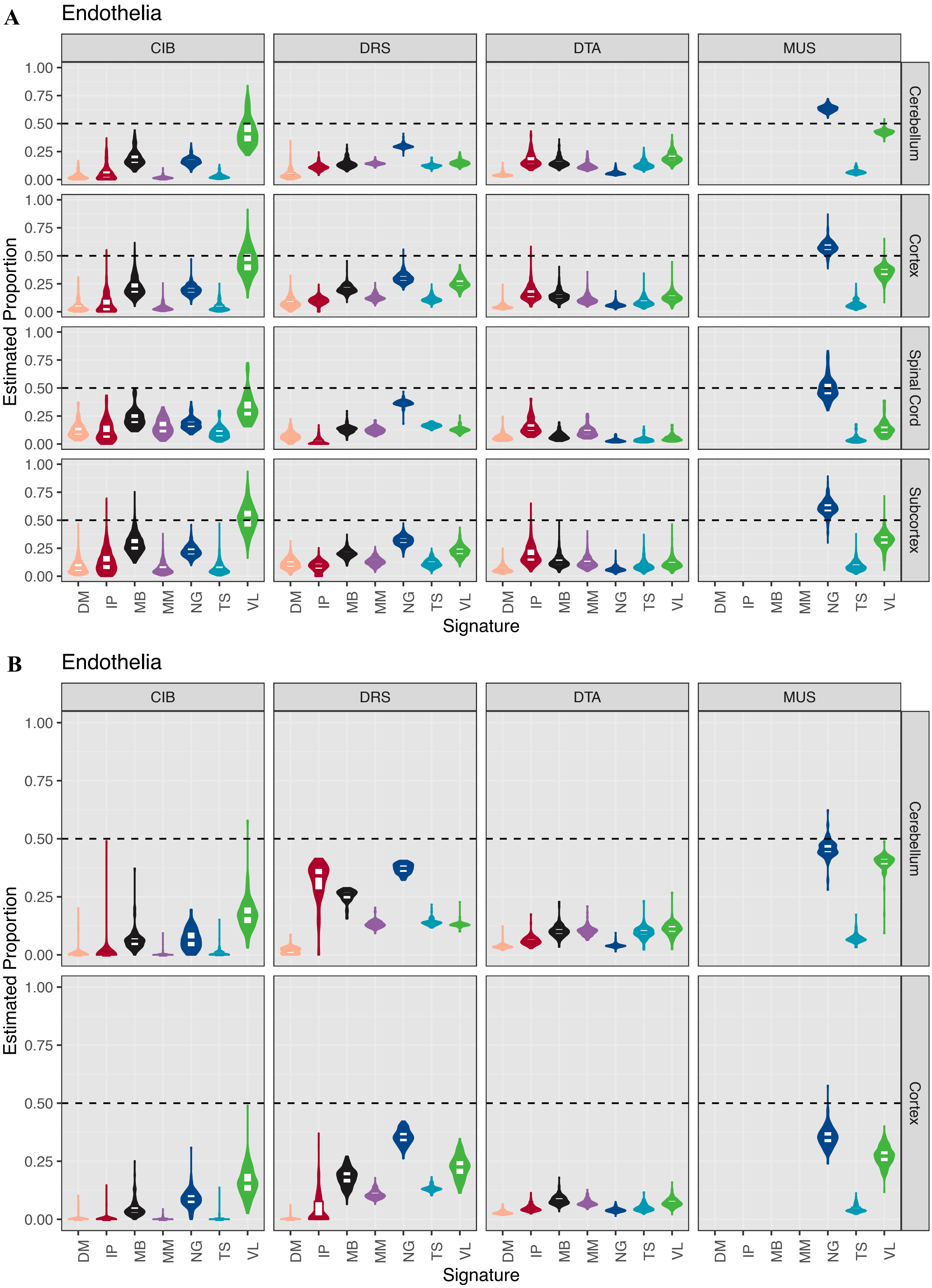


**Supplementary Figure 53. Distribution of endothelial deconvolutions estimates in bulk brain datasets. A.** GTEx. **B.** Parikshak *et al.*. The top, middle, and bottom of the white internal boxes mark the 75th, 50th, and 25th percentiles, respectively. *Dotted black line:* y=0.5.
